# Learning geometric models for developmental dynamics

**DOI:** 10.1101/2024.09.21.614191

**Authors:** Addison Howe, Madhav Mani

## Abstract

Successful development from a single cell to a complex, multicellular organism requires that cells differentiate in a coordinated and organized manner in response to a number of chemical morphogens. While the molecular underpinnings may be complex, the resulting phenomenon, in which a cell decides between one fate or another, is relatively simple. A body of work—rooted in dynamical systems theory—has formalized this notion of cellular decision making as flow in a Waddington-like landscape, in which cells evolve according to gradient-like dynamics within a potential that changes shape in response to a number of signals. We present a framework leveraging neural networks as universal function approximators to infer such a parameterized landscape from gene expression data. Inspired by the success of physics-informed machine learning in data-limited contexts, we enforce principled constraints motivated not by physical laws, but by this phenomenological understanding of differentiation. Our data-driven approach infers a governing landscape atop a manifold situated within expression space, thereby describing the dynamics of interest in a biologically meaningful context. The resulting system provides an intuitive, visualizable, and interpretable model of cellular differentiation dynamics.

## Introduction

Over the course of mammalian development, a single, totipotent cell gives rise to the trillions of cells that make up the mature organism. From this initial zygote, a vast array of functionally distinct cell types emerges at particular stages of development, in a self-organized, robust, and reproducible manner [1]. Cellular differentiation—the process by which a progenitor cell develops into one of several more specialized cell types—is a phenomenon essential to development, from embryogenesis to the continuous production of blood cells. Modern technologies allow us to study these processes at the level of individual cells. Single-cell RNA sequencing (scRNAseq) provides a snapshot of the transcriptome by measuring the abundance of mRNA transcripts for every gene in a cell [2]. A typical transcriptomic dataset may encompass thousands of cells and genes. Even more targeted approaches that quantify a limited number of gene products, like flow cytometry, yield datasets complex enough to require advanced analytical techniques [3, 4]. Significant progress has been made in discerning the different cell types present in such high-dimensional data, providing insight into the process of cellular differentiation across various contexts [5–9]. The identification of new cell types and their hierarchy in a developmental tree is often a celebrated achievement of sequencing studies, and treated as near-tantamount to a functional understanding of the phenomenon. However, cataloging cell types is only the beginning, and it is less obvious how one begins to make sense of the dynamical information captured in such experiments. That is, while we might be able to confidently identify the cell types into which a progenitor cell may differentiate, an understanding of the route through gene space that a cell takes in the process of making that decision, and the mechanisms by which such a trajectory is biologically controlled, remain elusive. Understanding the intricacies of this path, and the role of chemical signals that shape it, is at least as valuable as knowing the cell types appearing along its course. Deciphering these dynamics, and the ways in which they depend on signaling, may facilitate the design and control of patterns of cellular differentiation and development, with obvious applications to synthetic biology and medicine.

A number of approaches have been advanced to further our understanding of the dynamics of cellular differentiation. Classical models of gene regulatory networks describe the interplay of genes using a system of differential equations, and thus explicitly model high-dimensional dynamics [10, 11]. However, the true forms of these equations are generally unknown in the context of biology, and these models typically rely on a gross simplification of reality and suffer from a large number of parameters [12, 13]. In the context of scRNAseq, due the destructive nature of the technology cells can only be sequenced once, and therefore one cannot directly observe a cell’s trajectory through gene space. In light of this, traditional methods infer developmental trajectories along a pseudotemporal axis by organizing sampled cells according to a measure of transcriptomic similarity, thereby constructing a path through gene space along which cells may traverse [14–16]. As an extension, measures of RNA velocity estimate from sequencing data a transcriptomic derivative—the direction within gene space that a cell is changing—assuming a simple model of splicing [17, 18]. More recently, RNA velocity methods inspired by machine learning have allowed for more general modeling assumptions with respect to splicing and other regulatory dynamics [19, 20]. These methods have proved useful in improving trajectory inference and offering dynamical insight into cellular differentiation as flow through a vector field, often in a low-dimensional latent space [21–23]. They do not, however, provide a direct link from chemical signaling to cell fate decision making. That is, one obtains a vector field representation of cellular trajectories that is a snapshot of the global dynamics as they played out and were captured in the data. The question remains as to how the governing signals driving the differentiation process would alter that vector field, were they to be perturbed over the course of development.

An alternative approach casts the problem in geometric terms [24], and models cellular decision making as flow in a landscape—an idea dating back to Waddington, who introduced the “epigenetic landscape” as a metaphor for development [25]. This metaphor envisions a cell as a ball in a hilly terrain, which rolls downhill through valleys corresponding to particular cell states (Fig. 1a). Along the way, a valley may split into diverging paths, representing a point at which a cell chooses between alternate states. Eventually, the cell comes to reside in a particular valley, or basin, corresponding to its terminal fate. The tendency of the rolling ball to remain in a valley when slightly perturbed captures the biological notion of canalization—the robustness of a developmental process to various environmental or genetic insults [26].

**Figure 1.**
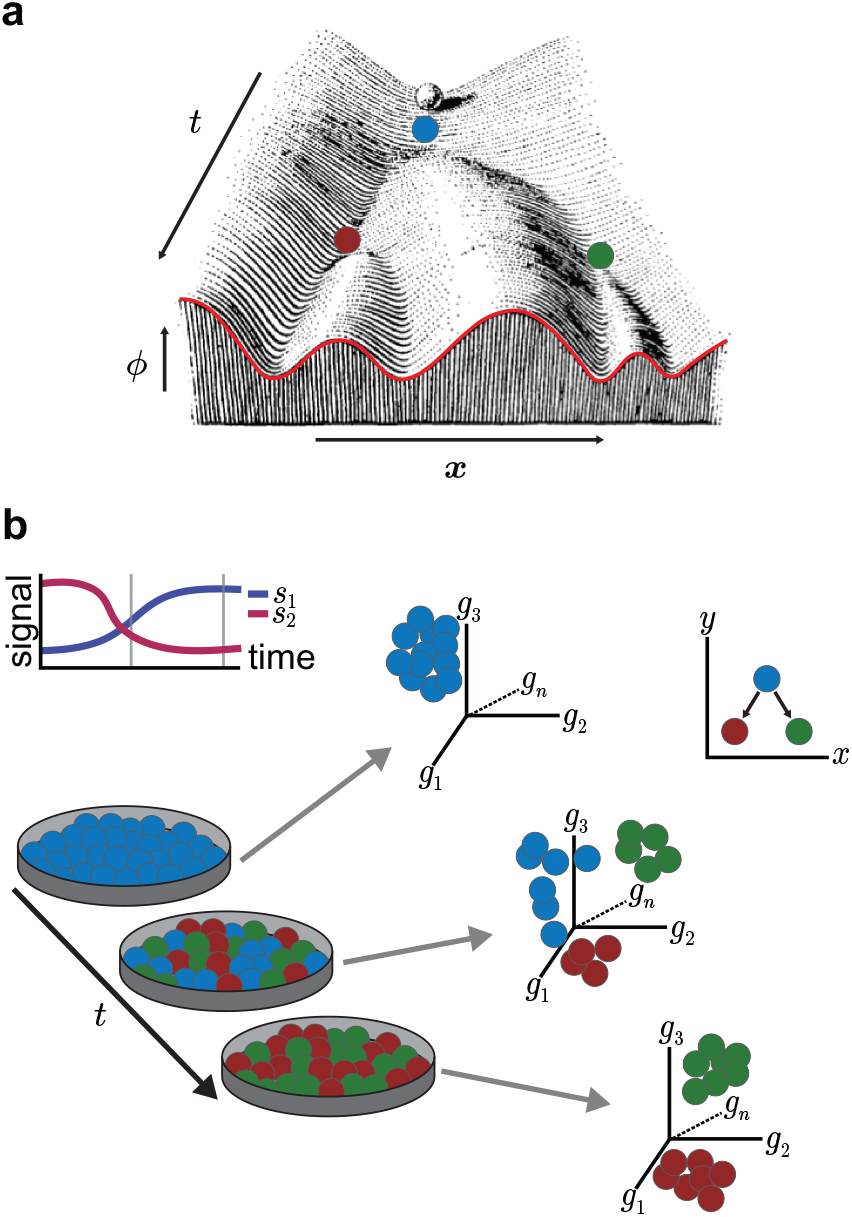
*In vitro* experiments capture cellular differentiation and quantify the cell state transitions depicted in Waddington’s metaphor. **(a)** Waddington’s original illustration of the epigenetic landscape [25], with annotations added. The horizontal axis (***x***) encapsulates the state of a cell, while depth (*t*) corresponds to a notion of developmental time. Waddington’s illustration can be interpreted as a one-dimensional potential varying either across a temporal axis or with respect to a control parameter (i.e. signal). **(b)** In *in vitro* experiments, a specified signaling timecourse induces an ensemble of pluripotent cells to transition into one of two possible states. Cells profiled at discrete timepoints capture the cell state transitions taking place in a high-dimensional gene space. At the same time, the phenomenon suggests a simple decision structure, which occurs in a lower-dimensional (*x, y*) space in which each signal serves to bias the movement of cells in a particular direction.

That these biological notions can be made mathematically rigorous using dynamical systems theory has long been appreciated [27, 28]. Various works have modeled cell fates as attractors in gene expression space and have used dynamical systems theory to explain and predict patterns of differentiation [29–34]. In recent years, a body of work has mathematically formalized a particular interpretation of Waddington’s landscape, in which signals parameterize a family of potential functions [35–39]. This work has been shown to be applicable to developmental systems as a way to understand the most basic developmental decision, in which a cell chooses between two alternate fates [40]. However, such applications typically rely on a prior understanding of the developmental system in question. One must have some qualitative understanding of the possible cell state transitions, or decision structure, *a priori*, in order to choose a mathematical ansatz suitable for the system. This knowledge may be available for certain model systems, to which decades of research have brought some degree of understanding, but otherwise must be deduced from careful analysis of the phenomenon, on a case-by-case basis. Moreover, current approaches generally represent cells in an abstract, low-dimensional space that lacks a concrete interpretation with respect to the original, high-dimensional gene space.

Motivated by these limitations, we investigate an approach to inferring a parameterized landscape model using a machine learning architecture, in which a time-independent, underlying potential function is parameterized by a neural network, and “tilted” in response to a number of time-dependent external signals. This data-driven framework simultaneously infers the underlying landscape, inducing gradient dynamics, along with a mapping of signals to their effect on the landscape. The landscape is learned atop a low-dimensional, but biologically meaningful, phase space, which we refer to as a decision manifold. We restrict ourselves to inferring two-dimensional landscapes, which are intuitive and easily visualized. At a more fundamental level, our search for a two-dimensional landscape is based on an assumption that there are two signals controlling the system, or at least the particular decision of interest, and that these signals serve to bias the flow of cells in particular directions along a manifold within the ambient state space. In this way, the signals act as constant forces conjugate to directions in state space, and it is this conjugacy that in general endows the low-dimensional space with biological meaning. The separation between the underlying landscape and the effect of signals in altering that landscape provides a powerful interpretation of the system, in which the manifold is exactly the space in which cells are controllable via signaling dynamics.

Our approach is inspired by applications of physics-informed machine learning to material physics and fluid dynamics [41, 42], in which a clear distinction is made between the physical laws that one insists be made explicit as part of a model, and those that are inferred from data, and thus generalizable. This separation makes possible an interpretable model, where overparameterization is constrained to a particular component. Although that component may be opaque, its function as part of the larger modeling framework remains transparent. This contrasts with more modern ML architectures, such as transformers, which, while often effective, are entirely opaque and not easily interpretable. In the present case, the physical principle that we explicitly enforce—namely gradient flow—does not arise from a physical law governing development, but rather the phenomenological, mathematically formalized insights of Waddington.

We begin by providing background on the mathematical formalization of a Waddington-like landscape as a potential that induces gradient dynamics and changes in response to signals. Then, we describe our contribution: a neural network-based architecture designed to represent such a system. We show that this architecture can be used to infer essential features of an underlying landscape, given synthetic, biologically motivated data. Finally, we apply this model to a real dataset detailing cellular differentiation in the context of an *in vitro* system modeling the early mouse epiblast.

## Background

### Landscape models of cellular differentiation

In Waddington’s landscape, the axis capturing depth (labeled *t* in Fig. 1a) is often associated with a notion of developmental time, and is understood through our impression that gravity attracts the ball, thereby producing a directed, downward flow. In embryonic development, for instance, where there is a clear temporal direction, this interpretation is sensible, and one can treat time as a parameterization of depth in the landscape. Biologically speaking, it is now understood that within the embryo tissues synthesize and secrete signaling molecules, in a manner dependent on their state, that themselves elicit changes in those cells receiving the signal. As such, the interplay between the states of cells within a tissue and the signals they secrete and sense is what drives the self-organized and time-ordered dynamics of the nascent embryo. This coupling, along with technical limitations inherent in studying live development at the cellular level, imposes a significant roadblock to direct investigation of development *in vivo*. In contrast, *in vitro* studies that measure the response of cells to experimentally prescribed signaling timecourses attempt to decouple the dynamics of states and signals, in a controlled setting (Fig. 1b). In these experiments, one subjects a sample of cells to a specified signaling timecourse, and profiles the state of the ensemble (or a subset of it) periodically over the course of the experiment, via flow cytometry or scRNAseq. While such simplified experimental scenarios preclude a deeper understanding of the exact nature of the coupling between signals and states, they allow us to study in quantitative precision the map from signals to cell state changes.

In this setting, the parameter of interest is not time, but rather the set of governing signals. Depth in Waddington’s landscape can be viewed as an axis along which one or more control parameters vary, thereby changing the states available to a developing cell. These states are still represented by the basins in each cross section of the terrain. In Fig. 1a, each cross section is a one-dimensional curve, or potential. As the signal varies, the potential changes shape, and bifurcations occur in response to a change in signal. There are now two notions of downhill flow. Within the one-dimensional potential, cells move downhill in the horizontal direction, according to the gradient. The second notion of downhill flow is only realized given a prescription of a signaling timecourse. That is, a temporal function describing the signaling dynamics induces an extension of the one-dimensional potential along a temporal axis, reproducing Waddington’s illustration. A more general picture, however, entirely disregards a temporal parameterization, and depicts all possible realizations of the potential across the signaling, or control, axis.

Mathematically, this notion of a *parameterized landscape* can be expressed as a potential function *ϕ*(***x***; ***p***) defined over a phase space, ***x***⊆ ℝ^*d*^, and parameterized by a vector ***p*** ⊆ℝ^*r*^. As ***p*** changes, the landscape changes shape. The gradient of the potential induces a flow according to the vector field

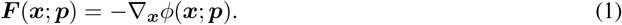

A priori, the vector ***p*** need not carry any biological significance, however, we will shortly associate its role of altering the shape of the landscape with a set of signals affecting the biological system, and it is appropriate to think of the parameters as functions of these signals.

Inspired by Waddington’s metaphor, we associate particular cell states to the landscape attractors, and interpret movement of cells from one attractor to another as a cell fate decision. This movement may be driven by stochastic fluctuations causing cells to escape one basin of attraction and to enter another. Alternatively, a change in the landscape, resulting from some change in the governing signals, might result in the disappearance of an occupied basin, causing cells to relocate as they are no longer in the vicinity of an attractor. In either case, the particular route that cells take as they flow through the landscape is defined by the stable and unstable manifolds of the saddles and attractors, with saddles serving as demarcations of a decision point [39]. Cells flow toward a saddle, then veer off in one direction or another, following a particular unstable manifold, to ultimately arrive at a new attractor.

Rand et al. [39] present parameterized landscapes as mathematical objects that derive from dynamical systems describing gene networks. They restrict their consideration to biologically relevant systems, namely those with a finite number of fixed points and no recurrent behavior, a class of dynamical systems captured by *gradient-like Morse–Smale systems* (see Supplementary Materials S1D for details). They then use the mathematical properties of these systems to enumerate the ways in which a 3-attractor landscape can undergo generic bifurcations, in response to two control parameters. Their geometric approach provides a small number of equivalence classes of dynamical systems, along with an algebraic normal form—a polynomial in two variables—for each. Each class is composed of systems that share a common, global bifurcation structure, and thus admit the same patterns of transitions between attractors.

Sáez et al. [40] utilize this classification to construct a parameterized landscape model of cellular differentiation in an *in vitro* experimental context relating to mouse embryonic stem cell differentiation. This application illustrates an effective use of geometric normal forms, which capture qualitative aspects of the dynamics, to infer quantitative features relating to the role of signaling in a biological system. The normal form is chosen in advance, and what is inferred is the role that signals play in altering that prespecified landscape. However, the use of an abstract normal form is limiting in that there is no direct connection between the landscape coordinates (i.e. the variables *x* and *y* appearing in the normal form) and the molecular composition of cells, the original space in which they are observed. Rather, each landscape attractor is associated with a cell type, and cells in the landscape are discretely classified in accordance with their proximity to an attractor. Moreover, the particular normal form used to construct the model must be chosen *a priori*, requiring at least qualitative, prior understanding of the phenomenon.

### Parameterized landscape archetypes

In this section, we describe two parameterized landscapes, each given by a particular algebraic normal form and representing one of the classes enumerated by Rand et al. [39], termed the *binary choice* and the *binary flip*. These landscapes will serve three primary functions moving forward. First, they make concrete the ideas discussed above regarding Waddington’s landscape and the interpretation of signals as parameters shaping it. Second, inspired by Sáez et al. [40], who used both forms to construct a larger landscape model, we utilize these forms individually as synthetic testing grounds for our modeling framework, where the ground-truth data generating process is fully specified. Finally, these forms motivate a restriction of the broad class of parameterized landscapes to a smaller subset boasting a specific decomposition that allows us to algebraically separate the underlying landscape from a simple signal effect.

Both the binary choice and the binary flip are examples of a particularly simple type of parameterized landscape, in which the essential structure of the landscape is fixed, and the influence of the parameters in altering the landscape is linear. For this reason, we will refer to the parameters as *tilt parameters*, denoted ***τ***, and to these parameterized landscapes as *tiltable landscapes*. Moving forward, we restrict our consideration to this class of landscapes. Formally, we can express a tiltable landscape as the sum of a static, nonlinear portion 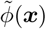 that does not depend on the parameters, and a linear term:

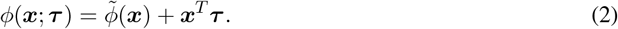

The binary choice landscape (Fig. 2a) is given by the normal form

**Figure 2.**
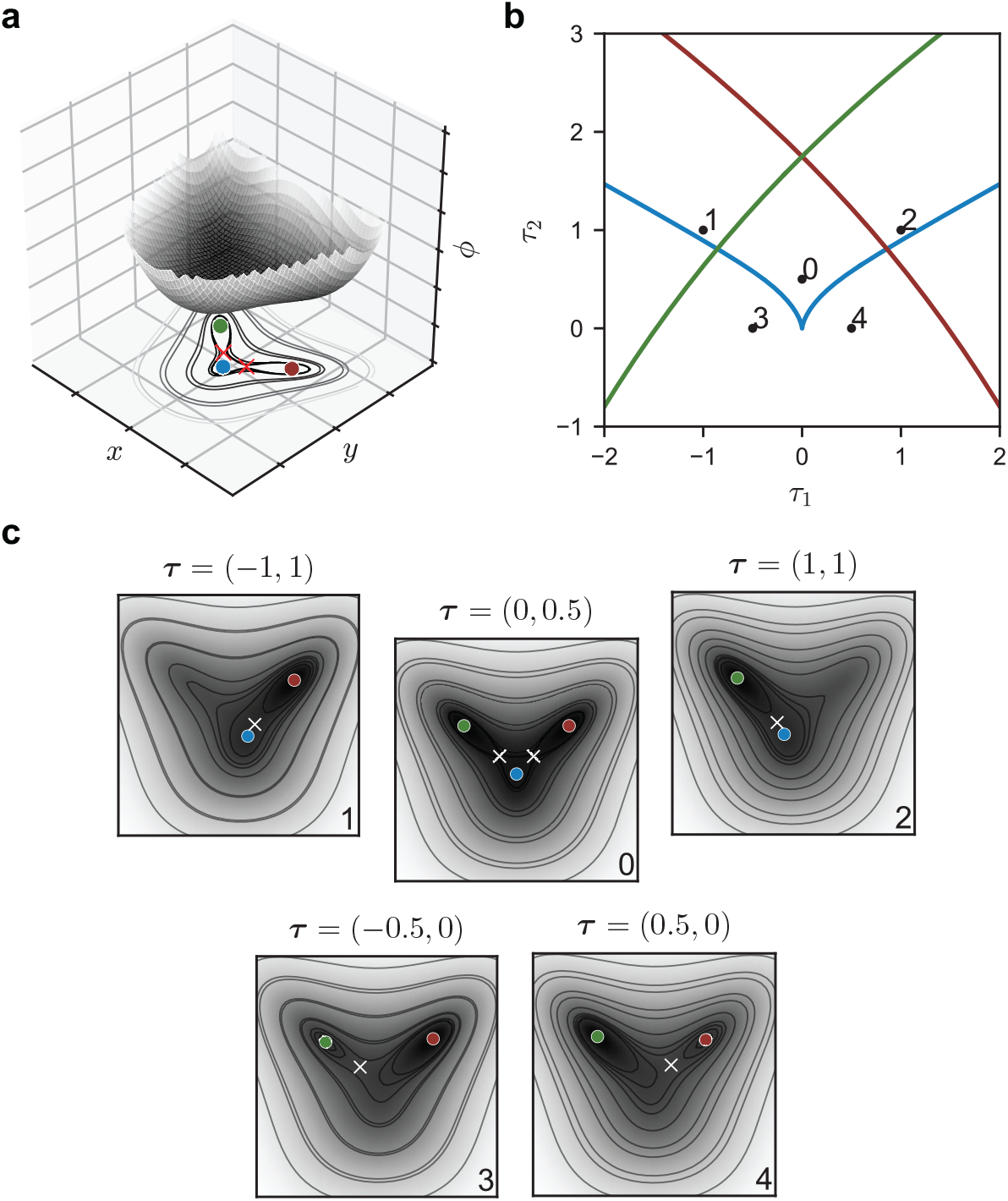
A parameterized landscape specified by an algebraic normal form exhibits bifurcations and models a binary decision. **(a)** An archetypal parameterized landscape, the binary choice [40]. The potential function is given by *ϕ*_*bc*_(*x, y*; ***τ***) = *x*^4^ + *y*^4^ + *y*^3^ −4*x*^2^*y* + *y*^2^ + *xτ*_1_ + *yτ*_2_. The landscape tilts as the vector ***τ*** changes, with the potential increasing in the direction of ***τ*** . The plotted landscape corresponds to the tilt vector ***τ*** = (0, 0.5), for which three attractors (circles) are separated by two saddles (crosses). **(b)** The potential *ϕ*_*bc*_ exhibits fold bifurcations at particular values of ***τ*** . The color of each fold curve corresponds to the fixed point that vanishes/appears as ***τ*** passes through it. In the central region, all three attractors are present. **(c)** The landscape given different values of the tilt ***τ*** . Each plot corresponds to a position in parameter space, as marked in panel (b). Minima are denoted with a circle, and saddle points with a cross. Of particular importance are the bifurcations that have occurred in subpanels 3 and 4, in which the central attractor bifurcates with one of the neighboring saddles.

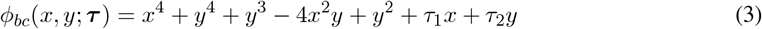

and admits fold bifurcations (a.k.a. saddle-node bifurcations) as *τ*_1_ and *τ*_2_ vary. In a generic fold bifurcation, a saddle point and an attractor or repeller either coalesce and vanish (the subcritical case) or appear together (the supercritical case). There is a central region of parameter space where three stable fixed points exist. These are configured such that a central attractor is separated from two peripheral attractors, with saddles in between (Figures 2b,c). The unstable manifold of a saddle connects the central attractor to one of the two peripheral ones. As *τ*_1_ and *τ*_2_ vary, the fixed points move and the saddles may coalesce with the attractors, resulting in the bifurcation diagram shown in Fig. 2b.

The second example, the binary flip landscape, is given by

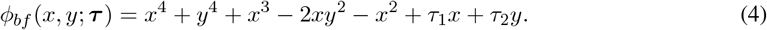

In this case, the configuration of the attractors and saddles is different. One saddle separates a central attractor from the two peripheral ones, and another saddle separates the two peripheral attractors from each other (Fig. S2). While saddle-node bifurcations can still occur, there is also the possibility of a *heteroclinic flip* bifurcation. In this global bifurcation, the unstable manifold of one saddle coalesces with the stable manifold of another, so that there is a heteroclinic orbit connecting one saddle to the other (see Supplementary Materials S1 for definitions). While all attractors persist smoothly through the bifurcation, there is a sudden change in the unstable manifold of the saddle separating the central and peripheral attractors, as it jumps abruptly from connecting the saddle to one instead of the other.

An essential difference between the binary choice and binary flip is the possible connection between the two peripheral attractors. In the case of the binary choice, a bifurcation that results in the vanishing of a peripheral attractor leads cells in its vicinity to return to the central attractor; there is no route directly to the remaining periphery. In the binary flip, however, the peripheral attractors are connected via the unstable manifold of a saddle, and this provides a route from the vicinity of one periphery to the other. This difference in the global arrangement of bifurcations between each archetypal landscape class motivates the use of the binary choice as a model for an “all-or-nothing” transition in which all cells move toward one periphery. The binary flip, in constrast, models a more flexible fate decision [39].

A final note must be made in regards to the role of signaling, as thus far the parameters appearing in the normal forms have lacked any biological meaning. In Sáez et al. [40] the landscape parameters ***τ*** serve as intermediates between a set of signals in the system and their effect on the underlying landscape. Conceptually, this captures the notion that a cell interprets a given set of chemical signals, which then informs, or biases, its fate decision. Mathematically, we express the tilt ***τ*** as a function of the signal vector ***s* ∈**ℝ^*d*^, where *d* is the number of signals present in the system, and write ***τ*** = *ψ*(***s***), where *ψ* : ℝ^*d*^ *→*ℝ^*d*^ is a function describing how a cell processes the signal. Implicit here is our assumption that the number of signals is equivalent to the dimensionality of the landscape. We can then express the tiltable landscape in terms of the signals directly:

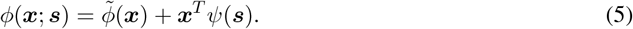

Now, specifying a signal profile as a function of time describes the temporal change of the landscape, as it tilts in response to the signal. In this way the signals act as forces, biasing the flow of cells in the landscape in particular directions, according to the currently abstract function *ψ*.

## Results

### A model encapsulating a tiltable landscape

We present a model architecture to capture gradient dynamics of stochastic cells in a landscape that changes in response to a set of external signals, what we term a *Parameterized Landscape Neural Network*, or PLNN. This model can be broken down into three core modules, each of which captures an aspect of a tiltable landscape: an underlying static potential function, a map defining the effect of signals on the landscape, and a noise kernel capturing stochasticity. Throughout, we denote the learnable parameters of this model by ***θ***. A schematic of this architecture is shown in Fig. 3a.

**Figure 3.**
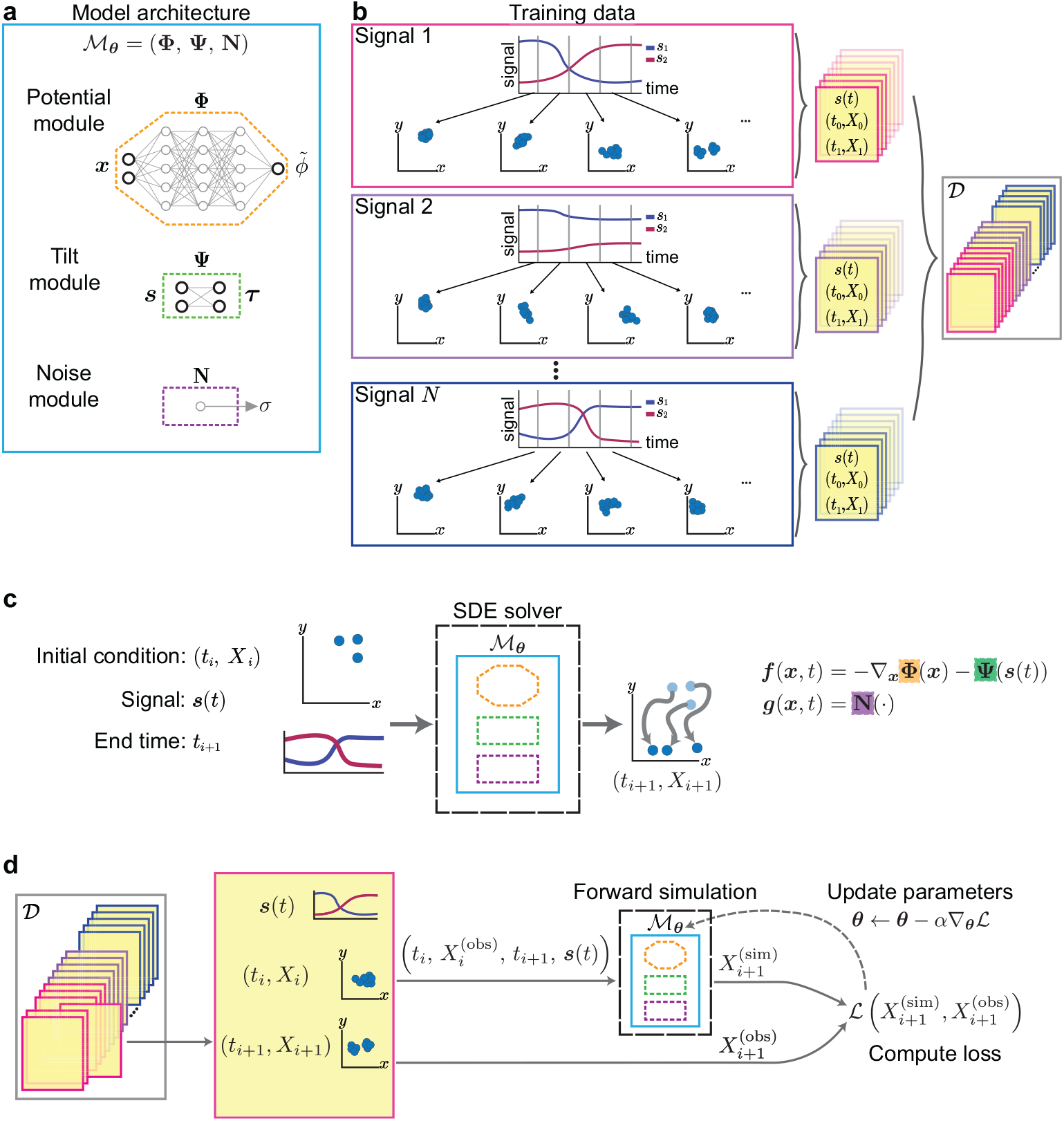
A proposed model architecture and the training process through which model parameters are inferred. **(a)** The PLNN model architecture. Three components capture the essential features of a tiltable landscape. The learnable parameters ***θ*** of the model ℳ_***θ***_ include the hidden weights and biases of the neural network **Φ**, the weights of the linear transformation **Ψ**, and the scalar noise parameter *σ*. **(b)** Training data comes from a number of experiments in which a cell ensemble is sampled at different timepoints under the condition of a particular signal profile. Two samples taken at different times constitute an initial/final state datapoint. The datapoints from each experiment are pooled together to form the training dataset 𝒟. Within a given experiment all datapoints share the same signal profile ***s***(*t*). **(c)** The inputs and outputs of a model simulation. The inputs are an initial ensemble state *X*_*i*_ at time *t*_*i*_, a signaling timecourse defined over an interval [*t*_*i*_, *t*_*i*+1_], and the terminal time *t*_*i*+1_. The model ℳis wrapped in a SDE solver that internally solves the SDE forward in time in order to sample a trajectory for each cell. The output is the simulated final state *X*_*i*+1_ corresponding to time *t*_*i*+1_. The governing equation of the SDE involves a deterministic function ***f*** and a stochastic function ***g***, each built from components of the model ℳ . **(d)** The PLNN training procedure. A datapoint is processed as in (c) to return a *simulated* final state 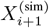.This is compared to the *observed* final state 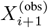 using a suitable loss function. Backpropagation is then used to compute the gradient of this loss with respect to the model parameters, which are then updated. In practice, this procedure is vectorized and performed in batches.

The first module is the *potential module*, **Φ** = **Φ**_***θ***_, and it is intended to represent an underlying static potential 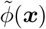 defined atop a *d*-dimensional phase space. Biologically speaking, this is the zero-signal state of the landscape. In order to represent a smooth, nonlinear potential function, we use a neural network architecture as part of our construction. We denote by 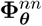 a feedforward neural network whose weights and biases are among the learnable model parameters ***θ***. So that we may compute its gradient—the dynamical quantity of interest—we use a smooth activation function in order to guarantee smoothness of the potential. We further regularize the potential module by including an additional *confinement term*, denoted Φ_0_. This regularization is meant to guarantee that trajectories are confined and do not escape to infinity during the training process. We define

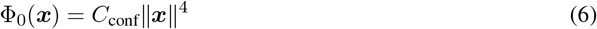

where *C*_conf_ is a non-negative constant that serves to greaten or lessen the degree of regularization. (The confinement factor *C*_conf_ is a hyperparameter, and not a learned model parameter.) Then, the complete potential module is given by the sum

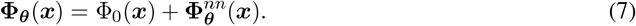

The second module captures the effect of signals on the underlying landscape, and we denote it by **Ψ** = **Ψ**_***θ***_. This module defines a transformation that maps a vector of signal values ***s*∈** ℝ^*d*^ to a vector ***τ* ∈**ℝ^*d*^, the effective tilt of the landscape. In parallel with (5), given a signal vector ***s***, the height of the tilted landscape is given by

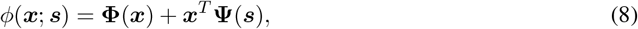

respectively.

By design, the resulting tilt ***τ*** = **Ψ**(***s***) acts linearly on the underlying landscape. In addition, we assume the signal processing function **Ψ** is linear, and write

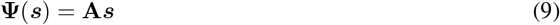

where **A** is a *d*× *d* matrix defining the linear map from signals to tilt. Enforcing a linear processing of the signals in addition to a linear effect of the resulting vector on the landscape passes all nonlinearities to the potential module **Φ** representing the underlying landscape 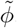.To learn the transformation **Ψ** is to learn the effect of each signal on each dimension of the underlying landscape. This transformation is the same across the entire landscape, and raises the question as to how to interpret the impact of the signal within the vicinity of each fixed point. By subsuming all of the nonlinearities under the potential module, the interpretation is that the structure of the landscape will inform the exact nature of the role of the signal in local regions of phase space.

Finally, the third component of our model captures stochasticity in the system, and we denote it by **N** = **N**_***θ***_. In general, this module maps a cell’s state ***x*** to a noise kernel, Σ(***x***) ∈ℝ^*d×d*^. We make a number of simplifying assumptions with regard to the system noise, but these assumptions may be weakened. We assume that stochasticity in each dimension is the result of an independent one-dimensional Wiener process, and that the noise is additive (i.e. it does not depend on the state). Moreover, we assume that a global parameter *σ* governs the magnitude of isotropic noise in the system. Together, these assumptions result in a constant, diagonal noise kernel

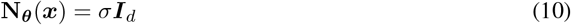

where ***I***_*d*_ is the *d* ×*d* identity matrix.

To summarize, our model is parameterized by the weights and biases of the neural network **Φ**_***θ***_, the weights of the linear transformation **Ψ**_***θ***_ : ℝ^*d*^ *→*ℝ^*d*^, and the global noise parameter *σ*. Further specifics of the model architecture, including the number of network layers and choice of activation function, can be found in Methods.

The modules of the PLNN provide all of the information needed to define a stochastic differential equation (SDE) governing the evolution of an ensemble of cells within the represented landscape, over an interval of time 0≤ *t*≤ *T*, as depicted in Fig. 3c A general SDE contains a drift and a diffusion term, denoted ***f*** (*t, x*) and ***g***(*t, x*), respectively, and can be written in the form

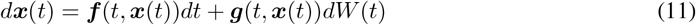

where *W* is a *d*_*w*_-dimensional Wiener process [43]. In this case, the drift term ***f*** is the deterministic flow induced by the potential, tilted in response to a particular signal. For a signal profile ***s***(*t*), this flow is given by the negative gradient of (8):

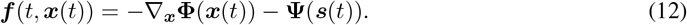

Under the assumption of isotropic additive noise, *d*_*w*_ = *d* and the diffusion term is given by the output of the noise module **N**:

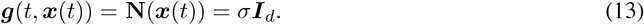

With these equations, given an initial cell state ***x***_0_ = ***x***(0), we sample a future state ***x***(*t*) by solving the SDE forward in time using, for example, the Euler-Maruyama. As we are interested in the evolution of not just a single cell, but an ensemble of cells, we simulate an array of SDEs

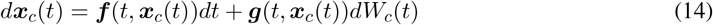

where *c* = 1, …, *N*_cells_ indexes across the total number of cells. Each cell is subject to an independent *d*-dimensional Brownian motion *W*_*c*_. We solve the resulting system of SDEs numerically, in a manner compatible with backpropagation, allowing us to differentiate through the operations of the solver (Methods). In this way, we formulate a generative model that we can train via backpropagation and standard machine learning techniques, discussed in more detail below.

### A landscape model inferred from synthetic data

We begin by showing that we can infer a nonlinear potential function and the way it tilts in response to two signals. We do this using *in silico* experiments where the potential we wish to infer, denoted *ϕ*^∗^, is known *a priori*, and used to generate synthetic data on which our model will be trained. To this end, we leverage the binary choice landscape (3), and define a ground-truth parameterized landscape

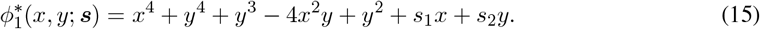

(The case of the binary flip landscape is provided in Supplementary Materials S1.)

We synthesize a training dataset from the ground-truth data generating process. Taking 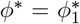,we perform a series of *in silico* experiments in which cells are observed at discrete points in time as they flow through the changing landscape (Methods). Each experiment involves simulating an ensemble of cells in the landscape, under the influence of a prescribed signal profile drawn from a prior distribution of feasible signal functions. The state of the ensemble is recorded at a number of sampling timepoints, from which we compile a set of consecutive timepoint-state pairs of the form (*t*_0_, *X*_0_; *t*_1_, *X*_1_; ***s***(*t*)), where *X*_0_ and *X*_1_ are the states of the ensemble at times *t*_0_ and *t*_1_, respectively. In addition, we record the parameterization of the signal profile, ***s***(*t*), so that we may compute the value of the signal at any time *t*. The data generated across all of the performed experiments are pooled, resulting in a synthetic training dataset 𝒟 (Fig. 3b).

We next train a PLNN on the synthetic data generated in the ground-truth landscape, as detailed in Methods. The training process simulates an ensemble of cells from the observed initial state at time *t*_*i*_, to the next timepoint *t*_*i*+1_ (Fig. 3c). The model parameters are updated based on the value of a specified loss function, that compares the *simulated* final condition to the *observed* (ground-truth) final condition, in a distributional sense (Fig. 3d). To this end, we utilize a loss function that computes from empirical samples an unbiased estimate of the maximum mean discrepancy (MMD) [44] between two distributions (Methods).

Fig. 4b depicts the landscape inferred from data generated in the binary choice system, 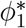. The inferred landscape is shown alongside the ground-truth landscape. Qualitatively, the inferred landscape resembles the ground-truth, with three fixed points separated by two saddles. We expect that the difference between the actual and inferred landscapes will be larger in the regions of state space with a lower density of observed cells across all of the training data. In those regions that are not in the support of the training data, we expect a poorer fit. This conjecture is supported by the level sets of the inferred landscape, which become more distorted away from the central region of attractors. The inferred linear map of signals to tilts is also depicted within the landscape plots, as arrows showing the direction each independent signal biases the flow. (Note that this direction is opposite the tilt vector, as we define it.) The inferred model captures both the magnitude and direction of the effect of both signals.

**Figure 4.**
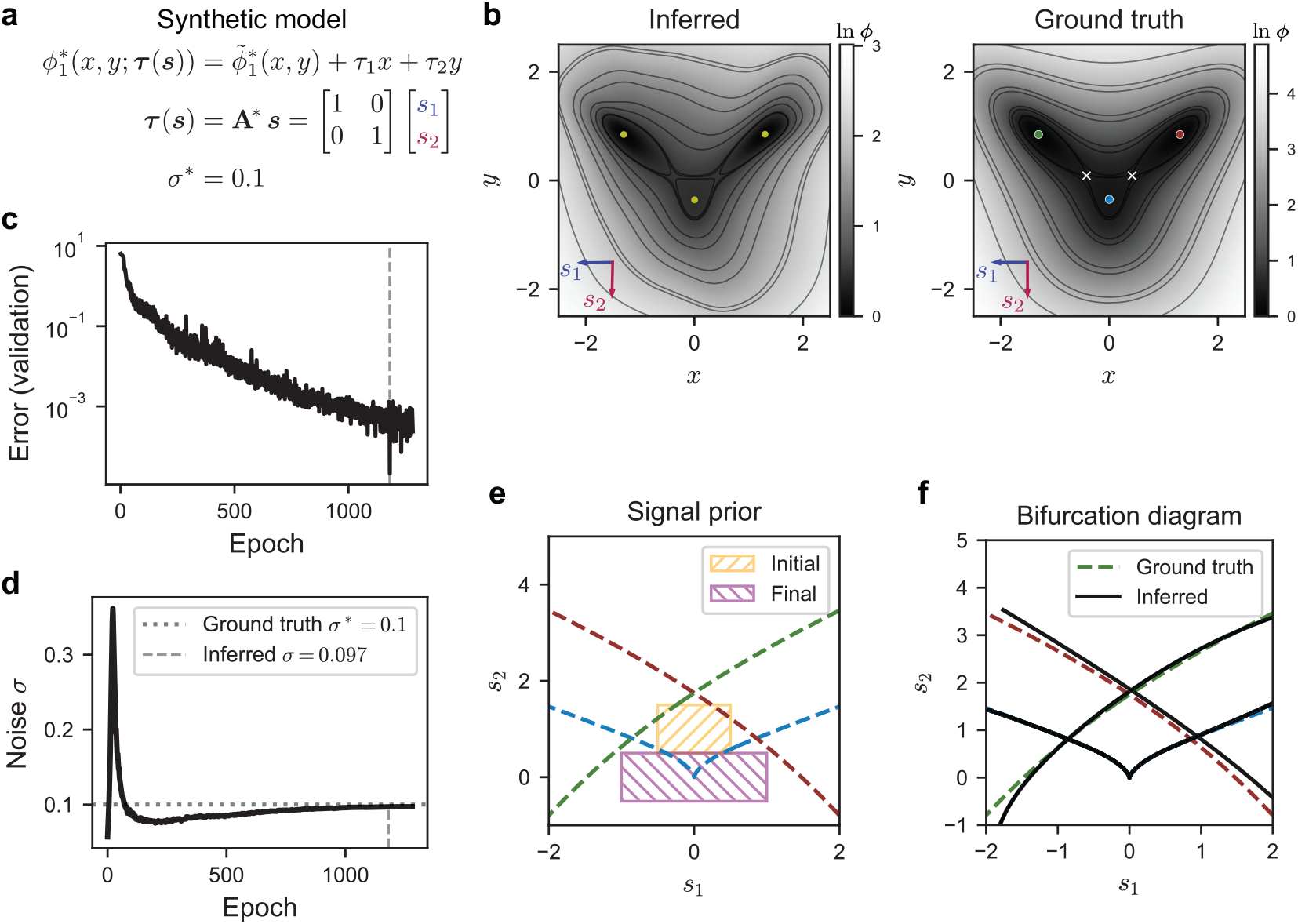
A trained model captures the dynamics and bifurcation structure of the binary choice landscape. **(a)** The algebraic form of the binary choice system, the ground-truth model generating the data used for training. This system tilts as the signals *s*_1_ and *s*_2_ vary. In the ground-truth system, the signals map identically to the tilt parameters, and we specify a noise level *σ*^∗^ = 0.1. **(b)** The ground-truth and inferred landscapes. The effects of the signals *s*_1_ and *s*_2_ are depicted as arrows in the lower left of each plot, pointing in the direction in which flow is biased when a unit signal vector is prescribed. The landscapes agree qualitatively, with larger disparities occurring in regions containing few cells from the training data, as evidenced by the differences in the contours shown. **(c)** The value of the validation loss, or error, over the course of training. The dashed line indicates the epoch corresponding to the optimal model, exhibiting the minimal validation loss. **(d)** The evolution of the inferred noise parameter *σ* during the training process. The true value, *σ*^∗^, is denoted with a horizontal line. The solid vertical line corresponds to the optimal model found during training, with an inferred *σ ≈*0.097. An initial phase of model training witnesses a rapid increase and decrease in *σ*, followed by a slower approach towards the true value. **(e)** The bifurcation diagram of the ground-truth parameterized landscape 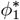,with the color of each fold curve corresponding to the color of the fixed point in panel (b) that vanishes when the signal passes through the curve. The hatched rectangles illustrate the range of sigmoidal signal functions used to generate the synthetic training data. They indicate the regions of parameter space from which the initial and final signal values are drawn, so that ***s***(*t*) passes from the central region into the lower region, crossing the fold curve corresponding to the bifurcation of the central attractor. **(f)** A comparison of the bifurcation diagrams corresponding to the ground-truth and inferred landscapes. Dashed lines correspond to the ground-truth system. Solid lines correspond to the inferred model. There is general agreement between the true and inferred fold curve with a cusp (depicted in blue) since this is the bifurcation captured by the training data. In addition, two inferred fold curves approximate those corresponding to the peripheral attractors.

In regards to the noise parameter *σ*, the training process does appear to converge toward the true value *σ*^∗^ = 0.1, as shown in Fig. 4d. Notably, we observe a transient phase at the outset of training during which the noise level increases past the true value, before converging back towards it. We suspect that this phase is necessary to allow cells to occupy a wider region of phase space, without which there is no chance of correctly inferring the landscape topology. Once the cells are sufficiently distributed throughout the landscape, the landscape is able to adjust in order to capture the particular features of the distribution. We note that in some instances the noise parameter, while converging, does not converge to the ground-truth level. We expect this is due to the degeneracy between the shape of the landscape and the noise level. While we can fix the noise parameter at an arbitrary level and infer only the landscape shape, this prevents the transient phase discussed above and inhibits the training process. For this reason, we leave the noise parameter free, and assess the accuracy of the model by analyzing the distribution of cells around stable fixed points.

The agreement between the ground-truth and inferred parameterized landscapes is further evidenced by comparing the bifurcation diagrams of the two dynamical systems. Fig. 4f depicts the fold curves of the ground-truth and inferred landscapes, which constitute the locus of signal values at which the landscape exhibits a saddle-node bifurcation. The bifurcation diagram for the inferred landscape is found using a pseudo-arclength continuation algorithm [45, 46], details of which can be found in Supplementary Materials SA. There is particularly close agreement with respect to the fold curve corresponding to the bifurcation of the central attractor, depicted as a dashed, blue line, and containing a dual cusp point. This is explained by the particular sigmoidal signal profiles used to generate the training data. As shown in Fig. 4e, the initial signal value falls within a region of signal space almost guaranteeing the presence of the central attractor, whereas the final value is largely confined to a region where the central attractor has vanished in a bifurcation with one of the neighboring saddle points. Thus, the bifurcation of the central attractor is well-captured in the training data, whereas the bifurcations of either peripheral attractor are not. In light of this, it is perhaps surprising that there is still relatively good agreement between the fold curves denoting the peripheral attractor bifurcations (shown in red and green). That they exist at all, and agree qualitatively, is indicative of the robust and generic structure of the archetypal landscape form. In some cases, the bifurcation diagram contains additional fold curves, indicating the presence of additional fixed points in the inferred model. These additional minima occur in regions outside of the support of the training data. This suggests again that while anomalies may occur, they tend to do so in regions of phase space that are never occupied by cells during the training process, and therefore for which there is no information to inform the landscape shape.

### Robustness of landscape inference to temporal sampling resolution

A number of factors influence our ability to infer a landscape system from data. Among these is the frequency of temporal sampling, which must be sufficient to capture the transition of cells between states. This is an important consideration, especially for systems in which the transition rate of cells is high. Here we examine the degree to which model inference is robust to the temporal sampling rate used to obtain data.

Fig. 5 depicts a collection of models trained on a series of three different datasets, each generated using the binary choice system (15), but with varying rates of sampling (Methods). Each dataset is constructed from a number of synthetic experiments over the interval 0 ≤ *t* ≤ 20, with snapshots taken at sampling intervals of Δ*T* ∈ {5, 10, 20}. Each is composed of a different number of experiments, in order to ensure that the total number of snapshots (i.e. training datapoints) is the same in each case. The three training datasets are depicted in Fig. 5a, highlighting the qualitative differences between them. Sparser sampling results in fewer cells captured in the transition between attractors. That is, the unstable manifolds of the saddles are poorly resolved.

**Figure 5.**
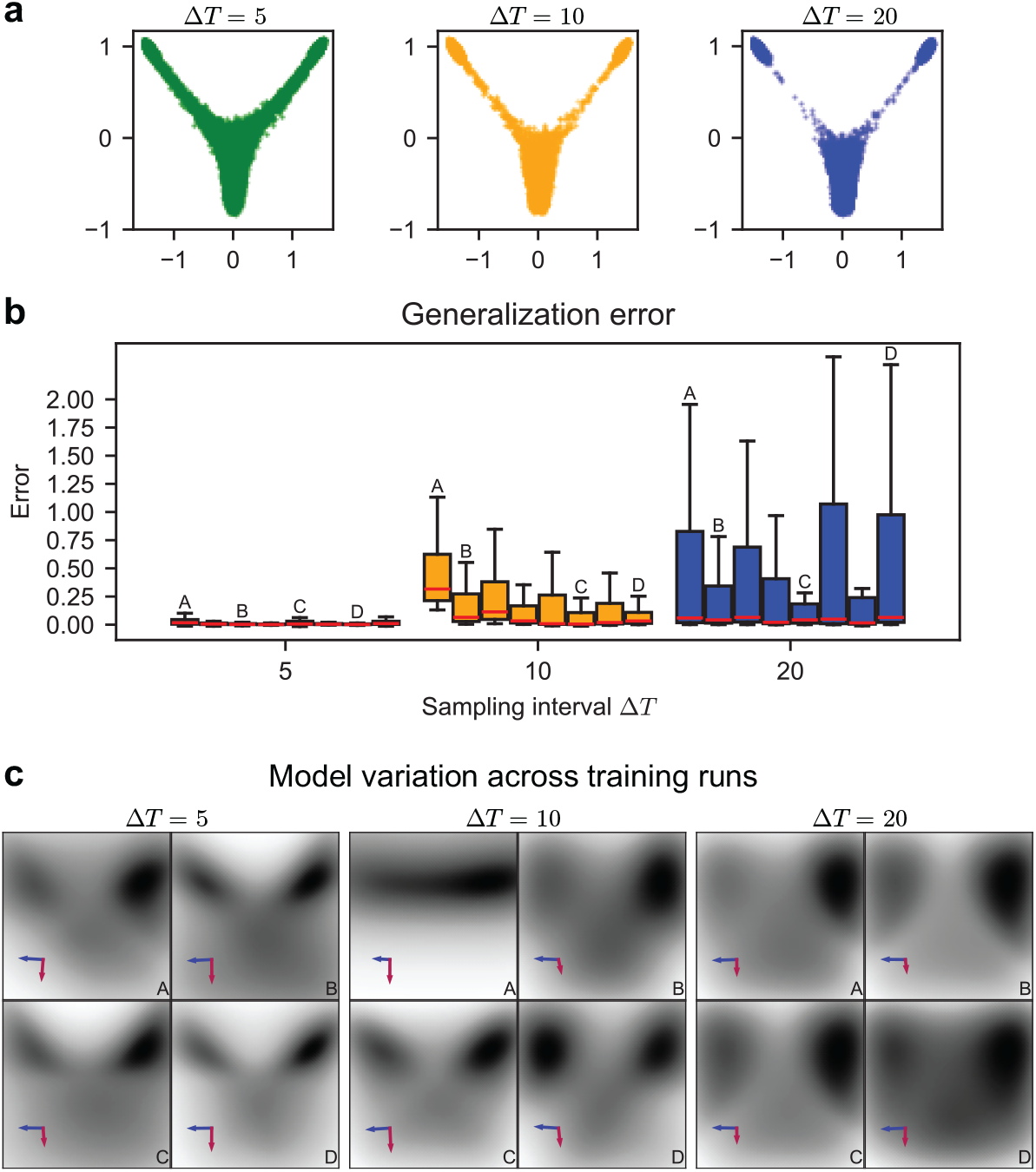
Landscape features vary in their degree of robustness to the sampling rate. **(a)** Three datasets generated from the binary choice landscape, simulated over the interval 0≤ *t* ≤ 20, with cells sampled at varying intervals. The plots display all cells making up a dataset, consisting of pooled experimental data. Each dataset consists of a different number of experiments, so that the total number of consecutive timepoint samplings is consistent. Although the same number of cells are depicted in each case, a shorter sampling interval Δ*T*, the duration between snapshots, provides more information about the routes between attractors. **(b)** For each dataset, eight models were trained and then evaluated against the same held-out dataset. Each box and whisker plot corresponds to a single model, and depicts the distribution of the generalization error (the MMD loss) across all evaluation datapoints. Note that outliers are excluded. Models trained on data with a higher rate of sampling perform better and more consistently. Letters correspond to the displayed models, below. **(c)** A subset of the inferred models, for each of the three sampling intervals. Most models accurately identify the stable fixed points of the ground-truth system, but those trained on data with low sampling resolution fail to correctly capture transition routes between states, corresponding to the unstable manifolds of saddles.

We trained a number of models on each dataset, and assessed the performance of each on the same held-out evaluation dataset, generated via the same process as the training data, using a sampling interval Δ*T* = 5. We test each model by applying it to every datapoint in the evaluation dataset, and computing the corresponding MMD loss. In this way, we assess a model by examining the distribution of the loss across every consecutive timepoint-state pair, providing a measure of generalization error. As expected, models trained on more densely sampled data (Δ*T* = 5) perform better on the evaluation dataset, with nearly all datapoints resulting in a loss below 0.1 (Fig. 5b). While the median loss of models trained on the dataset with a longer sampling interval of Δ*T* = 10 is consistently below 0.1, there is more variability in model performance, with half of the models incurring a greater loss on a significant proportion of the evaluation data. Notability, these models perform nearly as well on half of the data as the models trained on more frequently sampled data.

Finally, in the case of Δ*T* = 20, the training data consists of only the initial and terminal condition for each synthetic experiment, and the resulting data is much sparser in regions between attractors. Still, a number of peregrinating cells are captured, and while models trained in this case again perform more variably than those trained on denser data, it remains the case that some models perform well, with a majority of datapoints resulting in a loss below 0.2.

### Identifying a two-dimensional decision manifold in an *in vitro* dataset

Having shown that the model architecture is able to adequately capture elements of a synthetic cellular decision process, we now turn our consideration to the case of real data. Our purpose here is to demonstrate a framework with which a low-dimensional landscape model can be inferred from real data, and to illustrate the necessary considerations and current limitations of this approach. We start by outlining a general approach to working with high-dimensional data capturing a simple binary decision. We then apply this methodology to an *in vitro* dataset.

Fig. 6 summarizes a general procedure for handling high-dimensional data capturing a binary decision. We assume that high-dimensional timecourse data is collected from a number of experiments, each corresponding to a particular signal, or experimental condition. In Fig. 6a, such data is depicted in the differently-colored scatterplots (three-dimensional for the sake of example, and generated via a toy model, details of which are immaterial). Ideally, we want to identify a two-dimensional manifold within this high-dimensional expression space along which the controllable dynamics take place, and on which we can infer a landscape potential. The following approach illustrates a method that preserves an interpretation of cells in terms of the high-dimensional state space, using principal component analysis (PCA), and does not rely on any assignment of cell type or other clustering procedure to associate cells with a particular state.

**Figure 6.**
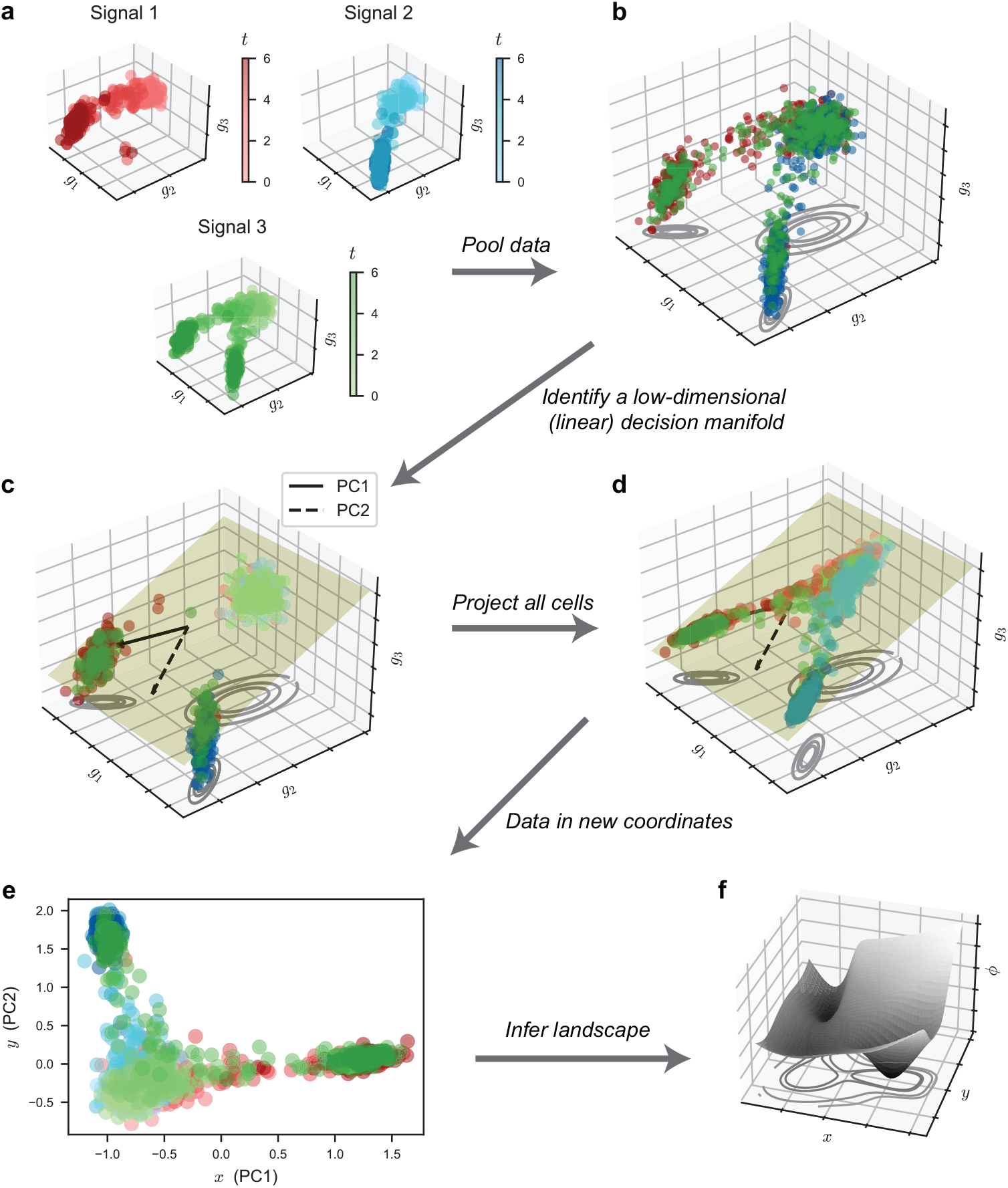
Identifying a two-dimensional decision manifold that captures the relevant cellular dynamics and underlies an inferred landscape model. **(a)** A collection of experiments, in which an ensemble of potentially high-dimensional cells have been sampled at a number of timepoints. Here we illustrate the profiled cells as points in a 3-dimensional gene space, sampled at times *t*∈{ 0, 2, 4, 6} . Each experimental condition is shown in a different color, and depicts cells moving noisily near the surface of a sphere. **(b)** The experimental conditions, when pooled, are assumed to lie along a curved manifold within the high-dimensional gene space, with identifiable initial and terminal cell types. Here, the plotted contours depict three identifiable clusters. **(c)** The subset of cells across all experimental conditions captured at the initial and final times are used to determine a linear decision manifold, an approximation to the supposed nonlinear, true manifold. We illustrate this step using the first two principal components found via PCA, which define a plane passing through the average cell. Cell states should be distinguishable along these axes. **(d)** Once the linear decision manifold is found, we project all cells—including those sampled at intermediate times—onto this plane, capturing the relevant transitions. **(e)** The resulting two-dimensional data, in this case given by the coordinates of each cell along the first two principal components, serve as training data with which to infer a PLNN. **(f)** An inferred landscape model overlays the decision manifold, and offers an intuitive picture of the dynamics in an interpretable space.

Under the assumption that the data capture a binary decision involving three cell states, these states should be discernible when the data is pooled across all experimental conditions (Fig. 6b). These states, as three attractors in phase space, define a 2-dimensional plane. While cells may transition along paths that are not restricted to this plane, the projection of the cells onto the plane should capture an approximation of the dynamics. Rather than attempting to identify the attractor states directly, we instead isolate the cells at the start and end of the experiment, as we expect these cells to be primarily settled around attractors that correspond to the initial and terminal cell states, respectively. We then construct a plane through these states using PCA. We apply PCA to the initial and terminal cells, and define a linear decision manifold by the plane spanned by the first two PCs and passing through the average cell (Fig. 6c). Once we determine this decision manifold, we project all of the observed cells onto it, and take the coordinates of each cell within this manifold as the (*x, y*) landscape coordinates (Fig. 6d-e). We then infer a landscape model defined on this manifold (Fig. 6f).

In order to demonstrate this approach on real data, we apply our framework to the *in vitro* dataset originally detailed in Sáez et al. [40]. This dataset captures an array of experiments, in which mouse embryonic stem cells (mESCs) were exposed to two chemical signals, WNT and FGF, for different durations over a span of five days. Starting at Day 2 (D2), the experimenters used flow cytometry and fluorescence-activated cell sorting (FACS) to profile the expression levels of five marker proteins at intervals of 12 hours. From the resulting FACS data, they identified five primary cell states arranged in two consecutive binary decisions. We utilize the five-dimensional FACS dataset and infer a landscape model for each of the two binary decisions captured therein. We briefly summarize the two binary decisions, and refer the reader to the original work for additional biological context.

After two days of exposure to FGF, mESCs take on an identity similar to that of early epiblast cells. This epiblast-like (EPI) identity serves as the initial state of cells in the system. Removing exogenous FGF results in EPI cells transitioning to an anterior neural (AN) identity. On the other hand, the activation of WNT signaling at D2 via addition of CHIRON99021 (CHIR) causes the EPI cells to instead transition to a caudal epiblast (CE) state, passing through an intermediate transitory (Tr) state. The CE state is shared between the two binary decisions, and marks the central attractor for the second. Withdrawing both the FGF and WNT signal results in CE cells transitioning to a posterior neural (PN) state, while a sustained WNT signal causes them to take on a paraxial mesoderm identity (M). Whereas in the first decision all cells either transition to the AN state or to the CE state (a binary choice), in this second decision, some cells may transition to the PN state while others transition to the M state (a binary flip), with the relative proportion governed by the duration of FGF signaling in addition to WNT.

An essential assumption in the original work [40], and in our treatment of the data, is a complete specification of the signal profile used in each experiment. The level of CHIR (in effect, the WNT signal) is treated as a binary function taking a value of either 0 or 1. That is, the WNT signal is either entirely absent, or fully saturated. On the other hand, the level of FGF varies between three discrete levels: blocked, partially active due to endogenous production, and fully saturated due to exogenous FGF. We use the same numerical prescription for the FGF signal function as in the original work, with the off, partial, and saturated levels given by 0, 0.9, and 1, respectively. The 0 level for FGF is obtained by prescription of a third chemical, PD, that blocks endogenous FGF production. However, we still treat the system as governed by only two signals, the effective level of FGF and WNT. Table 1 lists each experimental condition and the corresponding signal profile, as well as its use case (model training, validation, or evaluation). Note that while the signals are prescribed as step functions, we smooth them using a sigmoidal parameterization with a high transition rate.

**Table 1:** The 11 experimental conditions constituting the mESC *in vitro* dataset. The prescription of CHIR, FGF, and PD, over time results in a particular signal profile for each condition. The signals are piecewise constant, with *χ*[*t*_1_, *t*_2_] equal to one on the interval [*t*_1_, *t*_2_) and zero elsewhere. Each condition is used as part of a training, validation, or evaluation set. ^∗^The NO CHIR condition is excluded from the training dataset in the case of the second decision.

| Condition | Signal | Dataset |
| --- | --- | --- |
| NO CHIR | CHIR = 0<br>FGF = $1\chi[0, 3] + 0.9\chi[3, 5]$ | Training* |
| CHIR 2–2.5 | CHIR = $1\chi[2, 2.5]$<br>FGF = $1\chi[0, 3] + 0.9\chi[3, 5]$ | Validation |
| CHIR 2–3 | CHIR = $1\chi[2, 3]$<br>FGF = $1\chi[0, 3] + 0.9\chi[3, 5]$ | Training |
| CHIR 2–3.5 | CHIR = $1\chi[2, 3.5]$<br>FGF = $1\chi[0, 3] + 0.9\chi[3, 5]$ | Validation |
| CHIR 2–4 | CHIR = $1\chi[2, 4]$<br>FGF = $1\chi[0, 3] + 0.9\chi[3, 5]$ | Evaluation |
| CHIR 2–5 | CHIR = $1\chi[2, 5]$<br>FGF = $1\chi[0, 3] + 0.9\chi[3, 5]$ | Training |
| CHIR 2–5 FGF 2–3 | CHIR = $1\chi[2, 5]$<br>FGF = $1\chi[0, 3]$ | Training |
| CHIR 2–5 FGF 2–3.5 | CHIR = $1\chi[2, 5]$<br>FGF = $1\chi[0, 3.5]$ | Validation |
| CHIR 2–5 FGF 2–4 | CHIR = $1\chi[2, 5]$<br>FGF = $1\chi[0, 4]$ | Evaluation |
| CHIR 2–5 FGF 2–4.5 | CHIR = $1\chi[2, 5]$<br>FGF = $1\chi[0, 4.5]$ | Validation |
| CHIR 2–5 FGF 2–5 | CHIR = $1\chi[2, 5]$<br>FGF = $1\chi[0, 5]$ | Training |

We restrict our attention here to the first decision, and discuss the second in Supplementary Materials S5. In order to infer a landscape capturing the first binary decision, in which EPI cells transition to either the AN or CE state, we first recapitulate the classification algorithm detailed in the original work, assigning a cell type label to each cell (Methods). We then isolate those cells relevant to the initial transition, namely the EPI, AN, Tr, and CE cells. Next, using the general procedure described above, we reduce the dimensionality of the data, moving from the five-dimensional protein expression space to a two-dimensional space on which we can infer a PLNN. We apply PCA to the subset of cells sampled at D2 and D3.5, taking these times to correspond to the start and end of the first binary decision, since by D3.5 all cells have transitioned to the AN or CE states. We then use the first two principal components to define the coordinates of a two-dimensional plane onto which we project all of the observed cells. (Additional details in Methods.)

Fig. 7a shows two subsets of the resulting two-dimensional data, corresponding to the experimental conditions NO CHIR and CHIR 2–3 (see Table 1). The signal profile and resulting frequencies of each cell type are displayed above a series of snapshots of the data in the dimensionally-reduced (*x, y*) space. The first column below each frequency plot displays the density of the data in this space, capturing the evolution of the cell ensemble. The coloring included in the central columns, in accordance with each cell’s nominal type, allows us to associate regions of the (*x, y*) space, at each timepoint, to a particular cell state. Importantly, the data displayed in this column, which serve as input to the model during training, is colored independently from the landscape inference process, and the model is agnostic with respect to cell type labels. The rightmost columns show snapshots of a *predicted* flow of cells, generated from the resulting, inferred landscape model, described in the following section.

**Figure 7.**
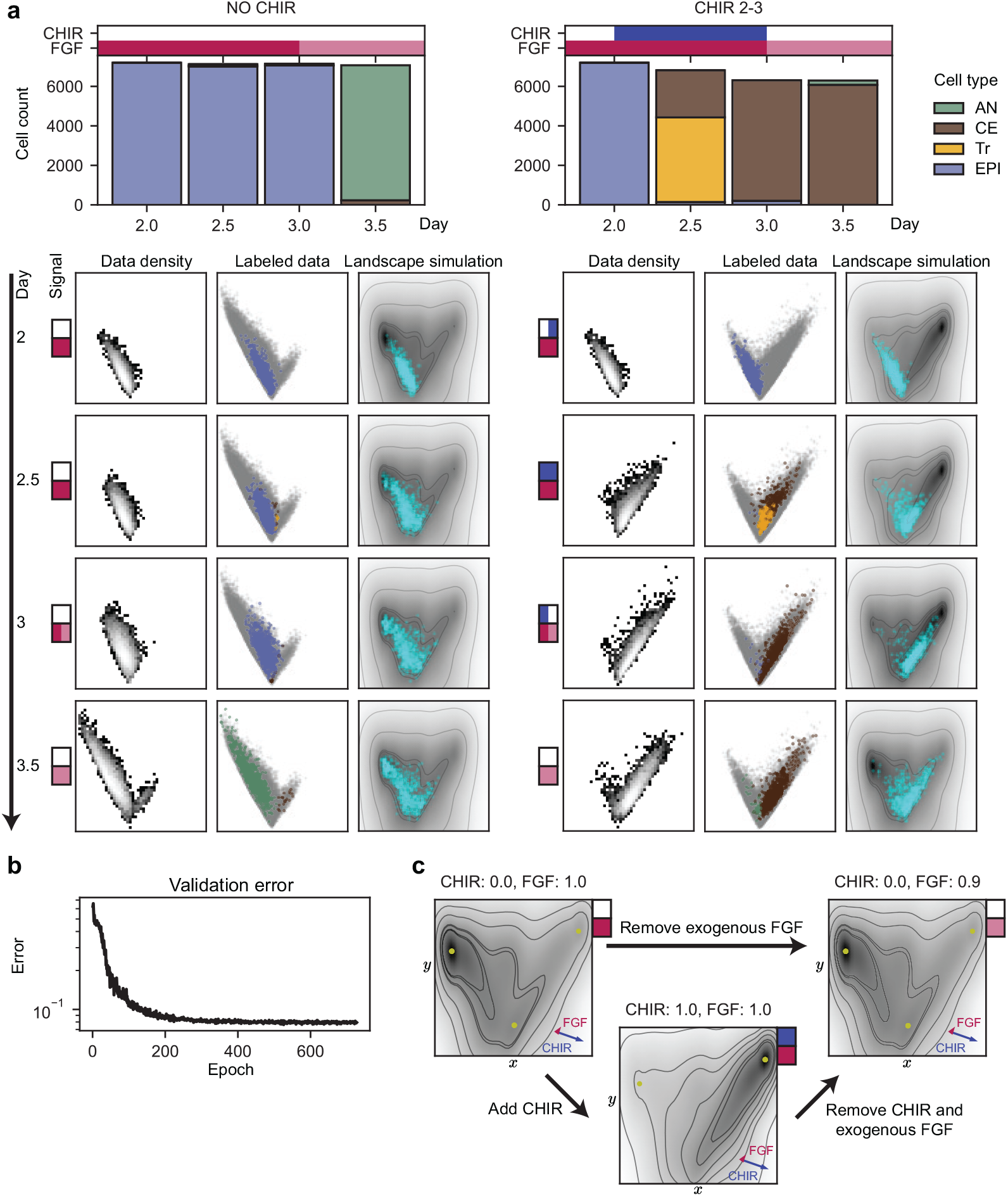
A parameterized landscape model inferred from *in vitro* experimental. **(a)** Two examples of the experimental conditions used for model training, each defined by a particular signal profile, and resulting in different proportions of each cell type across time. In the NO CHIR case, cells transition from the EPI state to the AN state. In the CHIR 2–3 case, cells move instead through the Tr state towards the CE state. Below, the temporal evolution of the cell ensemble is projected onto the two-dimensional decision manifold identified within the five-dimensional expression space. The colored squares depict the current signal level, with a change in signal depicted by a split coloring. The leftmost column shows a density plot of the cells at each timepoint. The center column shows a sample of cells, colored according to their nominal type. The background scatterplot, in grey, shows all cells associated with that experimental condition. The rightmost column shows snapshots of a simulation performed using the inferred model. **(b)** The validation error over the course of model training. **(c)** The inferred landscape tilted in response to different signals. Arrows in the lower right show the direction of bias resulting from addition of a unit CHIR or FGF signal. The effect of CHIR is evident, with its addition leading to the disappearance of the central attractor. The effect of FGF is less noticeable, and does not appear to qualitatively change the landscape.

The extent to which a linear decision manifold adequately captures the binary decision is questionable. It does appear from the labeled data that the initial population of EPI cells remains relatively stable under a constant FGF signal, and that when provided a CHIR signal, it transitions from the left half of the plane to the right. When the exogenous FGF signal halts, the distribution of EPI cells extends further into the upper-left region of the plane. However, there is not a clear separation between the EPI and AN cells when projected onto the linear decision manifold. We note here that the cell type assignment is performed in an *ad hoc* fashion, and involves a supervised assignment of cell types to particular components of a number of Gaussian mixture models, each corresponding to a particular timepoint (Methods). For this reason, the lack of distinguishability between the EPI and AN cells may simply be due to similarity between these cell types in the reduced two-dimensional space. Similarly, the transitioning population, identified only at D2.5, appears as expected between the EPI and CE states, but is not entirely distinguishable from either population.

### An inferred *in vitro* cellular decision landscape

With the dimensionally-reduced data in hand, we train a PLNN on a subset of the experimental conditions, holding out additional subsets for model validation and evaluation (Table 1). Due to the large and varying number of cells observed in each experiment, we perform an intermediate sampling step during the training process to fix the number of simulated cells (Methods). The resulting landscape is shown in Fig. 7c. We plot the landscape tilted in response to a number of relevant signals and identify local minima within the extent of the data. In the presence of FGF without a CHIR signal, ***s*** = (*s*_CHIR_, *s*_FGF_) = (0, 1), we identify three local minima. Given the initial ensemble state, cells primarily occupy a region between the central and leftmost attractors, a region associated with EPI and AN cells. There is significant overlap in the region of state space associated with these cell types, indicating that the dimension reduction is insufficient to entirely separate the two states. The central attractor coincides with the EPI and transitioning (Tr) cell state, and the rightmost attractor with a region occupied by CE cells.

The addition of a CHIR signal leads to the bifurcation of the central attractor, leaving the presumable AN and CE attractors, as expected from a binary choice decision. The inferred effect of CHIR on the landscape is greater in magnitude than that of FGF. A reduction in the level of FGF from 1 to 0.9, representing removal of the exogenous signal, does not appear to result in a significant qualitative change to the landscape. As CHIR is largely responsible for the dynamics of the first decision, this is unsurprising. While the effect of FGF on the landscape is minimal, it is at least qualitatively consistent with the distribution of the training data on the decision manifold. As we see from the direction of bias resulting from a unit FGF signal (Fig. 7c), removal of FGF results in a slight bias away from the CE state, and towards the AN one, consistent with cells transitioning from the initial EPI region to the AN region.

The level of noise inferred by the model is substantial (*σ* ≈ 1.8), though consistent with the degree of variation of cells around presumable attractors. This degree of noise may inhibit the association of identified minima with specific cell states.

After training, we generate sample trajectories of a cell ensemble within the landscape, under both of the signal profiles depicted in Fig. 7a, and display snapshots of the ensemble at the sampling times. These snapshots show varying degrees of qualitative agreement with the dynamics captured in the training data. In particular, the simulation of the CHIR 2–3 condition shows a movement of cells from the left branch to the right, in agreement with the training data that capture the movement of EPI cells to a CE state. The simulation of the NO CHIR condition fails to capture a clear transition of cells from the EPI to the AN state, primarily due to the overlap of the regions corresponding to these cell types. There is, however, a marked increase in the density of simulated cells in the upper-left region of the landscape, which may be interpreted as an increase in the AN population.

During the training process we monitor the average loss computed across the validation data, consisting of four experimental conditions. This assessment takes place at the end of each epoch, and equally weights all datapoints, each corresponding to a pair of consecutive timepoints. The validation loss history is depicted in Fig. 7b. While the model appears to converge, the loss does not converge towards 0, suggesting that elements of the training data are inconsistent with the model. We expect that these inconsistencies are primarily the result of our use of a linear approximation to the decision manifold, and also underfitting the NO CHIR experimental condition. This artifact is likely a result of insufficient data normalization, and the fact that the NO CHIR condition is under-represented in the data, compared to signaling timecourses that include an initial CHIR signal.

We withhold two of the eleven experimental conditions from the training procedure as an evaluation dataset, and use these as an out-of-sample test of model performance (Table 1). Our approach to model evaluation assesses the ability of the model to generate a feasible ensemble state at time *t*_*i*+1_ from an initial condition at time *t*_*i*_, where *t*_*i*_ and *t*_*i*+1_ are consecutive sampling times. An alternate approach assesses the ability of the model to reconstruct the full time course of an experimental condition. The distinction between these two evaluation methods is important, as the first serves as a direct evaluation of the computational training procedure on out-of-sample data, while the latter is likely of greater value for practical purposes, for example if one is interested in predicting the complete trajectory of an ensemble under the influence of a particular signal profile.

Fig. 8a shows ensemble trajectories for the two evaluation conditions, CHIR 2–4 and CHIR 2–5 FGF 2–4. It is important to note that the signal profiles for these conditions agree up to D3, and that the data for D2-3 consist of the same cells. This accounts for the similarity in the average loss between the first two transition intervals depicted in Fig. 8b. Moreover, this data is seen during the training process, due to overlap between the signaling conditions making up the training data. Thus, the true test of generalization on held-out data is the performance of the model at later intervals, and alternative approaches to assessing generalization error deserve further investigation.

**Figure 8.**
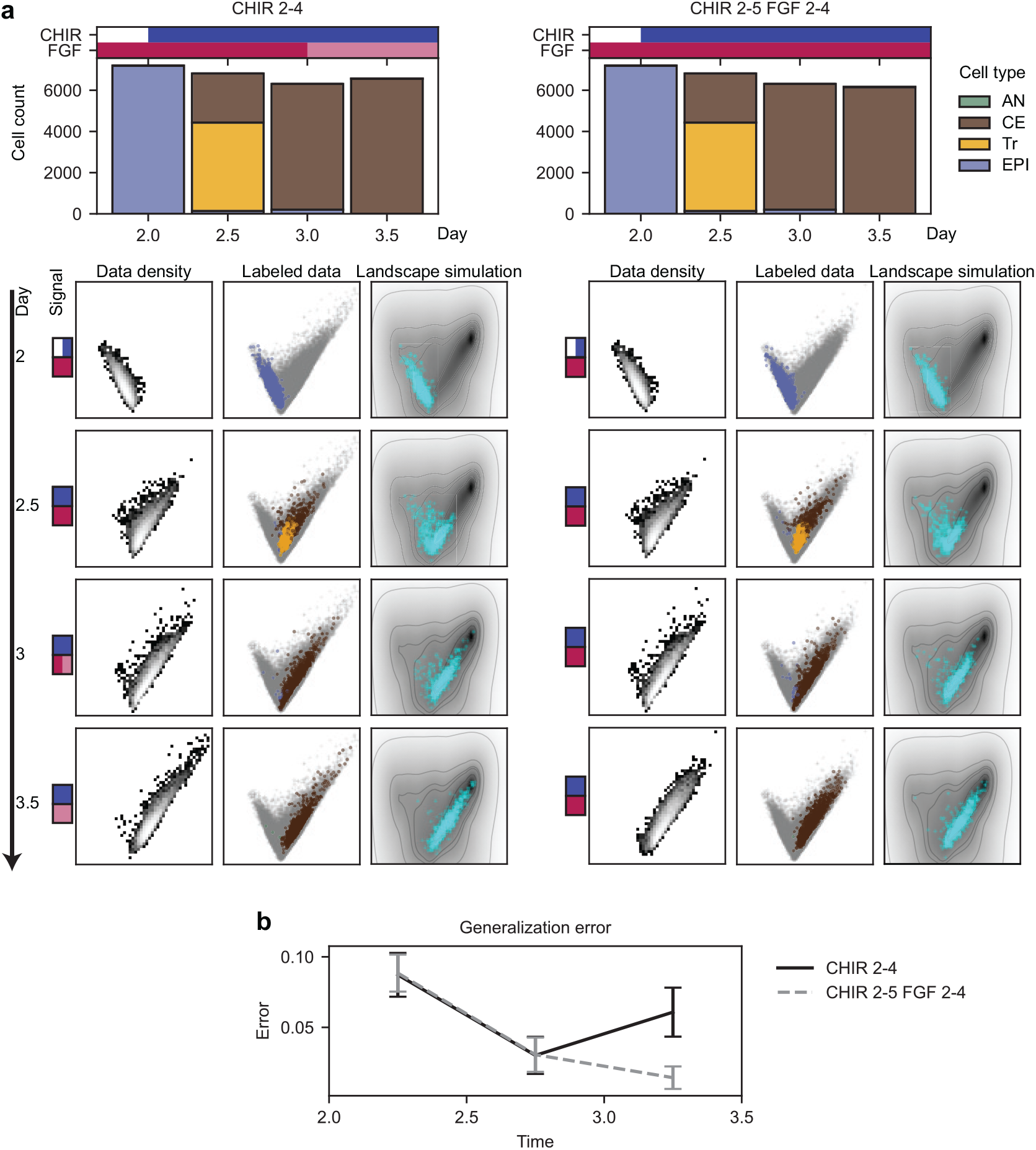
Evaluation of a trained PLNN model on a held out set of experimental conditions. **(a)** Experimental conditions held out of the training process and used for model evaluation. Plots are analogous to those depicted in Fig. 7a. **(b)** The generalization error corresponding to each held out signal and sampling interval, computed over multiple initial conditions and simulations, as described in the text. This average is plotted with error bars showing two standard deviations. Importantly, the data corresponding to the initial time intervals are seen during the training process, even though the signaling conditions are nominally held out. This is again due to overlap in the signal profiles making up the training and evaluation datasets, with some signals deviating only over the final interval.

## Discussion

We set out to leverage the universal function capacity of neural networks in order to infer a Waddington-like landscape model of cellular differentiation dynamics. Previous work has established the value of low-dimensional, geometric models, including their predictive capabilities, but has relied on abstract normal forms as the basis of their construction [40]. While the use of such normal forms is justified by a rigorous, mathematical theory [39], the abstract state variables appearing in these forms lack biological meaning, and obscure a direct connection to the molecular profile of a cell. Our physically-inspired machine learning approach bridges the intuitive, phenotypic understanding of cellular differentiation as depicted by Waddington, with the complex, molecular view wherein cells transition between states in high-dimensional gene space. A practical consequence of our modular framework is that each of the components—static landscape, signal processing, and noise—can be made more complex while not altering the overarching construction. Below, we discuss a number of possible modifications that may better capture biological realities. Ultimately, we return to the question of development as it takes place in the embryo, where spatial organization plays a pivotal role in the signaling dynamics driving cell fate decisions.

While the arsenal of machine learning architectures has expanded dramatically in the last decade, we chose to use a simple, feedforward neural network in a single module of our computational framework, namely the underlying static potential **Φ**. This module indirectly yields the governing dynamics of the system, via its gradient. One can imagine an alternative approach that attempts to learn directly the vector field governing cellular dynamics in a low-dimensional space. However, this approach would no longer be guaranteed to produce an interpretable picture of the dynamics as taking place in a Waddington-like landscape, as generic vector fields cannot be written as the gradient of a potential. Our understanding of the phenomenon—that it should be hierarchical and absent any recurrent flow—allows us to make the gradient assumption. Then, the inference of the scalar potential implicitly constrains the dynamics.

While the potential module **Φ** is given a great deal of flexibility as a neural network, the signal processing module **Ψ** and the noise module **N** are in contrast heavily constrained in their functional forms. The linear tilt effect evidenced by the term ***x***^*T*^ **Ψ**(***s***) in (8) is inspired by the normal forms enumerated in Rand et al. [39],

sufficient to describe biologically relevant transitions. This ansatz also gives a concrete, physical meaning to the latent variables that are the landscape coordinates. That the signal processing function **Ψ** is a simple linear transformation is a separate, simplifying assumption, one that has the effect of passing all nonlinearities to the landscape module **Φ**. In Sáez et al. [40], the authors make a similar assumption, expressing the linear coefficients in a parameterized landscape model as an affine transformation of signal levels. In addition, they introduce a nonlinearity in the way that a signal is processed by a cell by including a memory effect. In cases in which a memory or other nonlinear effect is essential, it is feasible to introduce additional model parameters as part of the functional form of **Ψ** without much additional difficulty. One could even take the extreme approach of parameterizing this module using an additional neural network. We emphasize that as long as the signal processing function does not depend on the state ***x***, its effect on the landscape will always be linear, in accordance with (8).

Similarly, the noise module **N**—which we assume to yield a constant diagonal matrix, i.e. homogenous, isotropic noise—can easily be relaxed to allow for more general stochastic effects. A relaxation to permit non-isotropic noise is a trivial computational adjustment, although the impact that this would have in the model training process is unclear, and remains an open route of future investigation.

One drawback of our current framework is that we do not have a way of assessing uncertainty in the inferred system. This deficiency is readily apparent in the anomalous landscape minima that appear in regions never occupied by cells. While the landscape may be consistent across all of the signal profiles seen during training, these minima may incorrectly suggest that cells can be driven to occupy regions of state space that are actually inaccessible. At present, we rely on our knowledge of the domain over which cells have been observed, and treat any artifacts appearing outside of this domain as spurious.

We have focused on the inference of a two-dimensional potential function that tilts in response to two effective signals. In our synthetic experiments, the data have been two-dimensional, and no attempt is made to infer the manifold on which the governing dynamics are given by the gradient of a potential. Rather, we take for granted that the data sit in Euclidean space, and infer the parameterized landscape directly, as a potential defined over this flat space. In our application to real data, the treatment of the underlying decision manifold is done prior to and in complete isolation from the subsequent dynamical inference. We first identify a two-dimensional manifold onto which we project the observed cells, and then use the resulting data as input to the training process. Moreover, we choose to identify a *linear* manifold, that is, a flat plane in gene expression space. Both the assumption of a strictly linear manifold, and the separation between its identification and the inference of the gradient system atop it, offer directions in which we can generalize our approach. We foresee the use of either more advanced methods of nonlinear manifold inference or the application of an autoencoder architecture to learn a low-dimensional representation of the data, directly from the high-dimensional input.

Our framework involves a number of critical assumptions when it comes to the role of signaling. Some of these assumptions stem from the particular context in which we expect the use of our framework to be most immediately applicable, namely experimental settings in which a signaling timecourse is prescribed, and all cells are subject to the same, uniform application of that signal. In this setting, signaling—or at least a subset of the relevant signaling—is assumed to be controllable; an experimenter can specify a particular signaling timecourse and observe the ensuing cell state transitions. This situates us well outside the realm of *in vivo* development, where exact signaling dynamics are largely unknown, let alone controllable. In the real developing embryo, the signals secreted by a cell inform the differentiation process of other, nearby cells and depend on the particular state of the secreting cell. The resulting feedback between cell state and signaling results in a far more complicated system from the one our model suggests. How, then, can we expect a model such as we have described to be informative in the broader context of real development? One missing piece is the role that space plays in directing development, through the spatial organization of signaling patterns. We foresee future directions that take into account the spatial organization of tissues, and allow for heterogeneous signaling across cell populations.

## Methods

### A neural network architecture for the potential module

We use a consistent architecture for the potential module **Φ** across all of the trained landscape models referenced herein. For the fully-connected neural network constituting **Φ**^*nn*^, we use four hidden layers consisting of 16, 32, 32, and 16 neurons, respectively, with the final hidden layer followed by a fully connected layer to a single neuron. After each hidden layer, we apply the smooth softplus activation function, given by *α*(*x*) = ln(1+*e*^*x*^). No activation is applied to the scalar output of the final layer. For the confinement factor we set *C*_conf_ = 0.1, so that the confinement term Φ_0_(***x***) is a shallow quartic potential. We randomly initialize the weights of both **Φ**^*nn*^ and the tilt module **Ψ** according to a Xavier uniform distribution [47]. The biases of **Φ**^*nn*^ are initialized at 0. Additional details can be found in Supplementary Materials S2.

### Generating synthetic data

We use the algebraic form of the binary choice landscape (15) to generate an initial synthetic dataset on which we train the PLNN shown in Fig. 4. We use the Euler-Maruyama method with a step size of Δ*t* = 0.001 to simulate *N*_exps_ = 100 experiments in which *N*_cells_ = 500 cells evolve in the landscape over an interval of time 0 ≤ *t* ≤ *T* = 100, under the influence of a sigmoidal signal profile. We capture snapshots of the ensemble at intervals of Δ*T* = 10, including the initial and final states, thereby yielding 10 training datapoints of the form (*t*_0_, *X*_0_; *t*_1_, *X*_1_; ***s***(*t*)) per experiment. We set *σ*^∗^ = 0.1, the magnitude of isotropic, homogeneous noise in the system. We take for the signal processing function the identity map, so that the signals *s*_1_ and *s*_2_ are identified with the tilt parameters *τ*_1_ and *τ*_2_, respectively.

For each experiment, a sigmoidal signal profile is drawn from a prior distribution of signal functions, thereby allowing us to restrict the domain of allowable signals. Each two-dimensional signal profile ***s***(*t*) = (*s*_1_(*t*), *s*_2_(*t*)) is of the form

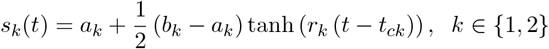

where *a*_*k*_, *b*_*k*_, *r*_*k*_, and *t*_*ck*_ are parameters drawn from specified distributions. *a*_*k*_ and *b*_*k*_ are the asymptotic initial and final values of signal *k*, respectively; *r*_*k*_ is the rate at which the signal transitions; and *t*_*ck*_ is the critical time at which the signal is halfway between its extreme values. The priors for *a*_*k*_, *b*_*k*_, *r*_*k*_, and *t*_*ck*_ (Table S13) are such that the central attractor bifurcates in nearly all experiments, while the peripheral attractors typically persist. This captures the phenomenon in which cells initially located at the central attractor proceed to move towards a peripheral one after it vanishes.

In each experiment all cells are initialized at the same point in the landscape, near the known location of the central attractor. The simulation then runs for a short burn-in duration *T*_*burn*_ = 0.1*T* under a given signal, ***s***_*burn*_, allowing for the randomization of the ensemble. The state of the ensemble at the end of this burn-in phase is taken as the initial condition of the ensemble at time *t* = 0.

We generated an analogous dataset using the binary flip landscape. The datasets used for the sampling rate analysis (Fig. 5a) were also generated similarly. Details can be found in Supplementary Materials S4.

### Training a PLNN

The training procedure for a PLNN involves simulating the forward evolution of an ensemble of cells from an initial state, and comparing the resulting *simulated* final state to an *observed* final state. A schematic of this process is illustrated in Fig. 3d. The observed training data are of the form

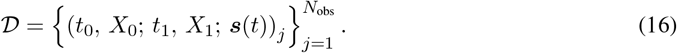

That is, each datapoint consists of an initial ensemble state *X*_0_ observed at time *t*_0_, a corresponding final state *X*_1_ at time *t*_1_, and the signal profile ***s***(*t*) used over the course of the corresponding experiment. Here, *X*_0_ and *X*_1_ are data matrices, with each row a *d*-dimensional cell. The number of rows in each observed data matrix may vary, as the number of observed cells at each timepoint and in each experiment may change. Moreover, the timepoints *t*_0_ and *t*_1_ need not be consecutive sampling times. For example, if in an experiment observations {*X*_1_, *X*_2_, *X*_3_} were made at times *t* ∈ {1, 2, 3}, then a datapoint (1, *X*_1_; 3, *X*_3_; ***s***(*t*)) is entirely valid. However, for the sake of computational efficiency it may be advantageous for all datapoints to consist of equally spaced observations.

Given a datapoint 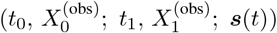 from the observed dataset, we must simulate a corresponding initial condition 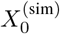 forward in time to obtain a simulated final state 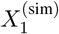.In order to take advantage of the accelerated linear algebra tools provided by JAX [48], we fix the number of simulated cells, 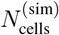, as a model hyperparameter, so that the simulated data matrices have consistent dimensions. In the case that the observed data matrices do share the same number of cells, we simply set 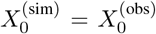, provided that the number of observed cells does not exceed a memory constraint. In the case that the number of cells is too large, or in the case that the observed data matrices are of inconsistent dimension, we construct the simulated initial condition by randomly sampling rows from the observed data matrix:

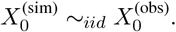

Using the PLNN architecture described above, we simulate the ensemble of cells 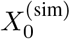 forward in time, constituting a sample trajectory of the system of SDEs in (14). This step of the training process is illustrated in Fig. 3c. The python package `Diffrax` [49] provides a number of (stochastic) differential equation solvers. We use Heun’s method, a second order explicit Runge–Kutta method, which provides sufficient accuracy while maintaining computational efficiency [50]. We must specify as a hyperparameter a timestep Δ*t*. In order to balance computational efficiency and accuracy, we vary this hyperparameter over the course of training, starting with a large timestep and periodically reducing it after a specified number of epochs.

We then compare the resulting simulated state 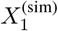 at time *t*_1_ *> t*_0_ to the observed final state 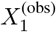 (or a subset of this matrix, if necessary) and compute the value of the specified loss function 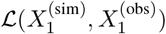.

As we compare the simulated and observed final states in a distributional sense, the dimensions of the matrices 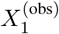 and 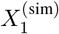 need not be consistent. Once the loss is computed, we update the model parameters using gradient updates, as implemented in the optimization package `Optax` [51] within the JAX ecosystem. We use the RMSProp algorithm [52] with an exponentially decaying learning rate to perform parameter updates. The values of all relevant hyperparameters are provided in Supplementary Materials S4.

### Choice of loss function

We require a loss function that compares observed and simulated ensemble states 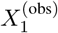 and 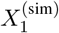 in a distributional sense. To this end, we experimented with two loss functions. The first is a numerical estimate of the KL divergence [53]. The second is based on the maximum mean discrepancy (MMD) which has been used in the context of the two-sample test [44, 54]. Additional mathematical details of both methods can be found in Supplementary Materials S3.

Ultimately, we settled on using an MMD-based loss function, as this provided a differentiable function through which we could effectively backpropagate during the training process. While training using the KL divergence estimate as a loss function did appear to converge—resulting in models similar to those found using the MMD loss—the training process was slower, and we hypothesize that this was a result of the use of a nearest neighbor computation in the calculation of the estimate.

As introduced in Gretton et al. [44], an unbiased empirical estimate of the squared population MMD is given by

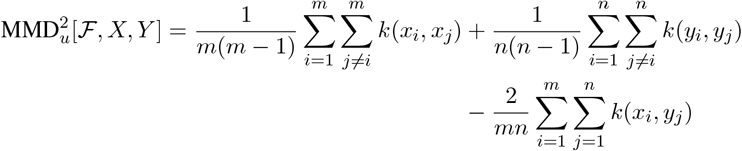

where observations *X* = {*x*_1_, …, *x*_*m*_ }and *Y* = {*y*_1_, …, *y*_*n*_ }are independently and identically distributed from distributions *p* and *q*, respectively, and ℱis a family of functions that we will take to be the unit ball in a reproducing kernel Hilbert space (RKHS) ℋwith characteristic kernel *k*. For our purposes, it suffices to say that Gaussian kernels are characteristic on ℝ^*d*^ [55], and for a characteristic kernel the MMD is a metric on the space of probability distributions. Thus, minimizing the MMD results in convergence of the distributions *p* and *q*, the observed and simulated data generating processes.

We use for a kernel a function of the form

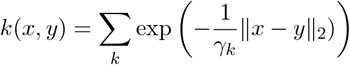

where the *γ*_*k*_ constitute a set of one or more bandwidth parameters. In practice, the median pairwise distance between sampled points, i.e. cells, has been shown to be a suitable choice for this bandwidth parameter [56].

### Cell type labeling for *in vitro* data

The cell type labels assigned to the observed cells in the *in vitro* mESC dataset, and appearing in Figs. 7 and 8 are determined using the procedure originally described in Sáez et al. [40]. We recapitulate this labeling and refer the reader to Supplementary Materials S5 and the original work for more details. An important detail, as it pertains to the following section, is the use of Gaussian mixture models (GMMs) in the cell type identification process. The algorithm used by Sáez et al. [40] associates to each time point *t* a GMM _𝒢*t*_ consisting of multiple components, with each component corresponding to a cell type. As part of our recapitulation of the cell type labeling, we construct and retain these GMMs in order to use them in the isolation process.

### Isolation of *in vitro* binary decision data

We use the cell type labels assigned to each cell in the *in vitro* mESC dataset to isolate those cells relevant to the first binary decision, involving the transition of EPI cells to either the AN or CE state. For each timepoint *t*, we remove all cells labeled M or PN, replacing these cells with “pseudo-cells” generated from the GMM 𝒢_*t*_ retained from the cell type labeling process. Specifically, we use the component(s) of the GMM associated with the CE state to generate random samples from a multivariate normal distribution. This allows us to replace the PN and M cells, which have presumably transitioned out of the CE state, with cells that are distributed around a CE attractor as determined by the labeling algorithm. This replacement strategy is necessary in order to respect the relative proportion of cells that have entered the CE attractor. In reality, this state is an intermediate one, and cells may escape it. Our approximation treats this state as a terminal one—a periphery in a binary decision.

### PLNN training on *in vitro* data

The model shown in Fig. 7 was trained on the processed mESC dataset. The model was trained for a maximum of 1000 epochs with early stopping implemented if the model did not improve for 200 epochs (with respect to the validation loss). Due to the large and varying number of cells captured at each timepoint (approximately 7200 in each case), we implemented a sampling step in the training process. Given a datapoint of the form (*t*_0_, *X*_0_; *t*_1_, *X*_1_; ***s***(*t*)), we randomly sampled 800 cells from the rows of *X*_0_ to yield the simulated initial condition 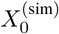.Each epoch then consisted of 10 passes through the training data, in mini-batches consisting of 50 training datapoints.

The neural network architecture was the same as for the synthetic landscape experiments, detailed above. We used Heun’s method as our choice of SDE solver, with an initial timestep of 0.005. This timestep was reduced periodically by a factor of 0.5, after 50, 100, 200, and 300 epochs. For the MMD loss function, we used a Gaussian kernel with a single, precomputed bandwidth parameter *γ ≈*1.355, the median pairwise distance between all observed cells at a given timepoint, averaged across timepoints.

## Code availability

A repository with code implementing the parameterized landscape architecture and training process, written in python, is available at https://github.com/AddisonHowe/plnn. A separate repository, illustrating an application of this architecture and including scripts to generate all figure panels, is available at https://github.com/AddisonHowe/dynamical-landscape-inference. Code recapitulating the cell type labeling process originally detailed in Sáez et al. [40], and isolating the data relevant to the first binary decision, can be found at https://github.com/AddisonHowe/mescs-invitro-facs. Original data from the *in vitro* mESC experiments performed by Sáez et al. [40] was requested from and provided by the authors.

## Acknowledgments

M.M. and A.H. thank Meritxell Sáez and Elena Camacho-Aguilar for providing data, and Eric Siggia for his guidance and ideas. A.H. also thanks Dominic Skinner and Connor Puritz for their feedback and critiques.

This research was supported in part by grants from the NSF (DMS-2235451) and Simons Foundation (MP-TMPS-00005320) to the NSF-Simons National Institute for Theory and Mathematics in Biology (NITMB).

M.M. and A.H. were supported by the NSF-Simons Center for Quantitative Biology at Northwestern University (DMS-1764421) and the Simons Foundation (597491-RWC). M.M. is a Simons Investigator.

A.H. was also supported by the NSF-RTG Interdisciplinary Training in Quantitative Biological Modeling (DMS-1547394). This project has been made possible in part by the Chan Zuckerberg Initiative DAF (DAF2023-329587), an advised fund of the Silicon Valley Community Foundation. This research was supported in part by grants from the NSF (PHY-1748958) and the Gordon and Betty Moore Foundation (2919.02) to the Kavli Institute for Theoretical Physics (M.M.).

This research was supported in part through the computational resources and staff contributions provided by the Genomics Compute Cluster which is jointly supported by the Feinberg School of Medicine, the Center for Genetic Medicine, and Feinberg’s Department of Biochemistry and Molecular Genetics, the Office of the Provost, the Office for Research, and Northwestern Information Technology. The Genomics Compute Cluster is part of Quest, Northwestern University’s high performance computing facility, with the purpose to advance research in genomics. We thank J. Milhans, A. Kinaci, S. Coughlin and all members of the Research Computing and Data Services team at Northwestern for their support.

