## Supplemental Materials for "Learning geometric models for developmental dynamics"

Supplementary Materials

Addison Howe*[1] and Madhav Mani*[1]

[1]Department of Engineering Sciences and Applied Mathematics,
Northwestern University,
Evanston, IL, 60208, USA

December 11, 2024

**SA Tracing fold curves** **41**

**References** **42**

### Introduction

This supplement contains additional information, including a complete description of the architectures and numerical procedures introduced in the main text. In Section S1 we discuss in detail the mathematical theory underpinning parameterized landscapes and their use in modeling cellular differentiation dynamics. In Section S2 we provide a full description of the Parameterized Landscape Neural Network (PLNN) model architecture introduced in the main text. Section S3 details the two loss functions we use for model training. In Section S4 we perform a number of *in silico* experiments, assessing the performance of the model in a variety of contexts. Section S5 details our application to an *in vitro* dataset, and includes a full recapitulation of the cell type labeling algorithm performed in the original work [1].

The following videos, referenced below, can be found at https://github.com/AddisonHowe/dynamical-landscape-inference.

1. Movie 1: Data generation for the binary choice landscape.
2. Movie 2: Data generation for the binary flip landscape.

### S1 Landscapes as models of cellular differentiation

The approach we take to modeling dynamics of cellular differentiation has as its original source of inspiration Waddington's metaphor of the epigenetic landscape [2]. In this metaphor, a ball representing a developing cell rolls downhill through a hilly landscape, with the valleys of the landscape representing different cell states. Points at which a valley diverges into two, or appears suddenly, signify points at which a cell chooses between two or more fates. The basins at the bottom of the landscape correspond to the ultimate fates a cell may take. The biological notions of commitment and canalization—that the outcome of a developmental process is discrete and robust to perturbations—are captured in the hills separating the different valleys, and the notion that once a cell has descended sufficiently far in a particular valley it is unlikely to jump into a neighboring one [3, 4].

The landscape metaphor, and the biological ideas that it encapsulates, can all be formalized using the language of dynamical systems theory, a branch of mathematics that has long been understood as appropriate to a rigorous formulation of developmental biology, especially in the context of cellular differentiation [5, 6]. At the basis of this formalization is the representation of a cell as a vector of the relevant molecular components that define its state:

$$\boldsymbol{x} = [g_1, g_2, \dots, g_n]^T.$$

Then, a system of differential equations

$$\frac{dg_i}{dt} = f_i(g_1, \dots, g_n) \tag{S1}$$

defines the dynamics by specifying the interactions between the molecular components. In Waddington's metaphor, the shape of the landscape is understood to be determined by a complex network of genes that provide an underlying scaffolding, and thus the relevant molecular components in the metaphor are the set of genes and gene products. Decades of work in developmental genetics, combined with the modern tools of cell profiling, have provided us a list of these relevant components [7]. Now, in the present era of single-cell RNA sequencing (scRNAseq), we are capable of unprecedented levels of insight into the transcriptomic makeup of a cell, and it is natural to represent a cell by its transcriptome, a vector detailing its expression of mRNA.

Given the tools available, a common approach to understanding the dynamics of development attempts to determine the equations on the right hand side of (S1), often for specific, well-studied chemical pathways. However, these pathways often involve numerous components, and the differential equations used to model the kinetics—typically Hill or Michaelis–Menten forms—grossly simplify the true biology, and involve a large number of parameters whose values are often unknown and difficult to determine [7, 8]. These deficiencies call for an alternative approach, one that focuses on the simple, emergent phenomenon—as depicted in Waddington's metaphor—that arises out of the complex set of governing equations.

To this end, a body of work has been pioneered that grounds the biological ideas of Waddington's landscape in the mathematics of dynamical systems theory, focusing on the emergent phenomenon rather than the governing equations at the molecular level [1, 9–20]. A review of this geometric perspective in the context of developmental biology is provided in Raju and Siggia [7]. To use the Waddington metaphor, this approach directly concerns itself with the low-dimensional dynamics defined by the landscape, rather than the complex gene regulatory dynamics that underlie it. One attempts to find a simple set of equations that recapitulate the observed phenomenon, for example the particular phenotypic progression of a cell over the course of differentiation. Just as Waddington's landscape offers a simple, low-dimensional representation of the phenomenon of differentiation—in actuality defined by an incredibly complicated and high-dimensional set of gene interactions—the geometric perspective emphasizes the value of a low-dimensional potential function encapsulating the relevant dynamics.

#### S1A Mathematical theory of parameterized landscapes

We express a parameterized landscape as a potential function $\phi(\boldsymbol{x};\boldsymbol{p})$ defined over a phase space $\Omega \subseteq \mathbb{R}^d$, with $\boldsymbol{x} \in \Omega$, and parameterized by a vector $\boldsymbol{p} \in \Gamma \subseteq \mathbb{R}^r$. A parameterized landscape of this form induces a vector field, or flow, $\boldsymbol{F}$, according to

$$\boldsymbol{F}(\boldsymbol{x}, t) = -\nabla_{\boldsymbol{x}} \phi(\boldsymbol{x};\boldsymbol{p}). \tag{S2}$$

We will typically consider the case where the parameter governing the shape of the landscape changes in time, and write $\boldsymbol{p} = \boldsymbol{p}(t)$.

The work of Sáez et al. [1] offers a motivating example. They construct a two-dimensional dynamical system using two polynomial forms, $\phi_{bc} = \phi_{bc}(x, y; p_1, p_2)$ and $\phi_{bf} = \phi_{bf}(x, y; p_3, p_4)$, each changing in response to two parameters. These forms represent two of the classes enumerated by Rand et al. [19], and are termed the *binary choice* and *binary flip*, respectively. Each is given by a simple polynomial equation, and is archetypal in the sense that it corresponds to a particular class of 3-attractor systems, which exhibit a particular global bifurcation structure. By algebraically stitching together these forms so that each governs the dynamics in one half of the plane, Sáez et al. construct a five-attractor system that changes shape in response to the full parameter vector[2] $\boldsymbol{p} = (p_1, p_2, p_3, p_4)$ and undergoes bifurcations in a fully-specified manner, at specific values of $(p_1, p_2)$ and $(p_3, p_4)$. Then, they express the landscape parameters as an affine function of two signals, $s_1$ and $s_2$, according to

$$\boldsymbol{p} = \boldsymbol{w}_0 + \boldsymbol{w}_1 s_1 + \boldsymbol{w}_2 s_2, \tag{S3}$$

with $\boldsymbol{w}_0, \boldsymbol{w}_1, \boldsymbol{w}_2 \in \mathbb{R}^4$ vectors of weights. Given $\boldsymbol{w}_0$, $\boldsymbol{w}_1$, and $\boldsymbol{w}_2$, the weights define a transformation from signals to the landscape parameters $\boldsymbol{p}$, and as the signals change, so too does the landscape. In this way, the signals are understood as the true controls driving the change in shape of the overall landscape, but with the change constrained by the specified forms. Inference of the weights is informative, as they shed light on how particular combinations of signals affect the system.

Critically, each landscape form, $\phi_{bc}$ and $\phi_{bf}$, depends on exactly two of the four parameters, and does so in a linear fashion, so that they effectively "tilt" the landscape. We call such systems *tiltable landscapes*. A tiltable landscape $\phi(\boldsymbol{x};\boldsymbol{p})$ is a parameterized landscape that can be written in the form

$$\phi(\boldsymbol{x};\boldsymbol{p}) = \tilde{\phi}(\boldsymbol{x}) + \boldsymbol{x}^T \mathbf{L} \boldsymbol{p} \tag{S4}$$

where $\mathbf{L} \in \mathbb{R}^{d \times r}$ is a constant matrix. Thus, the effect of varying $\boldsymbol{p}_i$ is to "tilt" the underlying *static landscape*, $\tilde{\phi}$, containing the essential nonlinear features of the landscape, linearly in the direction of the $i$th column of $\mathbf{L}$. The particular form of $\tilde{\phi}$ determines what kinds of bifurcations may occur as the landscape tilts. Without loss of generality, we consider the case where the number of parameters is given by the dimension $d$ of the landscape, with $\mathbf{L}$ the identity matrix. In this case we denote the landscape parameters as $\boldsymbol{\tau} \in \mathbb{R}^d$, and write the tiltable landscape in the form

$$\phi(\boldsymbol{x};\boldsymbol{\tau}) = \tilde{\phi}(\boldsymbol{x}) + \boldsymbol{x}^T \boldsymbol{\tau}. \tag{S5}$$

These systems—in the way that they represent an underlying landscape that tilts in response to exogenous signals—motivate our approach, and also serve as synthetic testing grounds for our model. we use the landscapes to generate synthetic, two-dimensional data, to which a model can be fit.

#### S1B The binary choice landscape

The first landscape we consider facilitates an "all-or-nothing" binary choice. The governing equation of this landscape is given by

$$\phi_{bc}(x, y; \tau_1, \tau_2) = x^4 + y^4 + y^3 - 4x^2 y + y^2 + \tau_1 x + \tau_2 y. \tag{S6}$$

The bifurcation curves corresponding to $\phi_{bc}$ are shown in Fig. S1, each colored in accordance with the particular bifurcating fixed point it represents. On these curves in parameter space, a fold, or saddle-node,

[2] In the original work, there are a number of additional parameters that we ignore here for the sake of simplicity. We refer the reader to Sáez et al. [1].

bifurcation takes place, whereby a stable fixed point and unstable saddle either coalesce and vanish, or appear together suddenly. Of particular note is the bifurcation of the central, blue attractor. Crossing the blue bifurcation curve—moving downwards from the bounded, central region of parameter space in which all three attractors are present—results in the disappearance of the central attractor. There is then a subsequent flow from the vicinity of the vanished attractor to whichever peripheral fixed point was connected, by an unstable manifold, to the intermediate, vanishing saddle. For example, moving from the labeled point 0 to point 3 would result in any cells located near the blue attractor to move to the red attractor, as both the blue attractor and intermediate saddle vanish in the bifurcation.

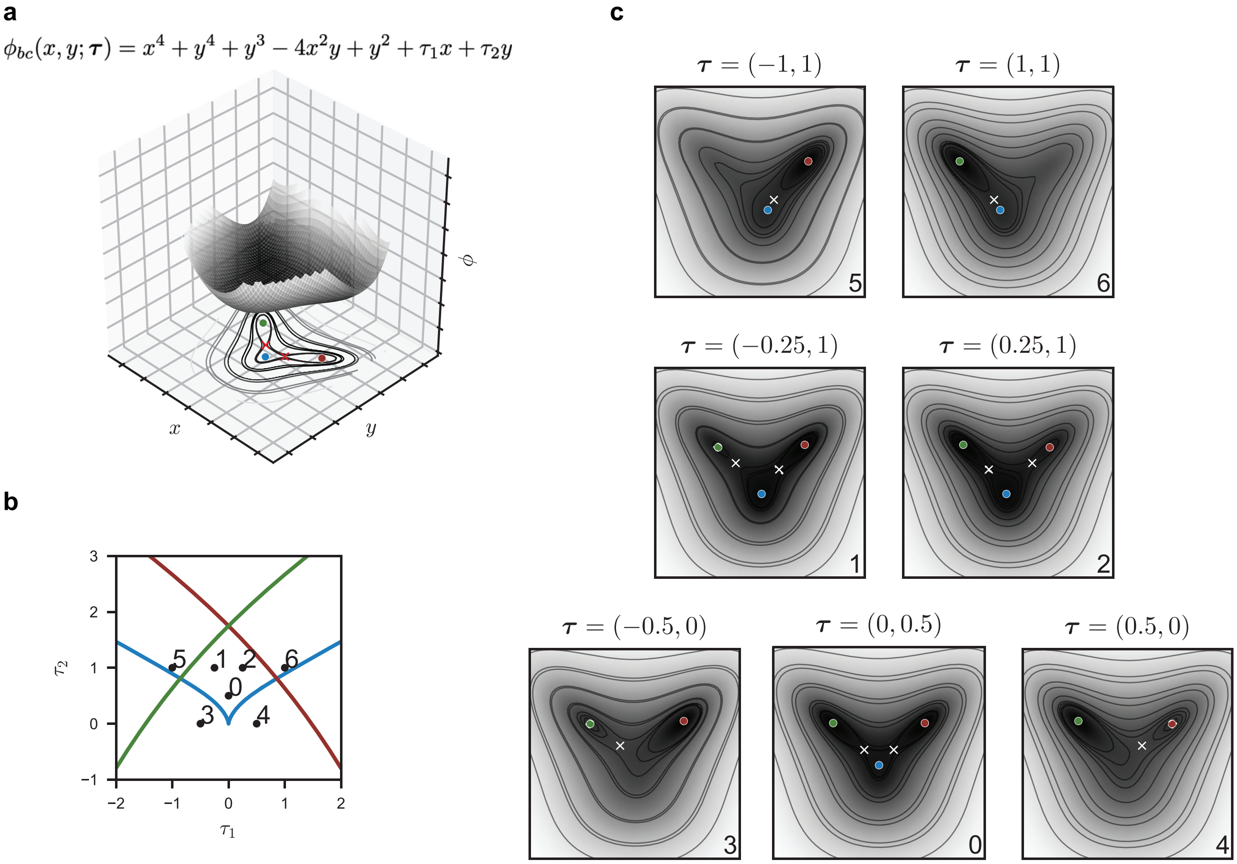

Figure S1: The binary choice landscape.

### S1C The binary flip landscape

The second landscape we consider is the binary flip landscape, given by

$$\phi_{bf}(x, y; \tau_1, \tau_2) = x^4 + y^4 + x^3 - 2xy^2 - x^2 + \tau_1 x + \tau_2 y. \tag{S7}$$

The bifurcation curves corresponding to $\phi_{bf}$ are shown in Fig. S2. In this case, in addition to saddle-node bifurcations, changing $\tau_1$ and $\tau_2$ can result in a heteroclinic flip bifurcation, denoted by dashed purple lines. In a heteroclinic flip bifurcation, no fixed points vanish, but the unstable manifold of a central saddle point "flips" from one peripheral attractor to another. At the exact point of the bifurcation, the unstable manifold connects the saddle to another saddle, so that there is a heteroclinic orbit between two saddles. The effect of this, in the stochastic dynamical systems context, is to favor flow from the central attractor towards one peripheral attractor over the other, while maintaining the presence of both. In addition, provided a sufficient degree of noise in the system, cells can move between the two peripheral attractors, without passing through the vicinity of the central attractor.

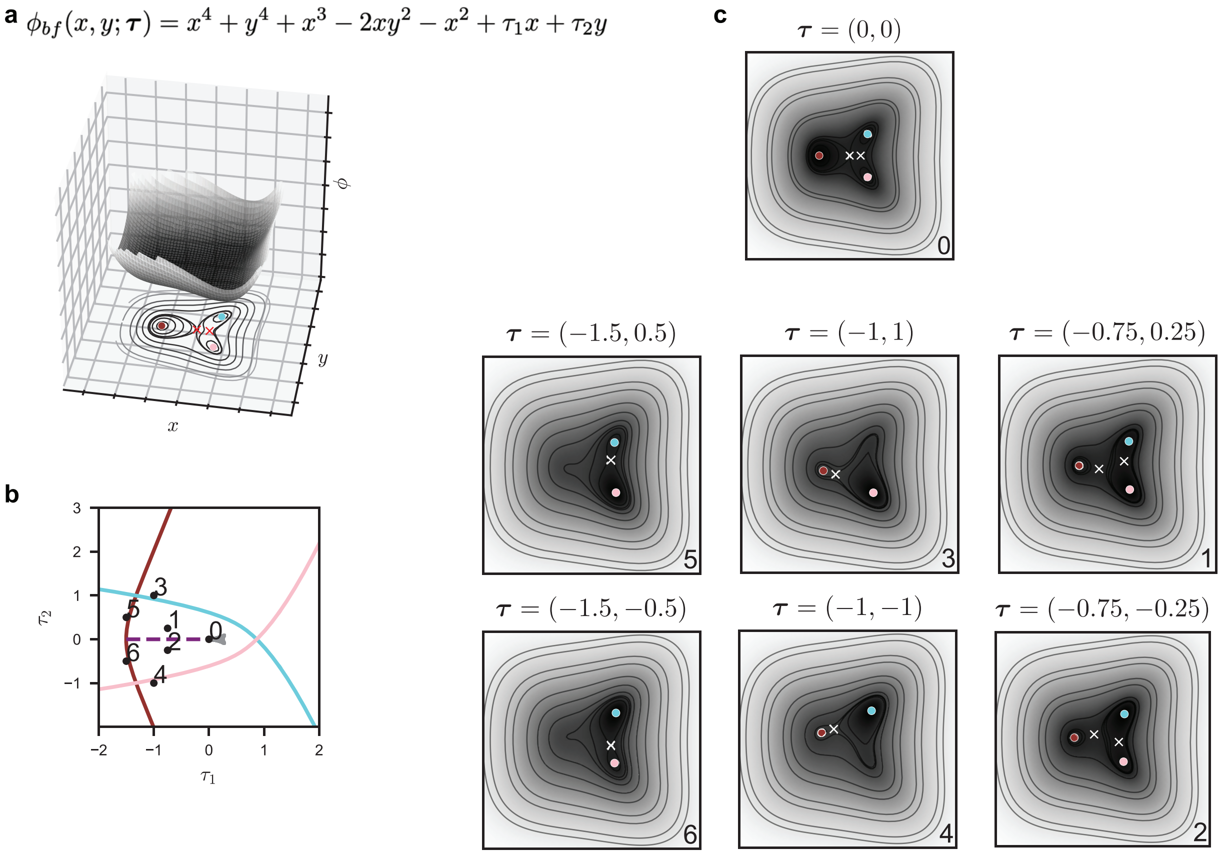

Figure S2: The binary flip landscape.

### S1D Morse–Smale systems and non-gradient behaviors

Our modeling framework assumes gradient dynamics, induced by the inferred potential. Such conservative vector fields constitute a highly-restricted class of dynamical systems, and do not capture non-gradient behaviors such as rotational flows. In the context of development, the assumption of gradient dynamics may not be unreasonable, as many examples of cellular differentiation lack any notion of recurrence or cyclical behavior. That being said, such cases are not outside of the realm of possibility, and if nothing else pose interesting academic questions. The theory presented by Rand et al. [19] makes room for dynamical behavior beyond the strictly gradient form. Rand et al. consider Morse–Smale systems, which encompass a class of dynamical systems that are suitable to represent the biological systems of interest [21].

Rand et al. restrict their consideration to compact systems with a finite number of fixed points and no periodic orbits. These restrictions are biologically justified. The developmental phenomena we aim to capture are those systems in which a cell transitions in an ordered manner between a number of discrete states until ultimately settling into a final, or terminal, state. This suggests a system with a hierarchical structure in which cells ultimately flow into one of a number of attractors, rather than continuously cycle between states. While periodic behavior, for example as exhibited in the cell cycle, may very well be observed in developmental processes, we view these dynamics as taking place in directions orthogonal to the space in which the cellular decision dynamics occur. In addition to limiting the allowable long-time dynamics, Rand et al. make use of the notion of *structural stability*. A dynamical system is structurally stable if it is robust to sufficiently small, smooth perturbations—that is, if such perturbations do not change its qualitative form. Biologically, structural stability is a natural assumption, asserting that the developmental system, defined by some underlying gene network, is itself robust to small perturbations.

Given these restrictions, a suitable class of dynamical systems to consider is the class of *gradient-like Morse–Smale* systems. A Morse–Smale (MS) system is one for which, (i) there are a finite number of fixed points and periodic orbits, all of which are hyperbolic; (ii) all stable and unstable manifolds intersect transversely, if at all; and (iii) the non-wandering set consists of fixed points and periodic orbits alone [22].

These systems are particularly well-behaved, boasting structural stability in arbitrary dimensions, and lacking complex recurrent behavior. The additional notion of *gradient-like* asserts that the system has only fixed points, and no periodic orbits (a simpler form of recurrent behavior permitted in general by the MS definition).

A particularly useful feature of gradient-like MS systems is that they guarantee the existence of a Lyapunov (or potential) function $f : \Omega \to \mathbb{R}$, whose critical points coincide with the fixed points of the system, and which decreases along all trajectories. In this way, the Lyapunov function captures a notion of "downhill flow" in the system, justifying the landscape metaphor [19]. While the Lyapunov function provides a notion of height in the system, it is not sufficient to determine the full dynamics, as multiple flows may satisfy the conditions above. To pin down the particular dynamics, one must either specify the stable and unstable manifolds of the saddles in the system, or provide a Riemannian metric that, together with the potential, defines the dynamics *almost* everywhere, outside of arbitrarily small regions around fixed points.

This caveat is necessary because MS systems are *almost* gradient systems, in the following sense. A *gradient system* is a system defined by a potential function $f$ and a metric $g_{ij}$ according to

$$\dot{x}_i = -\sum_j g^{ij}\frac{\partial f}{\partial x_j}, \tag{S8}$$

with $(g^{ij}) = (g_{ij})^{-1}$. Like conservative vector fields defined by the gradient of a potential on the plane, the fixed points of a gradient system necessarily have real eigenvalues. MS systems, on the other hand, allow for non-gradient phenomena around fixed points, such as spiral sinks, a consequence of a fixed point with complex eigenvalues. For any MS system, one can find a potential and a metric that will produce the flow as defined in (S8), at all points in phase space except possibly around neighborhoods of each fixed point, which can be made arbitrarily small. In this way, using a suitable potential and metric, a gradient-like MS system can be expressed as a gradient system *almost* everywhere, with differences occurring in arbitrarily small regions.

To summarize, Morse–Smale theory guarantees that given a gradient-like MS system, there exists a gradient system defined by an appropriate potential function and metric, that will effectively describe the dynamics of interest. The focus of the present work is the determination of such a gradient system from time-lapse data capturing the dynamics of an appropriate developmental system. The model that we present assumes purely gradient dynamics, and leaves the simultaneous inference of a metric along with the potential as a route of future investigation.

### S2 A neural network architecture modeling landscape dynamics

A primary contribution of this work is an architecture and conceptual framework for inferring landscape dynamics from data. Below, we describe the components of this model, which we refer to as a Parameterized Landscape Neural Network (PLNN). A PLNN is intended to model a data generating process $\mathcal{M}^*$ in which an ensemble of $d$-dimensional cells, $\{\boldsymbol{x}^{(i)}\}_{i=1}^{N_{\text{cells}}}$ evolve over an interval of time $t \in [0, T]$. The dynamics governing this evolution are assumed to be defined by the gradient of an unknown potential function $\phi^*$ that "tilts" in response to a signal vector $\boldsymbol{s} \in \mathbb{R}^{n_{\text{sigs}}}$. The signal $\boldsymbol{s} = \boldsymbol{s}(t)$ changes in time, and we refer to such a function as a *signaling timecourse*, or *signal profile*. We assume that the time-varying potential function $\phi^*(\boldsymbol{x}; \boldsymbol{s}(t))$, parameterized by the signal $\boldsymbol{s}$, is a tiltable landscape

$$\phi^*(\boldsymbol{x}; \boldsymbol{s}(t)) = \tilde{\phi}^*(\boldsymbol{x}) + \boldsymbol{x}^T\psi^*(\boldsymbol{s}(t)) \tag{S9}$$

where $\tilde{\phi}^*(\boldsymbol{x})$ is a signal-independent, or *static*, underlying potential, and $\psi^* : \mathbb{R}^{n_{\text{sigs}}} \to \mathbb{R}^d$ is a function mapping the signal $\boldsymbol{s}$ to a vector $\boldsymbol{\tau}$ in the phase space $\mathbb{R}^d$. We refer to the function $\psi^*$ as a *signal processing function*. We refer to $\boldsymbol{\tau}$ as a *tilt vector*, and say that the signal *tilts* the landscape. At times, we may wish to consider the effects of tilting $\phi^*$ absent any discussion of signaling, and will then write $\phi^*$ as a function of $\boldsymbol{\tau}$:

$$\phi^*(\boldsymbol{x}, \boldsymbol{\tau}) = \tilde{\phi}^*(\boldsymbol{x}) + \boldsymbol{x}^T\boldsymbol{\tau}.$$

Now, the dynamics that we assume to govern the data generating process $\mathcal{M}^*$ can be written in the form of a stochastic differential equation (SDE). In general, this SDE takes the form

$$\begin{aligned} d\boldsymbol{x}^{(i)}(t) &= \boldsymbol{f}(t, \boldsymbol{x}^{(i)}(t))dt + \boldsymbol{g}(t, \boldsymbol{x}^{(i)}(t))dW^{(i)}(t) \\ \boldsymbol{f} &: \mathbb{R} \times \mathbb{R}^d \to \mathbb{R}^d \\ \boldsymbol{g} &: \mathbb{R} \times \mathbb{R}^d \to \mathbb{R}^{d \times d_w} \end{aligned} \tag{S10}$$

where $\boldsymbol{f}(t, \boldsymbol{x})$ and $\boldsymbol{g}(t, \boldsymbol{x})$ are the drift and diffusion terms, respectively, and $W$ is a $d_w$-dimensional Wiener process. Here, the drift is derived from the gradient of the potential according to

$$\boldsymbol{f}(t, \boldsymbol{x}(t)) = -\nabla_{\boldsymbol{x}}\phi^*(\boldsymbol{x}; \boldsymbol{s}(t)) = -\nabla_{\boldsymbol{x}}\tilde{\phi}^*(\boldsymbol{x}) - \psi^*(\boldsymbol{s}(t)). \tag{S11}$$

While in general the diffusion term $\boldsymbol{g}(t, \boldsymbol{x})$ is state-dependent, and allows for non-isotropic noise, we will assume that the noise in the system $\mathcal{M}^*$ is isotropic, governed by $d$-dimensional brownian motion, with a state-independent scale $\sigma^* \in \mathbb{R}$, so that we have the simple case

$$\boldsymbol{g}(t, \boldsymbol{x}(t)) = \sigma^* \cdot \boldsymbol{I}_d \tag{S12}$$

where $\boldsymbol{I}_d$ is the $d \times d$ identity matrix.

#### S2A Defining the PLNN

Having specified the assumed ground-truth data generating process $\mathcal{M}^*$, we are now ready to easily define the components of a general PLNN, which we denote by $\mathcal{M}$. Three modules make up the PLNN: a *potential module* $\boldsymbol{\Phi}$, a *tilt module* $\boldsymbol{\Psi}$, and a *noise module* $\mathbf{N}$. We can then represent the PLNN $\mathcal{M}$ as a tuple $\mathcal{M}_{\boldsymbol{\theta}} = (\boldsymbol{\Phi}_{\boldsymbol{\theta}_\Phi}, \boldsymbol{\Psi}_{\boldsymbol{\theta}_\Psi}, \mathbf{N}_{\boldsymbol{\theta}_N})$, with the subscript indicating that the model depends on a number of parameters $\boldsymbol{\theta} = (\boldsymbol{\theta}_\Phi, \boldsymbol{\theta}_\Psi, \boldsymbol{\theta}_N)$. Each of these components captures an aspect of landscape dynamics described above. Note that we will often exclude the subscripts, in which case the dependence of a component on the model parameters should be understood as implied.

##### The potential module

The first module consists of a neural network $\boldsymbol{\Phi}^{nn}_{\boldsymbol{\theta}_\Phi} : \mathbb{R}^d \to \mathbb{R}$, which maps vectors $\boldsymbol{x}$ in the phase space $\mathbb{R}^d$ to a scalar $\boldsymbol{\Phi}^{nn}_{\boldsymbol{\theta}_\Phi}(\boldsymbol{x})$, and which depends on the parameters $\boldsymbol{\theta}_\Phi$, the weights and biases of the linear layers defining the neural network. This module is meant to approximate the underlying static landscape $\tilde{\phi}^*$. Our assumption that dynamics are induced by a *smooth* potential suggests that we should enforce a smoothness on $\boldsymbol{\Phi}^{nn}_{\boldsymbol{\theta}_\Phi}$. To this end, we require a differentiable activation function for the hidden layers of $\boldsymbol{\Phi}^{nn}_{\boldsymbol{\theta}_\Phi}$, guaranteeing that it is differentiable with respect to $\boldsymbol{x}$. In addition, because we assume that the phase space $\Omega \subseteq \mathbb{R}^d$ has the topology of a $d$-dimensional disk, with flow from the boundary always directed inward as described in [19], we want to ensure that the flow induced by the potential is such that for some $r \in \mathbb{R}$, it holds that all trajectories starting at $\boldsymbol{x}$ with $\|\boldsymbol{x}\| > r$ eventually flow into the region $\Omega$. Therefore, instead of a "pure" neural network defining the potential, we include an additive *confinement term* $\Phi_0(\boldsymbol{x}) = C_{\text{conf}}\|\boldsymbol{x}\|^4$ where the constant $C_{\text{conf}} \geq 0$ is a *confinement factor* that is a hyperparameter (i.e. untrained parameter) of the model. By specifying this hyperparameter as a positive value, we are able to regularize the potential represented by the PLNN, according to

$$\boldsymbol{\Phi}_{\boldsymbol{\theta}_\Phi}(\boldsymbol{x}) = \Phi_0(\boldsymbol{x}) + \boldsymbol{\Phi}^{nn}_{\boldsymbol{\theta}_\Phi}(\boldsymbol{x}). \tag{S13}$$

Going forward, we will be just the slightest bit cavalier, and at times refer to the potential module as a neural network, ignoring the fact that there is an additional confinement term. This should be inconsequential since the neural network is a universal approximator, and the addition of the confinement term serves only to bias the form of the landscape at the outset of the training process, and does not include any learnable parameters.

Here we note that while we take $\boldsymbol{\Phi}$ to be a neural network, leveraging the properties of a universal function approximator, in principle a number of parameterized function could be utilized for $\boldsymbol{\Phi}$. For example, a Gaussian Mixture Model (GMM) consisting of a linear combination of multivariate gaussian functions could be used to model the landscape, with the components' means and covariances constituting the parameters $\boldsymbol{\theta}_\Phi$.

In both the synthetic case, in which we train a PLNN on *in silico* data, and the application to real, *in vitro* data, we use $C_{\text{conf}} = 0.1$. In testing, this adequately confines the trajectories without introducing an extreme gradient that would disrupt the training procedure, by necessitating a smaller timestep used by the SDE solver.

With respect to the neural network $\boldsymbol{\Phi}^{nn}$, we use in all cases a feed-forward network architecture with 4 hidden layers of widths 16, 32, 32, and 16. To the output of all hidden layers we applied the softplus activation function

$$\alpha(x) = \ln(1 + e^x),$$

guaranteeing a smooth potential. No activation function was applied to the final output of the model.

#### The tilt module

The second module models the signal processing function $\psi^*$, and we denote it $\boldsymbol{\Psi}_{\boldsymbol{\theta}_\Psi} : \mathbb{R}^{n_{\text{sigs}}} \to \mathbb{R}^d$. While $\boldsymbol{\Phi}$ is given a great deal of freedom as a neural network, $\boldsymbol{\Psi}$ is heavily restricted in its functional form. We assume that the signal is interpreted in a linear fashion, so that

$$\boldsymbol{\Psi}_{\boldsymbol{\theta}_\Psi}(\boldsymbol{s}) = \mathbf{A}_{\boldsymbol{\Psi}} \cdot \boldsymbol{s} \tag{S14}$$

where $\mathbf{A}_{\boldsymbol{\Psi}} \in \mathbb{R}^{d \times n_{\text{sigs}}}$ is a matrix whose entries constitute the model parameters $\boldsymbol{\theta}_\Psi$. We need not include a bias term in (S14), as the addition of a bias can be subsumed into the underlying landscape given by $\boldsymbol{\Phi}$.

#### The noise module

The third and final module of the PLNN incorporates the stochastic elements of landscape dynamics. We will discuss both the general construction of a noise module, and the much-simplified case that we implement and utilize in this body of work. In general, the noise module is a function $\mathbf{N}_{\boldsymbol{\theta}_N} : \mathbb{R}^d \to \mathbb{R}^{d \times d}$ that maps the state vector $\boldsymbol{x} \in \mathbb{R}^d$ to a noise kernel $\Sigma \in \mathbb{R}^{d \times d}$. This general construction allows for two features that we will shortly do away with, given the simplifying assumptions that we are making. It allows for state-dependent, or *heterogeneous* noise, as well as *non-isotropic* noise, so that there are greater stochastic effects in certain directions of phase space.

However, we will proceed under the assumption of homogeneous, isotropic noise, so that a single scalar $\sigma \in \mathbb{R}$ is sufficient to describe the stochasticity of the system, and this parameter $\sigma$ constitutes the single parameter of $\boldsymbol{\theta}_N$. We provide the general construction in its entirety for practical and theoretical reasons. Practically, it may be that in certain systems it is demonstrably not the case that noise is either isotropic or homogeneous, and in such systems one may wish to allow for a more general parameterization of the noise. This is easily achieved given the implementation of the PLNN. Theoretically, it is important to understand that the noise module outputs more than just a single scalar. It returns a higher-dimensional quantity, providing the diffusion term $\boldsymbol{g}$ in (S10).

#### Summary

In summary, a PLNN model $\mathcal{M}_{\boldsymbol{\theta}}$ with model parameters $\boldsymbol{\theta}$ can be written compactly as a tuple $\mathcal{M}_{\boldsymbol{\theta}} = (\boldsymbol{\Psi}_{\boldsymbol{\theta}}, \boldsymbol{\Phi}_{\boldsymbol{\theta}}, \mathbf{N}_{\boldsymbol{\theta}})$ with

$$\begin{aligned} \boldsymbol{\Phi}_{\boldsymbol{\theta}} &: \mathbb{R}^d \to \mathbb{R} \\ \boldsymbol{\Psi}_{\boldsymbol{\theta}} &: \mathbb{R}^{n_{\text{sigs}}} \to \mathbb{R}^d \\ \mathbf{N}_{\boldsymbol{\theta}} &: \mathbb{R}^d \to \mathbb{R}^{d \times d}. \end{aligned} \tag{S15}$$

The parameter vector $\boldsymbol{\theta}$ includes the weights and biases of the neural network defining the potential, the linear transformation defining the signal processing function, and the scalar noise parameter $\sigma$.

### S2B The PLNN as a generative model

The construction of the PLNN allows us to simulate landscape dynamics, via an SDE defined in terms of the PLNN modules. From a PLNN $\mathcal{M} = (\boldsymbol{\Phi}, \boldsymbol{\Psi}, \mathbf{N})$ we can consider an SDE of the same form as (S10):

$$\begin{aligned} d\boldsymbol{x}(t) &= \boldsymbol{f}(t, \boldsymbol{x}(t))dt + \boldsymbol{g}(t, \boldsymbol{x}(t))dW(t) \\ \boldsymbol{f}(t, \boldsymbol{x}(t)) &= -\nabla_{\boldsymbol{x}} \boldsymbol{\Phi}(\boldsymbol{x}) - \boldsymbol{\Psi}(\boldsymbol{s}(t)) \\ \boldsymbol{g}(t, \boldsymbol{x}(t)) &= \mathbf{N}(\boldsymbol{x}) \end{aligned} \tag{S16}$$

The PLNN $\mathcal{M}$ then serves as a generative model, which can be used to sample trajectories $\boldsymbol{x}(t)$ with $t \in [t_0, t_1]$, given an initial condition $\boldsymbol{x}_0 = \boldsymbol{x}(t_0)$ and a signal profile $\boldsymbol{s}(t)$ defined over the interval $[t_0, t_1]$.

In terms of inputs and outputs, we can think of a PLNN as taking in a tuple $(t_0, \boldsymbol{x}_0, t_1, \boldsymbol{s}(t))$ containing the initial condition, final time, and signal profile, and returning a sample trajectory $\boldsymbol{x}^t = \{\boldsymbol{x}(t) \,|\, t_0 \leq t \leq t_1\}$. We can simulate ensembles of cell trajectories, given a number $N_{\text{cells}}$ of initial conditions $\left\{\boldsymbol{x}^{(i)}(t_0)\right\}_{i=1}^{N_{\text{cells}}}$ all evolving under the influence of the same signal $\boldsymbol{s}(t)$. Letting $X_0$ be an $N_{\text{cells}} \times d$ matrix of initial conditions, where each row constitutes the state of an individual cell at time $t_0$, we can view the model as taking inputs $(t_0, X_0, t_1; \boldsymbol{s}(t))$ and returning a sample of ensemble trajectories $X^t = \left\{\boldsymbol{x}^{(i)}(t) \,|\, t_0 \leq t \leq t_1\right\}_{i=1}^{N_{\text{cells}}}$. In particular, considering only the final state of the ensemble at time $t_1$, we have a matrix output $X_1 \in \mathbb{R}^{N_{\text{cells}} \times d}$ of the same dimensions as $X_0$. This, then, is the essential point. Given a matrix of initial conditions, the model $\mathcal{M}$ returns a matrix of the same shape, representing the state of the system at a later time.

#### S2C Computational tools for vectorizing landscape dynamics

Sampling a trajectory of a single cell in the landscape requires solving (S16) over a specified interval, with a particular signal profile. Sampling the trajectory of an *ensemble* of cells requires solving the same SDE a number of times, once for each cell. Sampling the trajectory of *multiple ensembles*, each one potentially over a different time interval and with a distinct signal profile, requires yet another layer of vectorization. It is this tertiary level that we would like to achieve.

In order to efficiently handle the computational task at hand, we take advantage of the JAX computing ecosystem [23] that provides a number of automatic differentiation and vectorization capabilities. Specifically, we utilize the python package Diffrax [24] which implements a number of SDE solvers and allows for easy vectorization, allowing us to efficiently simulate the trajectories of multiple ensembles at once. Diffrax provides a number of SDE solvers, including the standard Euler-Maruyama method as well as higher-order methods.

Ultimately, we use backpropagation to infer model parameters. This can be achieved using one of a number of now standard packages, including TensorFlow [25] and PyTorch [26]. We choose to use JAX in particular because of the ease of automatically differentiating with respect to arbitrary vectors. This is a necessary requirement in our case, as for the drift term $\boldsymbol{f}$ we need to be able to compute the gradient of the potential module with respect to the state variable $\boldsymbol{x}$. This is made possible in JAX using simple, readily available built-in functions. In addition, the machine learning and optimization packages Equinox [27] and Optax [28] leverage the computational efficiency of JAX.

#### S2D Inferring a parameterized landscape system: The general PLNN training procedure

The outline for the training procedure is as follows: Given experimental data containing the initial and final state of an ensemble of cells, we use the model $\mathcal{M}$ to generate a *simulated* final state from the given initial state, and compare the observed and simulated final states in a distributional sense in order to update the model parameters. This procedure is depicted in Fig. 3 of the main text.

We assume that we have a training dataset $\mathcal{D}_{\text{train}}$ consisting of a number $N_{\text{train}}$ of datapoints:

$$\mathcal{D}_{\text{train}} = \{(t_0, X_0; t_1, X_1; \boldsymbol{s}(t))_i\}_{i=1}^{N_{\text{train}}} . \tag{S17}$$

Each datapoint tuple consists of an initial observation $X_0$, which is the observed state at time $t_0$, a later observation $X_1$ at $t_1$, and the signal profile $\boldsymbol{s}(t)$ which should be defined over the interval $[t_0, t_1]$. In general, the observed initial and final states need not contain the same cells, or even the same number of cells. Consider for example an experimental procedure in which a population of cells is periodically sampled, resulting in differently sized empirical samples of the population at a number of timepoints.

For a given training datapoint $(t_0, X_0; t_1, X_1; \boldsymbol{s}(t))$, a step of training involves first sampling from the SDE defined by (S16). The SDE is sampled by simulating the stochastic evolution from the initial condition at $t = t_0$ forward in time to $t = t_1$. However, for computational purposes, it is convenient to fix the number of cells internally represented by the model, and we denote this number by $N_{\text{cells}}$. Therefore, instead of using the observed initial condition $X_0$ directly, we first sample $N_{\text{cells}}$ cells with replacement from the rows of $X_0$, resulting in a matrix $\hat{X}_0 \in \mathbb{R}^{N_{\text{cells}} \times d}$. This matrix then serves as the input to the model $\mathcal{M}$, which returns a matrix of the same shape, the sampled final state of the cells.

We denote this *simulated* final state by $X_1^{(\text{sim})}$, which we can compare to the true, *observed* state $\boldsymbol{X}_1$, which we will now denote with a superscript, $X_1^{(\text{obs})}$, to make clear the distinction between the true, observed state in the dataset, and the simulated final state generated by the model. We can now compare the simulate and observed final states $X_1^{(\text{sim})}$ and $X_1^{(\text{obs})}$ via a prescribed loss function $\mathcal{L}(X^{(\text{sim})}, X^{(\text{obs})})$. We discuss our choice of loss function below, in S3. While the choice of loss function may of course vary depending on the particular use case, the essential condition is that we can compute gradients of the loss function with respect to the model parameters $\boldsymbol{\theta}$. We achieve this using the autodifferentiation tools provided by the JAX ecosystem [23, 27], which allow us to differentiate through the operations of the differential equation solver used to sample the SDE.

Training takes place over a number of epochs $N_{\text{epochs}}$, and we train in mini-batches of size $B$. Thus, a random selection of $B$ initial conditions is used to simultaneously simulate a number of ensemble trajectories forward in time, and each is then compared to the corresponding observed final state. The value of the loss is averaged over the batch, and an optimization step is taken to update the model parameters $\boldsymbol{\theta}$. We use the optimization package Optax [28] to perform the optimization step, using the RMSProp algorithm [29].

### S3 Choice of loss function

As part of the training of a PLNN, we require a loss function that assesses the difference between two empirical samples $X$ and $Y$, where $X$ is the observed ensemble state at a given time, and $Y$ is the simulated ensemble state, the output of the model. We investigate two approaches to this two-sample problem. The first utilizes the KL divergence, and the second the maximum mean discrepancy (MMD), both measures being applicable to discerning whether two samples have been drawn from the same distribution [30, 31].

#### S3A A loss function estimating the KL divergence

The Kullback-Leibler (KL) divergence [32] provides a measure of the distance between distributions. It is defined for densities $P$ and $Q$ according to

$$D_{KL}(P\|Q) = \int_{\mathbb{R}^d} p(x) \log \frac{p(x)}{q(x)} dx \geq 0 \tag{S18}$$

and is zero only if $P = Q$. In our context, we would like to estimate the KL divergence $D_{KL}(P\|Q)$ where $P$ corresponds to the observed data $X \sim p(x)$, and $Q$ corresponds to the simulated data $Y \sim q(x)$. To this end, we take advantage of an estimate of the KL divergence introduced in Perez-Cruz [33]. This method uses $k$-nearest-neighbor density estimation and does not rely on a prior estimate of the density functions $p$ and $q$. Moreover, the algorithm is applicable to vectorial data of arbitrary dimension. The primary result from Perez-Cruz [33] that we employ is that given $n$ i.i.d. samples from $p(x)$ and $m$ i.i.d. samples from $q(x)$, an estimate $\hat{D}_{KL}(P\|Q)$ of the KL divergence $D_{KL}(P\|Q)$ is given by

$$\hat{D}_{KL}(P\|Q) = \frac{1}{n}\sum_{i=1}^{n} \log \frac{\hat{p}_k(\boldsymbol{x}_i)}{\hat{q}_k(\boldsymbol{x}_i)} = \frac{d}{n}\sum_{i=1}^{n} \log \frac{r_k(\boldsymbol{x}_i)}{s_k(\boldsymbol{x}_i)} + \log \frac{m}{n-1} \tag{S19}$$

with

$$\begin{aligned} \hat{p}_k(\boldsymbol{x}_i) &= \frac{k}{(n-1)} \frac{\Gamma(d/2+1)}{\pi^{d/2} r_k(\boldsymbol{x}_i)^d} \\ \hat{q}_k(\boldsymbol{x}_i) &= \frac{k}{m} \frac{\Gamma(d/2+1)}{\pi^{d/2} s_k(\boldsymbol{x}_i)^d} \end{aligned} \tag{S20}$$

and $r_k(\boldsymbol{x}_i)$ and $s_k(\boldsymbol{x}i)$ are the Eulidean distances to the $k$th nearest-neighbors of $\boldsymbol{x}_i$ in $X\backslash\boldsymbol{x}_i$ and $Y$, respectively.

We define the KL divergence-based loss function $\mathcal{L}_{KL}(X^{(\text{sim})}, X^{(\text{obs})})$ as

$$\mathcal{L}_{KL}(X^{(\text{sim})}, X^{(\text{obs})}) = \hat{D}_{KL}(X^{(\text{obs})}\|X^{(\text{sim})}) \tag{S21}$$

taking $k = 1$ for an estimate based on the first-nearest neighbor.

Thus, the loss function compares the similarity of the pairwise distances between points *within* the sample $X$, with the distances between the points in $X$ and the points in $Y$, computing the $k$-nearest neighbors for the pairwise distances $d(\boldsymbol{x}_i, \boldsymbol{x}_j)$ within $X$ and the cross-distances $d(\boldsymbol{x}_i, \boldsymbol{y}_j)$ between $X$ and $Y$. Because of this, the resulting loss is not affected by infinitesimal changes to points that are not among the nearest neighbors of either set, and thus the gradient of the loss with respect to these points is zero. We suspect that this reliance on only a fraction of the points in either sample results in a slower training process, and thus turn our consideration to a loss function that utilizes *all* points.

#### S3B A loss function estimating the maximum mean discrepancy

The second loss function that we consider, $\mathcal{L}_{MMD}$, utilizes a kernel method for the two-sample test proposed by Gretton et al. [31]. For the sake of completeness, the following summarizes the discussion of the MMD presented in the original work. This statistical test assesses the similarity between distributions $p$ and $q$ using a family $\mathcal{F}$ of test functions, where $\mathcal{F}$ is the unit ball in a reproducing kernel Hilbert space (RKHS) $\mathcal{H}$. An RKHS $\mathcal{H}$ is a Hilbert space of real-valued functions defined on a non-empty set $\mathcal{X}$ with inner product $\langle\cdot,\cdot\rangle_{\mathcal{H}}$, for which there exists a function $k : \mathcal{X} \times \mathcal{X} \to \mathbb{R}$ satisfying

- $\forall x \in \mathcal{X}$, $k(\cdot, x) \in \mathcal{H}$, and
- $\forall x \in \mathcal{X}, \forall f \in \mathcal{H}$, $\langle f(\cdot), k(\cdot, x)\rangle_{\mathcal{H}} = f(x)$.

The second condition expresses a reproducing property, and the function $k$ is called a *reproducing kernel*.

For a class $\mathcal{F}$ of functions $f : \mathcal{X} \to \mathbb{R}$, the *maximum mean discrepancy* (MMD) between $p$ and $q$ is defined as

$$\text{MMD}[\mathcal{F}, p, q] = \sup_{f\in\mathcal{F}} \left(\mathbb{E}_x[f(x)] - \mathbb{E}_y[f(y)]\right). \tag{S22}$$

Given a distribution $p$, the *mean embedding* of $p$ is an element $\mu_p \in \mathcal{H}$ satisfying $\mathbb{E}_x[f] = \langle f, \mu_p\rangle_{\mathcal{H}}$ for all $f \in \mathcal{H}$. The MMD can be expressed as a distance in $\mathcal{H}$ between mean embeddings (assuming they exist):

$$\text{MMD}^2[\mathcal{F}, p, q] = \|\mu_p - \mu_q\|^2_{\mathcal{H}}. \tag{S23}$$

Then, the question becomes whether the mean embedding $\mu_p$ is injective. If so, it follows that the MMD is a metric on probability distributions over $\mathcal{X}$. A sufficient condition for injectiviy of $\mu_p$ is that $\mathcal{H}$ be a *universal* RKHS on compact metric space $\mathcal{X}$. The universality condition stipulates $k(\cdot,\cdot)$ be continuous and $\mathcal{H}$ dense in $C(\mathcal{X})$ with respect to the $L_\infty$ norm [31]. A more general result due to Fukumizu et al. [34] introduces *characteristic kernels* as those for which the mean map is injective, and establishes the characteristic property of Gaussian and Laplace kernels on the entirety of $\mathbb{R}^d$. For our purposes, it suffices that Gaussian RKHSs are characteristic, and these are the kernel functions that we make use of.

Thus, by minimizing the MMD, we can minimize the discrepancy between the distributions $p$ and $q$. In practice we rely on the finite samples $X$ and $Y$ in place of the true distributions $p$ and $q$, and use the following unbiased estimate of the MMD, the culmination of the work presented by Gretton et al. [31]. For $x, x' \sim_{iid} p$ and $y, y' \sim_{iid} q$, the squared population MMD can be found in terms of the kernel, according to

$$\text{MMD}^2[\mathcal{F}, p, q] = \mathbb{E}_{x,x'}\left[k(x,x')\right] - 2\mathbb{E}_{x,y}\left[k(x,y)\right] + \mathbb{E}_{y,y'}\left[k(y,y')\right]. \tag{S24}$$

An unbiased empirical estimate is then given by

$$\begin{aligned}\text{MMD}_u^2[\mathcal{F}, X, Y] = &\frac{1}{n(n-1)}\sum_{i\neq j}^{n} k(x_i, x_j)\\ &+ \frac{1}{m(m-1)}\sum_{i\neq j}^{m} k(y_i, y_j)\\ &- \frac{2}{mn}\sum_{i=1}^{n}\sum_{j=1}^{m} k(x_i, y_j).\end{aligned} \tag{S25}$$

We use this unbiased estimate of the MMD to define an MMD-based loss function:

$$\mathcal{L}_{\text{MMD}}(X^{(\text{sim})}, X^{(\text{obs})}) = \text{MMD}_u^2[\mathcal{F}, X^{(\text{sim})}, X^{(\text{obs})}] \tag{S26}$$

where for our purposes $\mathcal{F}$ is a Gaussian RKHS. We take $k$ to be given by a combination of Gaussians with chosen bandwidth(s), specified as a hyperparameter of the training process. In practice this hyperparameter can be chosen in accordance with the median distance between observed datapoints [35], although additional methods can be used to select a suitable bandwidth [36].

### S4 *In silico* experiments

In this section, we describe a number of *in silico* experiments in which we infer a PLNN from synthetic data, generated within a known landscape system. In S4A we train models on data generated in the binary choice and binary flip systems, providing full details with respect to the training data, model architecture, and hyperparameters. In S4B, we detail the synthetic experiments used to assess the robustness of model training to the temporal sampling resolution.

#### S4A Inferring synthetic binary decision landscapes

##### S4A.1 Generating synthetic training data

In order to asses the ability of a PLNN to accurately infer parameterized landscape dynamics, we generate data via *in silico* simulations where the ground-truth system is known. That is, we specify the underlying static landscape, tilting effect, and noise *a priori*, thereby defining a ground-truth model $\mathcal{M}^*$. We then simulate a number $N_{\text{exps}}$ of experiments in which a fixed number $N_{\text{cells}}$ of cells evolve in the landscape over an interval $t \in [0, T]$, under a signal profile $\boldsymbol{s}^{(j)}(t)$, where $j$ is an index ranging over the number of experiments.

For these *in silico* experiments, we take for the ground-truth either the binary choice or binary flip landscape, as described above, and given by either (S6) or (S7) We specify an initial state for all cells in the ensemble, and let the system evolve for a burn-in period under the value of a burn-in signal $\boldsymbol{s}_{\text{burn}}$. The state of the ensemble at the end of the burn-in period is used as the initial condition at time $t = 0$. This ensures distinct, non-delta distributions of cells at the start of each experiment. Over the course of each experiment $j$, we sample the state of the full ensemble $X(t) \in \mathbb{R}^{N_{\text{cells}} \times d}$ at evenly spaced intervals of $\Delta T$, beginning with the initial state $X(0)$. In this way, we generate a training dataset in the form of (S17) where each datapoint $d^{(i)}$ comes from one of the experiments, and consists of a starting state $X_0^{(i)}$ observed at time $t_0^{(i)}$ and an ending state $X_1^{(i)}$ observed at time $t_1^{(i)} = t_0^{(i)} + \Delta T$, as well as the signal profile $\boldsymbol{s}^{j(i)}(t)$, where $j(i)$ gives the association of each datapoint to a specific experiment.

Using either the binary choice or the binary flip landscape, we have an underlying potential $\tilde{\phi}^*(\boldsymbol{x})$ with $\boldsymbol{x} = (x, y)^T$, which is tilted in response to a 2-dimensional signal profile

$$\boldsymbol{s}(t) = (s_1(t),\, s_2(t))^T\,.$$

We use sigmoidal functions for the signal profiles, such that

$$s_k(t) = a_k + \frac{1}{2}\left(b_k - a_k\right)\tanh\left(r_k\left(t - t_{ck}\right)\right),\quad k \in \{1, 2\} \tag{S27}$$

where $a_k$ and $b_k$ are the extremes of the $k$th component of the signal vector, $t_{ck}$ is the critical point at which the signal component is halfway between its extremes, and $r_k$ specifies its rate of transition. Using this parameterization of the signal profile, we can simulate a diversity of experiments by sampling the signal parameters from a specified distribution.

We assume that the tilt effect is a linear function of the input signal, as in (S9), so that we can write the ground-truth, *in silico* parameterized landscape as

$$\begin{aligned}\phi^*(\boldsymbol{x}; \boldsymbol{s}(t)) &= \tilde{\phi}^*(\boldsymbol{x}) + \boldsymbol{x}^T\psi^*(\boldsymbol{s}(t))\\ \psi^*(\boldsymbol{s}) &= \mathbf{A}^*\boldsymbol{s}\end{aligned} \tag{S28}$$

with $\mathbf{A}^* \in \mathbb{R}^{2\times 2}$ prescribed *a priori*, defining the tilt effect. The governing dynamics for each cell in the system is then provided by a stochastic differential equation

$$d\boldsymbol{x}_c(t) = -\nabla_{\boldsymbol{x}}\phi^*(\boldsymbol{x}_c(t), \boldsymbol{s}^{(i)}(t))dt + \sigma^* dW_c \tag{S29}$$

where $c = 1, \ldots, N_{\text{cells}}$ indexes across the cells in the system. $\sigma^*$ is a global parameter defining the magnitude of noise in the system, and $W_c$ is a $d$-dimensional Wiener process describing the stochastic fluctuations of cell $c$. It is this SDE that we then simulate for each cell in the system in order to obtain a synthetic dataset.

#### S4A.2 Inferring the binary choice landscape

We first apply the PLNN modeling framework to infer the binary choice landscape given by (S6). We define

$$\begin{aligned}\phi_1^*(\boldsymbol{x}; \boldsymbol{s}(t)) &= \tilde{\phi}_1^*(\boldsymbol{x}) + \boldsymbol{x}^T\psi_1^*(\boldsymbol{s}(t))\\ \tilde{\phi}_1^*(\boldsymbol{x}) &= \tilde{\phi}_{bc} = x^4 + y^4 + y^3 - 4x^2y + y^2\\ \psi_1^*(\boldsymbol{s}) &= \mathbf{A}_1^*\boldsymbol{s}\\ \mathbf{A}_1^* &= \boldsymbol{I}_2\\ \sigma^* &= 0.1.\end{aligned} \tag{S30}$$

This is the binary choice landscape, parameterized by signals according to $\boldsymbol{\tau} = \psi_1^*(\boldsymbol{s}) = \boldsymbol{s}$. For simplicity, we have assumed the identity mapping for the signal processing function $\psi_1^*$.

To generate *in silico* training data, we perform a series of $N_{\text{exps}} = 100$ experiments, each subject to a distinct signal profile, and consisting of an ensemble of $N_{\text{cells}} = 500$ cells. The complete list of parameters used for these simulations are shown in Table S1. The resulting data is pooled across experiments to yield a training dataset

$$\mathcal{D}_{1,\text{train}} = \left\{\left(t_0^{(i)},\, X_0^{(i)};\, t_1^{(i)},\, X_1^{(i)};\, \boldsymbol{s}^{j(i)}(t)\right)\right\}_{i=1}^{1000}\,. \tag{S31}$$

Movie 1 exemplifies this data generating process for a single experiment.

In addition to generating the training dataset $\mathcal{D}_{1,\text{train}}$, we also generate a validation dataset $\mathcal{D}_{1,\text{valid}}$ and a test dataset $\mathcal{D}_{1,\text{test}}$, each consisting of $20$ experimental runs, in an identical fashion as the training dataset. The purpose of the validation dataset is to serve as an out-of-sample test of the model during the training process. The purpose of the test dataset is to evaluate the model's performance after training is complete, and to compare models trained using different hyperparameters.

The neural network architecture and initial parameter values are specified in Table S2. We train the model in mini-batches, using the RMSProp algorithm [29], implemented in Optax [28], with an exponentially decaying learning rate. The complete list of parameters used for the training process is detailed in Table S3. We implement early stopping, and halt training if the validation loss does not improve for a specified number of epochs. In addition, in order to balance computational efficiency with accuracy, at the start of the training process we use a relative large timestep $\delta t = 0.1$ within the SDE solver, and periodically reduce this timestep by a factor of $0.5$ over the course of training. This allows us to form an initial approximation to the underlying landscape in a relatively short amount of time, and then to improve this approximation using a more accurate, but more computationally demanding, timestep. We use Heun's method as our choice of SDE solver, a second order explicit Runge–Kutta method, which provides sufficient accuracy while maintaining computational efficiency [37]. Fig. S3 details the evolution of the model over the course of training, and Fig. S4 shows the final inferred model.

#### S4A.3 Inferring the binary flip landscape

As a second example, we apply the PLNN modeling framework to the binary flip landscape given by (S7). We define

$$\begin{aligned}
\phi_2^*(\boldsymbol{x};\boldsymbol{s}(t)) &= \tilde{\phi}_2^*(\boldsymbol{x}) + \boldsymbol{x}^T\psi_2^*(\boldsymbol{s}(t)) \\
\tilde{\phi}_2^*(\boldsymbol{x}) &= \tilde{\phi}_{bf} = x^4 + y^4 + x^3 - 2xy^2 - x^2 \\
\psi_2^*(\boldsymbol{s}) &= \mathbf{A}_2^*\boldsymbol{s} \\
\mathbf{A}_2^* &= \boldsymbol{I}_2 \\
\sigma^* &= 0.3.
\end{aligned} \tag{S32}$$

Again, we assume an identity map between signals and tilts. The parameters used to generate the dataset are shown in Table S4. Movie 2 shows an example experiment. We use a larger noise scale in this case, as binary flip landscape has been shown to be useful in modeling stochastically-driven cell transitions [1].

The training procedure for this case is analogous to that of the binary choice landscape. Relevant hyperparameters are shown in Tables S5 and S6, while Figs. S5 and S6 detail the training process and resulting model.

| Parameter | Value | Description |
|---|---|---|
| $N_{\text{exps}}$ | 100 | Number of simulated experiments. |
| $N_{\text{cells}}$ | 500 | Number of cells in the ensemble. |
| $\boldsymbol{x}_0$ | $(0, -0.5)$ | Initial state of all cells before burn-in phase. |
| $T$ | 100.0 | Simulation ending time. |
| $\Delta t$ | 0.001 | Euler-Maruyama timestep. |
| $\Delta T$ | 10.0 | Sampling interval. |
| $T_{\text{burn}}$ | $0.1T$ | Duration of burn-in phase. |
| $\boldsymbol{s}_{\text{burn}}$ | $\boldsymbol{s}(0)$ | Burn-in signal. |
| $\sigma^*$ | 0.1 | Noise scale. |
| $\pi(a_1)$ | $\mathcal{U}[-0.5, 0.5]$ | Dist. of signal profile parameter $a_1 = \lim_{t\to-\infty} s_1(t)$. |
| $\pi(a_2)$ | $\mathcal{U}[0.5, 1.5]$ | Dist. of signal profile parameter $a_2 = \lim_{t\to-\infty} s_2(t)$. |
| $\pi(b_1)$ | $\mathcal{U}[-1, 1]$ | Dist. of signal profile parameter $b_1 = \lim_{t\to\infty} s_1(t)$. |
| $\pi(b_2)$ | $\mathcal{U}[-0.5, 0.5]$ | Dist. of signal profile parameter $b_2 = \lim_{t\to\infty} s_2(t)$. |
| $\pi(t_{c1})$ | $\mathcal{U}[0.1T, 0.9T]$ | Dist. of signal profile parameter $t_{c1}$. |
| $\pi(t_{c2})$ | $\mathcal{U}[0.1T, 0.9T]$ | Dist. of signal profile parameter $t_{c2}$. |
| $\pi(r_1)$ | $\ln r_1 \sim \mathcal{U}[-3, 2]$ | Dist. of signal profile parameter $r_1$. |
| $\pi(r_2)$ | $\ln r_2 \sim \mathcal{U}[-3, 2]$ | Dist. of signal profile parameter $r_2$. |
| $N_{\text{samps}}$ | $1 + (T/\Delta T) = 11$ | Number of sampling timepoints per experiment. |
| $\lvert\mathcal{D}\rvert$ | $N_{\text{exps}} \cdot (N_{\text{samps}} - 1) = 1000$ | Total number of datapoints generated. |

Table S1: Parameters used to generate the *in silico* dataset $\mathcal{D}_{1,\text{train}}$ using the ground-truth model $\mathcal{M}_1^*$ given by the binary choice system (S30).

| Parameter | Value | Description |
|---|---|---|
| $\dim(\boldsymbol{\Phi})$ | $[2, 16, 32, 32, 16, 1]$ | $\boldsymbol{\Phi}^{nn}$ layer sizes from input to output. |
| $\alpha^h_{\boldsymbol{\Phi}}$ | `softplus` | Activation function for the hidden layers of $\boldsymbol{\Phi}^{nn}$. |
| $\alpha^f_{\boldsymbol{\Phi}}$ | `identity` | Activation function for the final layer of $\boldsymbol{\Phi}^{nn}$. |
| $C_{\text{conf}}$ | 0.1 | Confinement factor. |
| $\sigma_0$ | 0.05 | Initial value of noise parameter $\sigma$. |
| $\boldsymbol{\Phi}^0_W$ | `uniform xavier` | Initialization scheme for the weights of $\boldsymbol{\Phi}^{nn}$. |
| $\boldsymbol{\Phi}^0_b$ | 0.0 | Initialization scheme for the biases of $\boldsymbol{\Phi}^{nn}$. |
| $\mathbf{A}^0$ | `uniform xavier` | Initialization scheme for the weights of $\boldsymbol{\Psi}$. |

Table S2: Hyperparameters used to define the architecture and initialization of PLNN $\mathcal{M}_1$, intended to infer the binary choice landscape using the training data $\mathcal{D}_{1,\text{train}}$.

| Parameter | Value | Description |
|---|---|---|
| $N_{\text{epochs}}$ | 2000 | Number of training epochs. |
| `patience` | 100 | Number of epochs before early halting. |
| `min_epochs` | 500 | Minimum number of required epochs. |
| $B$ | 250 | Batch size. |
| $N_{\text{cells}}$ | 200 | Number of cells simulated in each ensemble. |
| `solver` | `heun` | SDE Solver method. |
| $\Delta t$ | $0.1, [200 : 0.5, 500 : 0.5, 1000 : 0.5]$ | Timestep schedule (initial, [epoch:reduction]). |
| $\mathcal{L}$ | $MMD$ | Loss function. |
| `kernel` | Gaussian | MMD kernel function. |
| $\gamma$ | $[0.2, 0.5, 0.9, 1.3]$ | kernel bandwidth parameter(s). |
| `optimizer` | `RMSProp` | Optimizer algorithm. |
| `momentum` | 0.5 | Optimizer hyperparameter. |
| `weight decay` | 0.9 | Optimizer hyperparameter. |
| `clipping` | 1.0 | Gradient clipping hyperparameter. |
| `lr` | `exponential`$[10^{-2}, 10^{-5}, 50]$ | Learning rate schedule [initial, final, warmup]. |

Table S3: Hyperparameters used for the training of PLNN $\mathcal{M}_1$, trained on the *in silico* dataset $\mathcal{D}_{1,\text{train}}$ corresponding to the binary choice landscape.

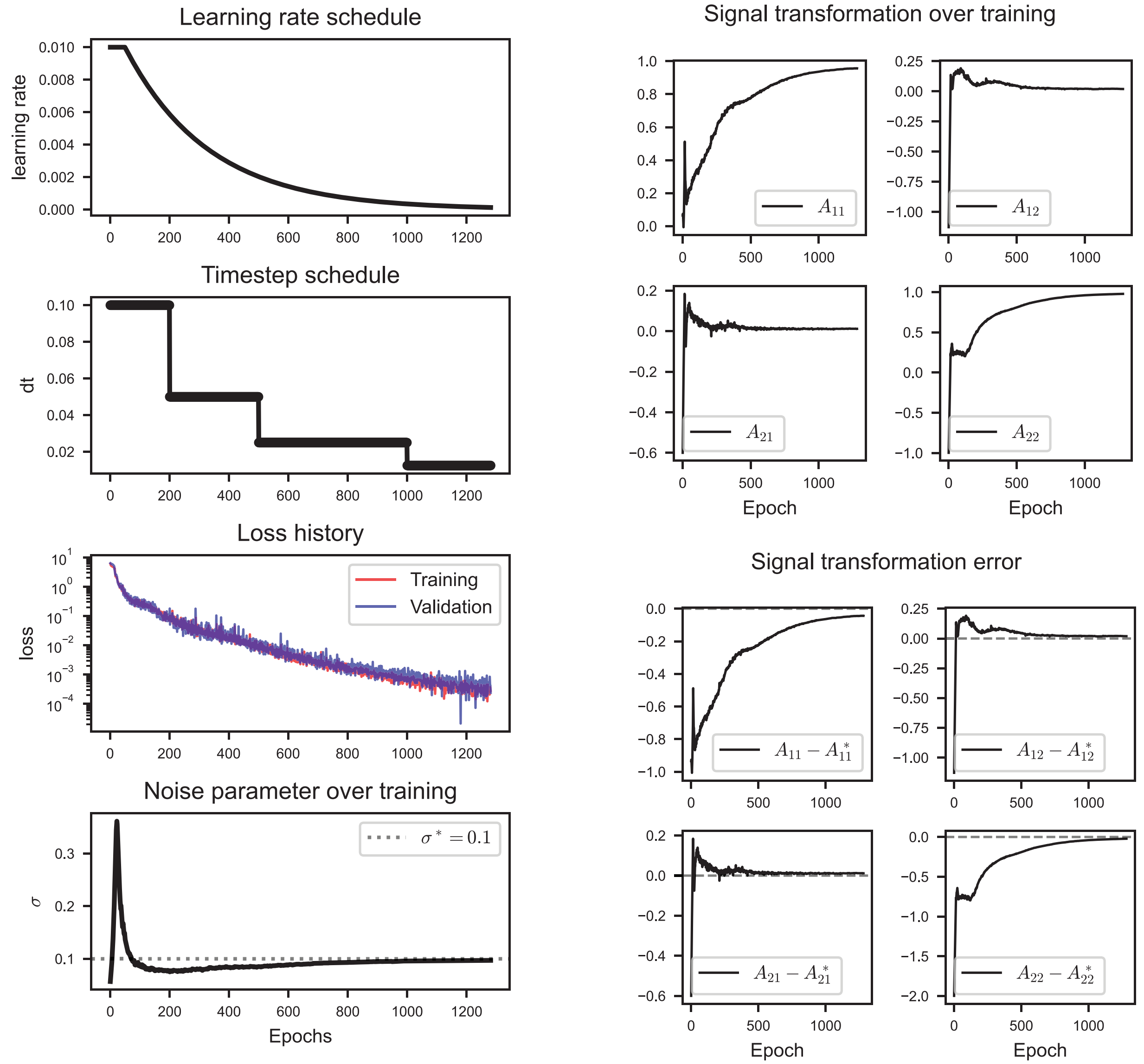

Figure S3: Evolution of the model $\mathcal{M}_1$ over the course of training. Subplots include the training and validation loss history; the learning rate and timestep over the course of training; and the evolution of the model parameters $\mathbf{A}$ (weights defining $\mathbf{\Psi}$) and $\sigma$ (noise scale).

| Parameter | Value | Description |
|---|---|---|
| $N_{\text{exps}}$ | 100 | Number of simulated experiments. |
| $N_{\text{cells}}$ | 500 | Number of cells in the ensemble. |
| $\boldsymbol{x}_0$ | $(-1, 0)$ | Initial state of all cells before burn-in phase. |
| $T$ | 100.0 | Simulation ending time. |
| $\Delta t$ | 0.001 | Euler-Maruyama timestep. |
| $\Delta T$ | 10.0 | Sampling interval. |
| $T_{\text{burn}}$ | $0.05T$ | Duration of burn-in phase. |
| $\boldsymbol{s}_{\text{burn}}$ | $(-0.25, 0)$ | Burn-in signal. |
| $\sigma^*$ | 0.3 | Noise scale. |
| $\pi(a_1)$ | $\mathcal{U}[-1.5, -1.0]$ | Dist. of signal profile parameter $a_1 = \lim_{t\to-\infty} s_1(t)$. |
| $\pi(a_2)$ | $\mathcal{U}[-0.75, 0.75]$ | Dist. of signal profile parameter $a_2 = \lim_{t\to-\infty} s_2(t)$. |
| $\pi(b_1)$ | $\mathcal{U}[-1.5, -1]$ | Dist. of signal profile parameter $b_1 = \lim_{t\to\infty} s_1(t)$. |
| $\pi(b_2)$ | $\mathcal{U}[-0.75, 0.75]$ | Dist. of signal profile parameter $b_2 = \lim_{t\to\infty} s_2(t)$. |
| $\pi(t_{c1})$ | $\mathcal{U}[0.1T, 0.9T]$ | Dist. of signal profile parameter $t_{c1}$. |
| $\pi(t_{c2})$ | $\mathcal{U}[0.1T, 0.9T]$ | Dist. of signal profile parameter $t_{c2}$. |
| $\pi(r_1)$ | $\ln r_1 \sim \mathcal{U}[-3, 2]$ | Dist. of signal profile parameter $r_1$. |
| $\pi(r_2)$ | $\ln r_2 \sim \mathcal{U}[-3, 2]$ | Dist. of signal profile parameter $r_2$. |
| $N_{\text{samps}}$ | $1 + (T/\Delta T) = 11$ | Number of sampling timepoints per experiment. |
| $\lvert\mathcal{D}\rvert$ | $N_{\text{exps}} \cdot (N_{\text{samps}} - 1) = 1000$ | Total number of datapoints generated. |

Table S4: Parameters used to generate the *in silico* dataset $\mathcal{D}_{2,\text{train}}$ using the ground-truth model $\mathcal{M}_2^*$ given by the binary flip system (S32).

| Parameter | Value | Description |
|---|---|---|
| $\dim(\mathbf{\Phi})$ | $[2, 16, 32, 32, 16, 1]$ | $\mathbf{\Phi}^{nn}$ layer sizes from input to output. |
| $\alpha_{\mathbf{\Phi}}^h$ | `softplus` | Activation function for the hidden layers of $\mathbf{\Phi}^{nn}$. |
| $\alpha_{\mathbf{\Phi}}^f$ | `identity` | Activation function for the final layer of $\mathbf{\Phi}^{nn}$. |
| $C_{\text{conf}}$ | 0.1 | Confinement factor. |
| $\sigma_0$ | 0.05 | Initial value of noise parameter $\sigma$. |
| $\mathbf{\Phi}_W^0$ | `uniform xavier` | Initialization scheme for the weights of $\mathbf{\Phi}^{nn}$. |
| $\mathbf{\Phi}_b^0$ | 0.0 | Initialization scheme for the biases of $\mathbf{\Phi}^{nn}$. |
| $\mathbf{A}^0$ | `uniform xavier` | Initialization scheme for the weights of $\mathbf{\Psi}$. |

Table S5: Hyperparameters used to define the architecture and initialization of PLNN $\mathcal{M}_2$, intended to infer the binary flip landscape using the training data $\mathcal{D}_2$.

| Parameter | Value | Description |
|---|---|---|
| $N_{\text{epochs}}$ | 2000 | Number of training epochs. |
| `patience` | 100 | Number of epochs before early halting. |
| `min_epochs` | 500 | Minimum number of required epochs. |
| $B$ | 250 | Batch size. |
| $N_{\text{cells}}$ | 200 | Number of cells simulated in each ensemble. |
| `solver` | `heun` | SDE Solver method. |
| $\Delta t$ | $0.1, [200 : 0.5, 500 : 0.5, 1000 : 0.5]$ | Timestep schedule (initial, [epoch:reduction]). |
| $\mathcal{L}$ | $MMD$ | Loss function. |
| `kernel` | Gaussian | MMD kernel function. |
| $\gamma$ | $[0.2, 0.5, 0.9, 1.3]$ | kernel bandwidth parameter(s). |
| `optimizer` | `RMSProp` | Optimizer algorithm. |
| `momentum` | 0.5 | Optimizer hyperparameter. |
| `weight decay` | 0.9 | Optimizer hyperparameter. |
| `clipping` | 1.0 | Gradient clipping hyperparameter. |
| `lr` | `exponential`$[10^{-2}, 10^{-5}, 50]$ | Learning rate schedule [initial, final, warmup]. |

Table S6: Hyperparameters used for the training of PLNN $\mathcal{M}_2$, trained on the *in silico* dataset $\mathcal{D}_{2,\text{train}}$ corresponding to the binary flip landscape.

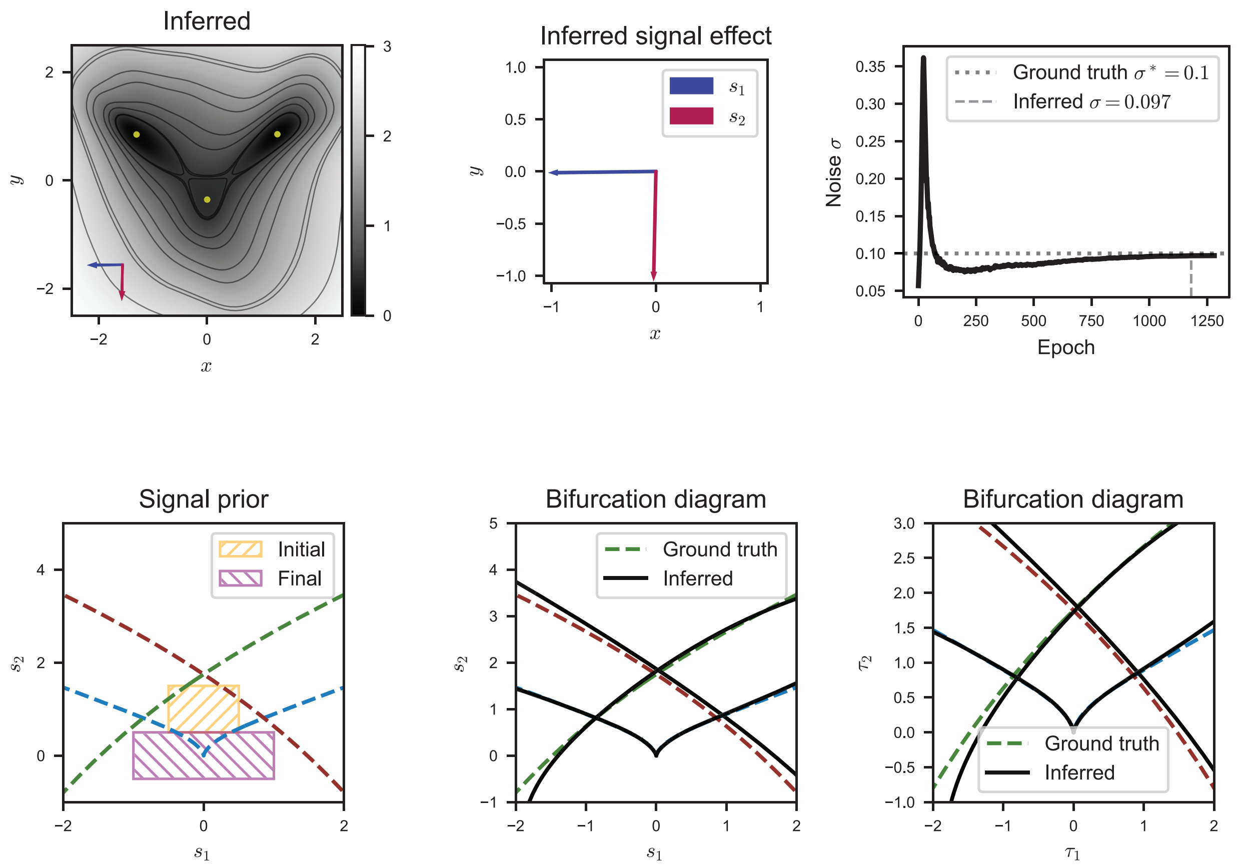

Figure S4: The inferred model $\mathcal{M}_1$.

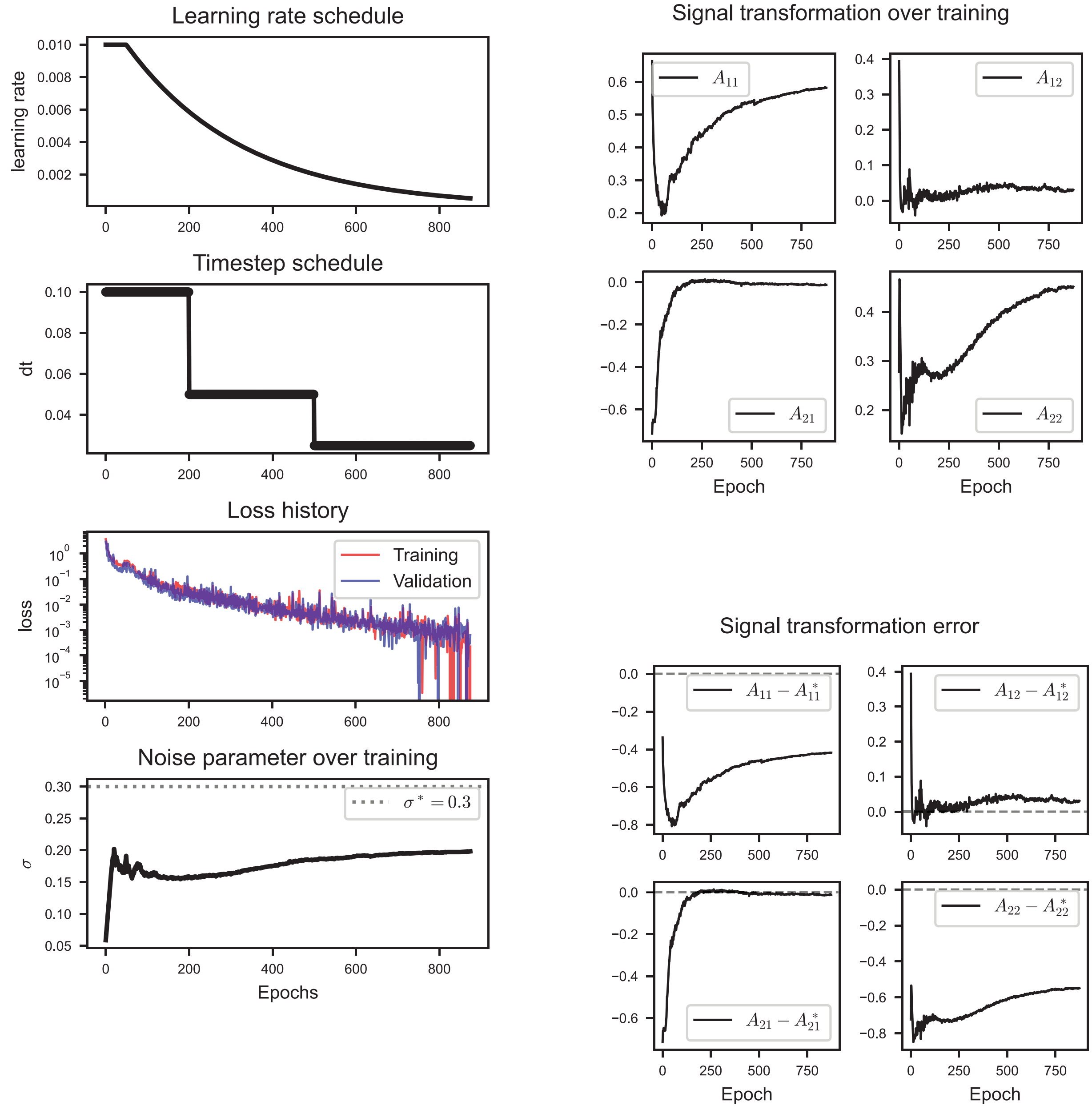

Figure S5: Evolution of the model $\mathcal{M}_2$ over the course of training. Subplots include the training and validation loss history; the learning rate and timestep over the course of training; and the evolution of the model parameters $\mathbf{A}$ (weights defining $\mathbf{\Psi}$) and $\sigma$ (noise scale).

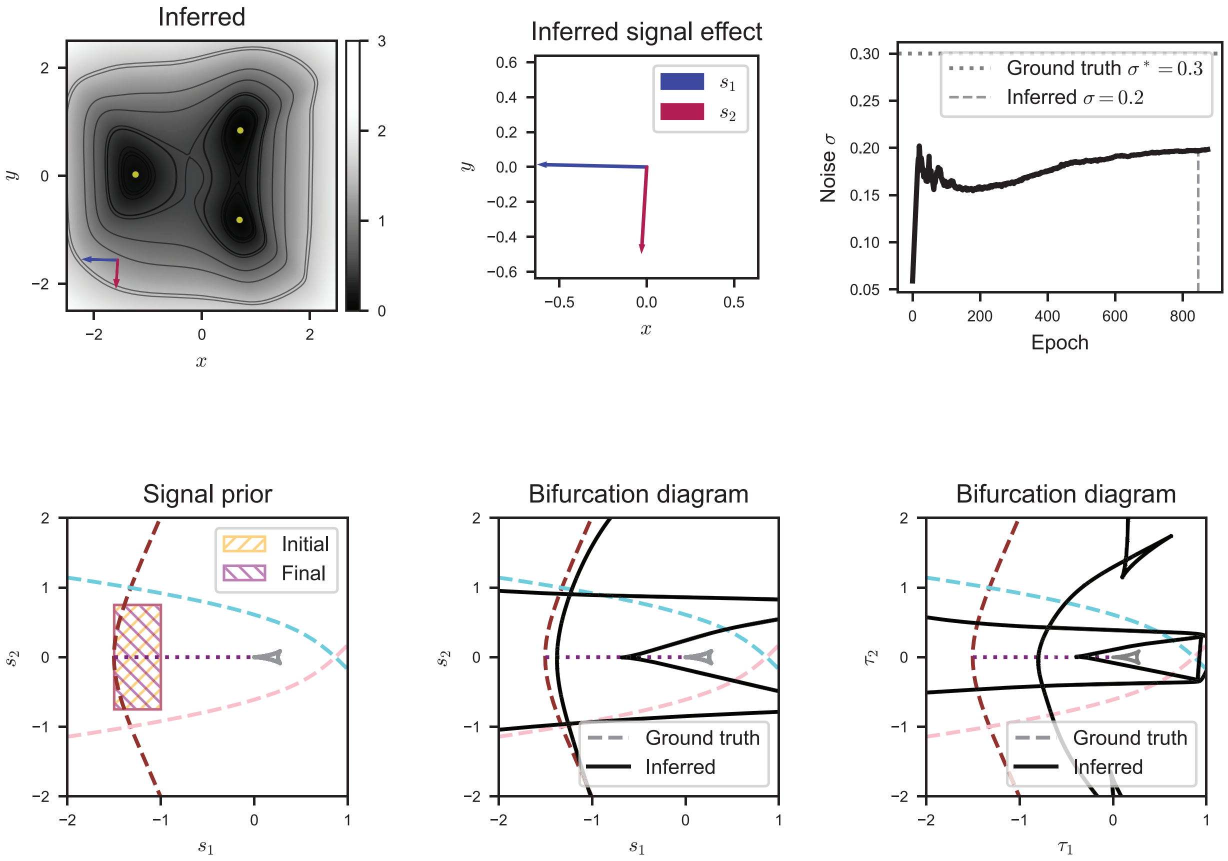

Figure S6: The inferred model $\mathcal{M}_2$.

#### S4B Inference robustness to temporal sampling resolution

The rate at which sampling takes place in the data collection process impacts how accurately we are able to infer an underlying decision landscape. In order to quantitatively assess this sensitivity of the model inference procedure, we generated a number of synthetic datasets in which we varied the rate of sampling. Table S7 shows the parameters used to generate these datasets. The sampling interval $\Delta T$, defined as the interval of time between consecutive snapshots, is the inverse of the sampling rate, and is the principal parameter that we vary. Changing $\Delta T$, however, changes the number of consecutive snapshot pairs generated for each signaling condition, and we therefore also vary the number of simulated experiments for each dataset, so that the total number of training datapoints is consistent.

In order to ensure that a transition takes place, and that this transition happens over a consistent interval of time, we specify particular distributions for the parameters defining the sigmoidal signal profiles. We require that the transition rate is fast, and that the signal change takes place within the interval $0.1T$ to $0.15T$, where $T$ is the duration of the experiment.

| Parameter | Value | Description |
|---|---|---|
| $N_{\text{exps}}$ | $[50, 100, 200]$ | Number of simulated experiments. |
| $N_{\text{cells}}$ | 200 | Number of cells in the ensemble. |
| $\boldsymbol{x}_0$ | $(0, -0.5)$ | Initial state of all cells before burn-in phase. |
| $T$ | 20.0 | Simulation ending time. |
| $\Delta t$ | 0.001 | Euler-Maruyama timestep. |
| $\Delta T$ | $[5, 10, 20]$ | Sampling interval. |
| $T_{\text{burn}}$ | $0.1T$ | Duration of burn-in phase. |
| $\sigma^*$ | 0.1 | Noise scale. |
| $\pi(a_1)$ | $\mathcal{U}[-0.5, 0.5]$ | Dist. of signal profile parameter $a_1 = \lim_{t\to-\infty} s_1(t)$. |
| $\pi(a_2)$ | $\mathcal{U}[0.5, 1.5]$ | Dist. of signal profile parameter $a_2 = \lim_{t\to-\infty} s_2(t)$. |
| $\pi(b_1)$ | $\mathcal{U}[-1.0, 1.0]$ | Dist. of signal profile parameter $b_1 = \lim_{t\to\infty} s_1(t)$. |
| $\pi(b_2)$ | $\mathcal{U}[-0.5, 0.5]$ | Dist. of signal profile parameter $b_2 = \lim_{t\to\infty} s_2(t)$. |
| $\pi(t_{c1})$ | $\mathcal{U}[0.1T, 0.15T]$ | Dist. of signal profile parameter $t_{c1}$. |
| $\pi(t_{c2})$ | $\mathcal{U}[0.1T, 0.15T]$ | Dist. of signal profile parameter $t_{c2}$. |
| $\pi(r_1)$ | $\ln r_1 \sim \mathcal{U}[3, 4]$ | Dist. of signal profile parameter $r_1$. |
| $\pi(r_2)$ | $\ln r_2 \sim \mathcal{U}[3, 4]$ | Dist. of signal profile parameter $r_2$. |
| $N_{\text{samps}}$ | $1 + (T/\Delta T) = [5, 3, 2]$ | Number of sampling timepoints per experiment. |
| $\lvert\mathcal{D}_{\text{train}}\rvert$ | $N_{\text{exps}} \cdot (N_{\text{samps}} - 1) = 200$ | Total number of training datapoints. |

Table S7: Parameters used to generate a series of three synthetic training datasets, in which the sampling interval $\Delta T$ was varied in order to assess the sensitivity of landscape inference to the sampling rate. The ground-truth model $\mathcal{M}_1^*$ given by (S30) was used to generate the data in each case. The bracketed values shown for $N_{\text{exps}}$, $\Delta T$, and $N_{\text{samps}}$ correspond to each of the three datasets.

| Parameter | Value | Description |
|---|---|---|
| $\dim(\boldsymbol{\Phi})$ | $[2, 16, 32, 32, 16, 1]$ | $\boldsymbol{\Phi}^{nn}$ layer sizes from input to output. |
| $\alpha_{\boldsymbol{\Phi}}^h$ | `softplus` | Activation function for the hidden layers of $\boldsymbol{\Phi}^{nn}$. |
| $\alpha_{\boldsymbol{\Phi}}^f$ | `identity` | Activation function for the final layer of $\boldsymbol{\Phi}^{nn}$. |
| $C_{\text{conf}}$ | 0.1 | Confinement factor. |
| $\sigma_0$ | 0.05 | Initial value of noise parameter $\sigma$. |
| $\boldsymbol{\Phi}_W^0$ | `uniform xavier` | Initialization scheme for the weights of $\boldsymbol{\Phi}^{nn}$. |
| $\boldsymbol{\Phi}_b^0$ | 0.0 | Initialization scheme for the biases of $\boldsymbol{\Phi}^{nn}$. |
| $\mathbf{A}^0$ | `uniform xavier` | Initialization scheme for the weights of $\boldsymbol{\Psi}$. |

Table S8: Hyperparameters used to define the architecture and initialization of the PLNNs trained in the sampling rate sensitivity analysis.

| Parameter | Value | Description |
|---|---|---|
| $N_{\text{epochs}}$ | 2000 | Number of training epochs. |
| `patience` | 100 | Number of epochs before early halting. |
| `min_epochs` | 500 | Minimum number of required epochs. |
| $B$ | 250 | Batch size. |
| $N_{\text{cells}}$ | 200 | Number of cells simulated in each ensemble. |
| `solver` | `heun` | SDE Solver method. |
| $\Delta t$ | $0.1, [200 : 0.5, 500 : 0.5, 1000 : 0.5]$ | Timestep schedule (initial, [epoch:reduction]). |
| $\mathcal{L}$ | $MMD$ | Loss function. |
| `kernel` | Gaussian | MMD kernel function. |
| $\gamma$ | $[0.2, 0.5, 0.9, 1.3]$ | kernel bandwidth parameter(s). |
| `optimizer` | `RMSProp` | Optimizer algorithm. |
| `momentum` | 0.5 | Optimizer hyperparameter. |
| `weight decay` | 0.9 | Optimizer hyperparameter. |
| `clipping` | 1.0 | Gradient clipping hyperparameter. |
| `lr` | `exponential`$[10^{-2}, 10^{-5}, 50]$ | Learning rate schedule [initial, final, warmup]. |

Table S9: Hyperparameters used in the training of PLNNs for the sampling rate sensitivity analysis.

### S5 Application: An *in vitro* model of stem cell differentiation

In this section we apply our model to the *in vitro* system originally presented by Sáez et al. [1], concerning the differentiation of mouse embryonic stem cells (mESCs) into either a neural or mesoderm fate. In Section S5A we summarize the *in vitro* system and recapitulate the cell type labeling procedure of the original work. In Section S5B we use the assigned cell type labels to identify and isolate those cells involved in each of the two fate decisions. Section S5C details the dimension reduction and projection process used to identify a two-dimensional decision manifold on which we infer a landscape model. The training details are provided in Section S5D, and results in Section S5E.

#### S5A Cell type identification (recapitulation)

In Sáez et al. [1], the authors conducted a number of *in vitro* experiments, in which a population of mouse epiblast-like (EPI) stem cells was exposed to varying levels of two chemical morphogens, FGF and WNT. The level of FGF in the system was controlled by direct application and withdrawal of exogenous FGF2 (FGF), and by application of the small-molecule PD0325901 (PD), which inhibited downstream endogenous FGF production. The level of WNT signaling was controlled by application of the indirect WNT activator CHIRON99021 (CHIR). Different timecourses of CHIR, FGF, and PD resulted in particular patterns of cell differentiation. For a full discussion of how the particular combinations and durations of the signals map to cell fate outcomes, we refer the reader to the original work.

Over the course of five days, in response to the particular prescribed signal profile, the EPI cells took on either an anterior neural progenitor (AN) state, or posteriorized to acquire a caudal epiblast (CE) identity, thereby undergoing an initial binary decision. In a second binary decision, CE cells subsequently differentiated into either a posterior neural progenitor (PN) or paraxial mesoderm (M) state. The ultimate and essential observation is that the pattern of differentiation in the first binary decision is indicative of a binary choice, while the second is reminiscent of a binary flip (see original work for details).

At 12-hour intervals between and including days 2 and 5 (D2-5), the cells were profiled using a flow-cytometry assay that measured the expression of five marker proteins: BRA, CDX2, SOX1, SOX2, and TBX6. In addition, the assay measured OTX2 and FOXC2, though the previous five proteins were deemed sufficient to distinguish cell types. In this five-dimensional space, six cell states were identified using a clustering methodology based on fitting a Gaussian Mixture Model (GMM) at each timepoint. The identified states included the EPI, AN, CE, PN, and M states, as well as a transitioning (Tr) population between the EPI and CE states. Cells not confidently associated with any of these six states were labeled as *unidentified transitioning* (UT). Here, "confidently associated" implies that a cell had a posterior probability of at least $p^* = 0.65$ of belonging to a component of the fitted GMM. The authors labeled each cell in the dataset, and constructed for each signal profile a sequence of distributions (one per timepoint) detailing the time evolution of the distribution of cells across types.

An initial experimental series consisted of 11 conditions, each defined by a particular signaling timecourse. These conditions are shown in Table S10 and Fig. S7. The temporal sequence of cell type distributions, as reported in the original work, is depicted in Fig. S8. The results of our recapitulation of this clustering algorithm are shown in Figs. S9 and S10. In the latter, we have applied a correction to the cells identified as AN and PN at times D4.5 and D5, as is done in the original work. This correction is necessary because at these times the distinction between AN and PN cells is lost, and it is assumed that since no EPI cells remain at D4 to transition to the AN state, any subsequent appearance of neural cells indicates an increase in PN cells [1]. These nascent PN cells, however, may be classified by the clustering algorithm as AN, due to the similarity between the cell types at this stage. Therefore, for times D4.5 and D5, the authors applied a correction as follows. First, they recorded the number of AN cells at D4, which we'll denote by $\#(\mathrm{AN}|4)$. Then, for a subsequent timepoint $t \in \{4.5,\ 5\}$, they found the total number of neural cells labeled either AN or PN, denoted $\#(\mathrm{N}|t)$. The corrected number of AN cells and PN cells at time $t$, $\#(\mathrm{AN}|t)$ and $\#(\mathrm{PN}|t)$, respectively, were then determined according to

$$\#(\mathrm{AN}|t) = \min\{\#(\mathrm{AN}|4),\ \#(\mathrm{N}|t)\}$$
$$\#(\mathrm{PN}|t) = \max\{\#(\mathrm{N}|t) - \#(\mathrm{N}|4),\ 0\}.$$

That is, the number of AN cells at a subsequent timepoint could not exceed the number found at D4, and any additional neural cells were assumed to be PN.

In the original work, this correction posed little issue, as the adjustment took place by simply adjusting the cell type proportions, and did not involve relabeling *individual* cells. In our case, however, in order to proceed we need the labels of individual cells, and the question becomes, for each of the days D4.5 and D5,

| Condition | CHIR | FGF | PD |
|---|---|---|---|
| **NO CHIR** | NA | $(0.0, 3.0)$ | NA |
| CHIR 2-2.5 | $(2.0, 2.5)$ | $(0.0, 3.0)$ | NA |
| **CHIR 2-3** | $(2.0, 3.0)$ | $(0.0, 3.0)$ | NA |
| CHIR 2-3.5 | $(2.0, 3.5)$ | $(0.0, 3.0)$ | NA |
| CHIR 2-4 | $(2.0, 4.0)$ | $(0.0, 3.0)$ | NA |
| **CHIR 2-5** | $(2.0, 5.0)$ | $(0.0, 3.0)$ | NA |
| CHIR 2-5 FGF 2-3 | $(2.0, 5.0)$ | $(0.0, 3.0)$ | $(3.0, 5.0)$ |
| CHIR 2-5 FGF 2-3.5 | $(2.0, 5.0)$ | $(0.0, 3.5)$ | $(3.5, 5.0)$ |
| CHIR 2-5 FGF 2-4 | $(2.0, 5.0)$ | $(0.0, 4.0)$ | $(4.0, 5.0)$ |
| CHIR 2-5 FGF 2-4.5 | $(2.0, 5.0)$ | $(0.0, 4.5)$ | $(4.5, 5.0)$ |
| CHIR 2-5 FGF 2-5 | $(2.0, 5.0)$ | $(0.0, 5.0)$ | NA |

Table S10: The 11 experimental conditions making up the initial experimental series in Sáez et al. [1]. The boldfaced conditions consitute the reference set, which was used to train a GMM at each timepoint in order to identify cell type clusters. The columns depict the intervals of time between D0 and D5 during which the exogenous chemical signal was prescribed.

out of the combined population of neural cells, how do we determine which ones should be assigned to the AN state, and which should be assigned to the PN state?

Given the combined neural population consisting of cells initially labeled either AN or PN, we first assign the AN label to the nominal neural cells with the highest posterior probabilities corresponding to the AN component, assigning at most $\#(\text{AN}|4)$ cells to this label. Then, assuming that there is an excess of neural cells, the remaining are assigned the PN label.

| | **BRA** | **CDX2** | **SOX1** | **SOX2** | **TBX6** | OTX2 | FOXC2 |
|---|---|---|---|---|---|---|---|
| EPI | | | | ++ | | + | |
| Tr | | | | + | | + | |
| AN | | | + | ++ | | + | |
| CE | (+) | + | | | | | |
| PN | | (+) | ++ | + | | | |
| EM | (+) | | | | + | | |
| LM | | | | | | | + |

Table S11: Reproduction of Figure 2D of Sáez et al. [1]. Marker proteins used to identify cell types. ++ denotes high levels, + denotes moderate levels, and (+) denotes an optional presence, based on transitioning populations. Cell identities can be determined by the core set of 5 markers, shown in bold.

### S5B Isolating two binary decisions

Having recapitulated the cell type identification, we now turn our attention to each of the two binary decisions in turn. Our goal here is to isolate the dynamics of the two decisions individually, and then infer a PLNN that captures the 3-attractor landscape dynamics of that particular decision. This approach stands in contrast to attempting to infer a 5-attractor system capturing the entire system as a whole. The difficulty with the latter is that our model, by construction, assumes a global effect of signals. That is, the effect of a signal is to tilt the *entire* landscape in a particular direction, thus biasing the movement of cells uniformly. Our core assumption is that the dynamics of a binary cell fate decision can be captured in a two-dimensional linear manifold, or plane, within gene expression space. In the case of one binary decision followed by another, however, we do not believe that the same plane serves to capture the dynamics of both decisions.

Let us note here that in Sáez et al. [1], the authors do indeed infer a 5-attractor landscape, by partitioning the plane into two regions, each with its own mapping of signals to tilts. In this way, they infer two signal processing functions, one for each binary decision. The inference of a more complex landscape—involving more than three attractors—is a natural route of future investigation. For now, however, we will make clear that the present goal is to infer a 3-attractor landscape for each of the two decisions, directly from gene expression data.

We start with the first decision, in which EPI cells transition to either the AN or CE state. The transitioning (Tr) population should also be captured in this decision, as an intermediate state between EPI and CE, but

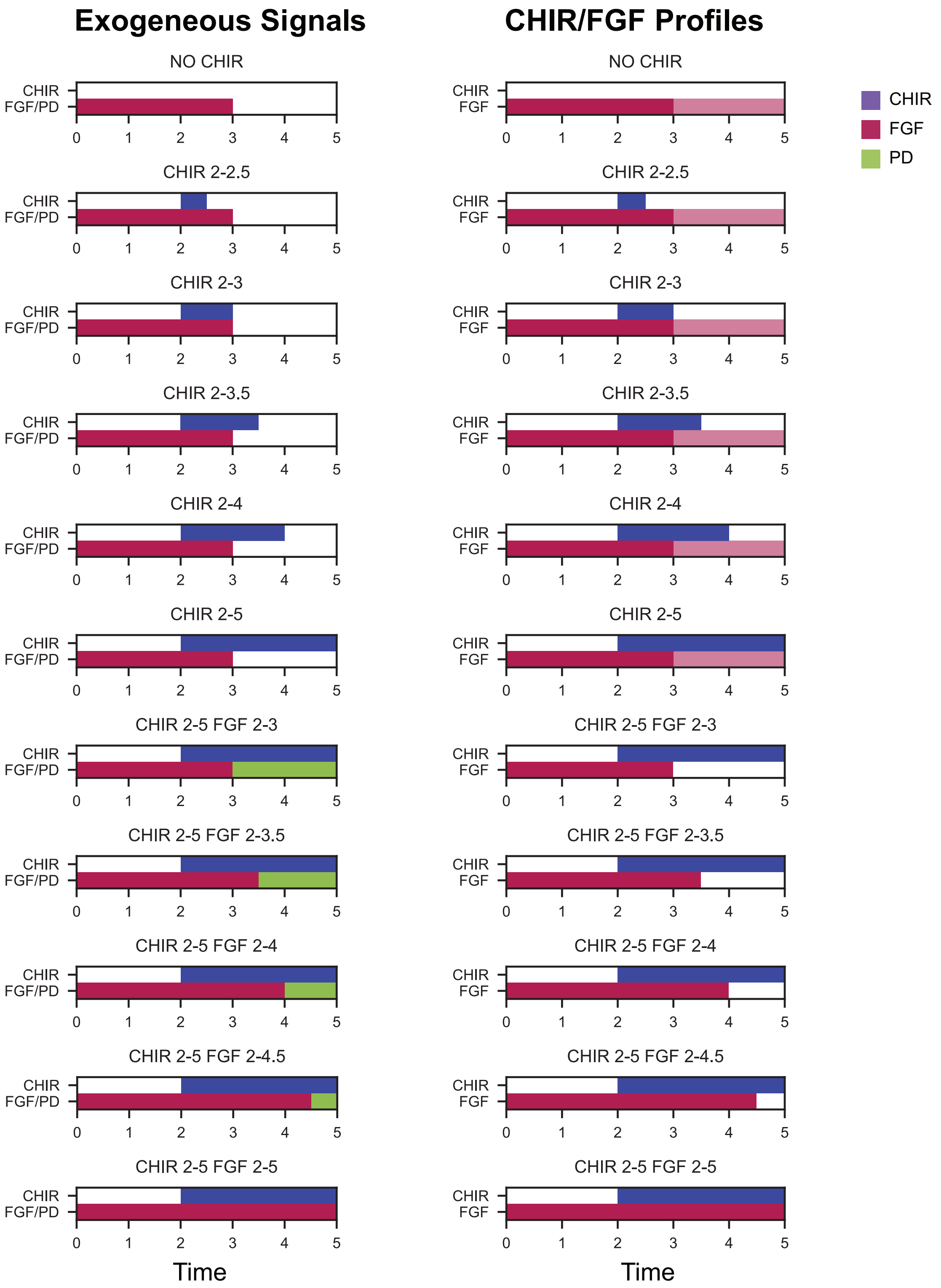

Figure S7: Recapitulation of the analysis of the initial experimental series conducted in Sáez et al. [1]. The light-red shading of the FGF profile indicates that a level of endogenously-produced FGF is assumed to be present in the system, and occurs when PD is not prescribed to block this production.

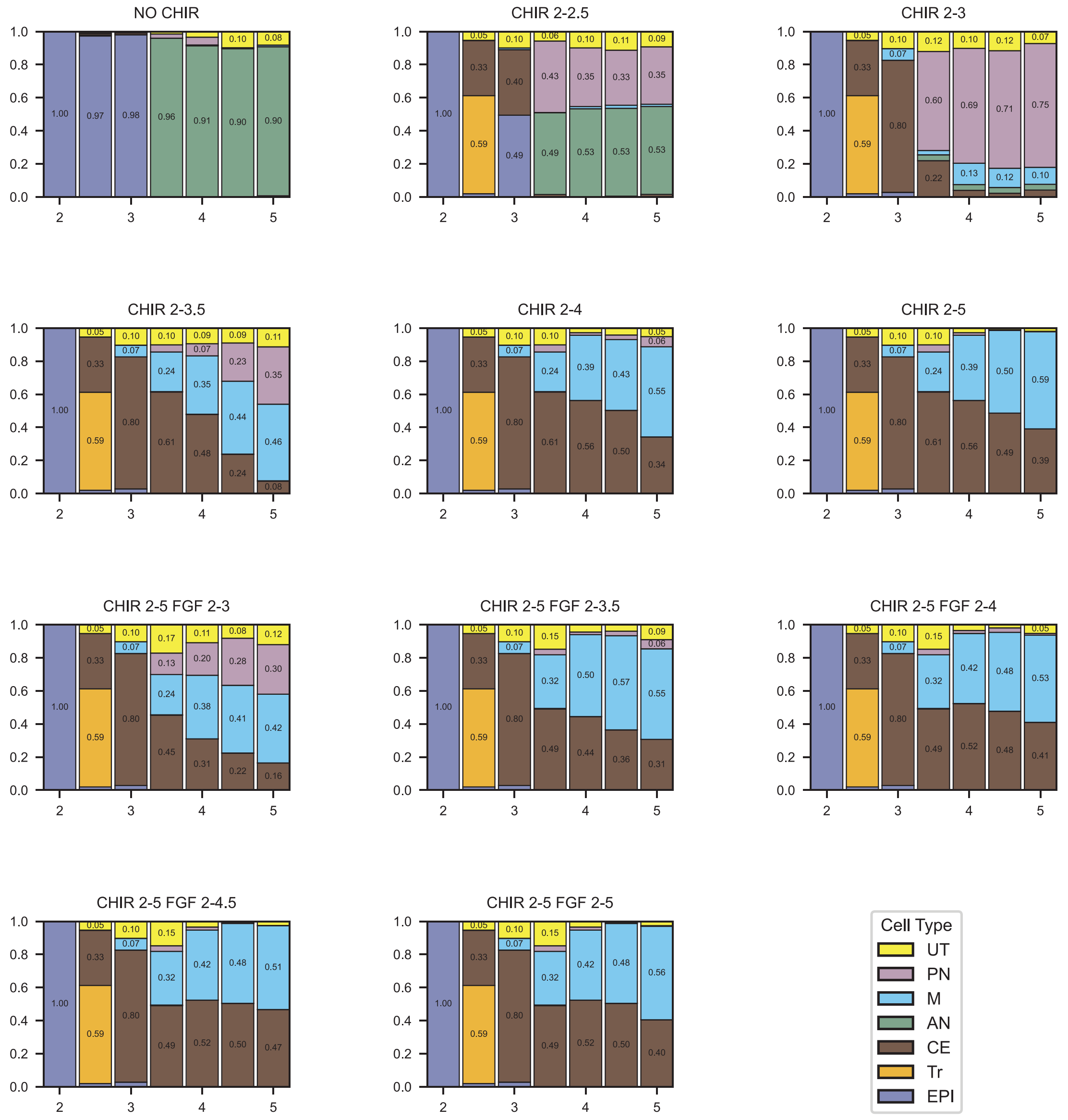

Figure S8: Results of the clustering algorithm performed on the initial experimental series as conducted by Sáez et al. [1]. For each of the 11 conditions in the initial experimental series, cells are clustered among six distinguishable cell types. Note that in Sáez et al. [1], the number of AN cells found at D4 was used as a maximum for the number of AN cells found at the following days (D4.5 and D5), so that if the number of cells labeled as AN at D4.5 or D5 exceeded the number found at D4, the difference was instead assumed to be indicative of incorrectly-classified PN cells, as no EPI cells remained at that point to transition to AN.

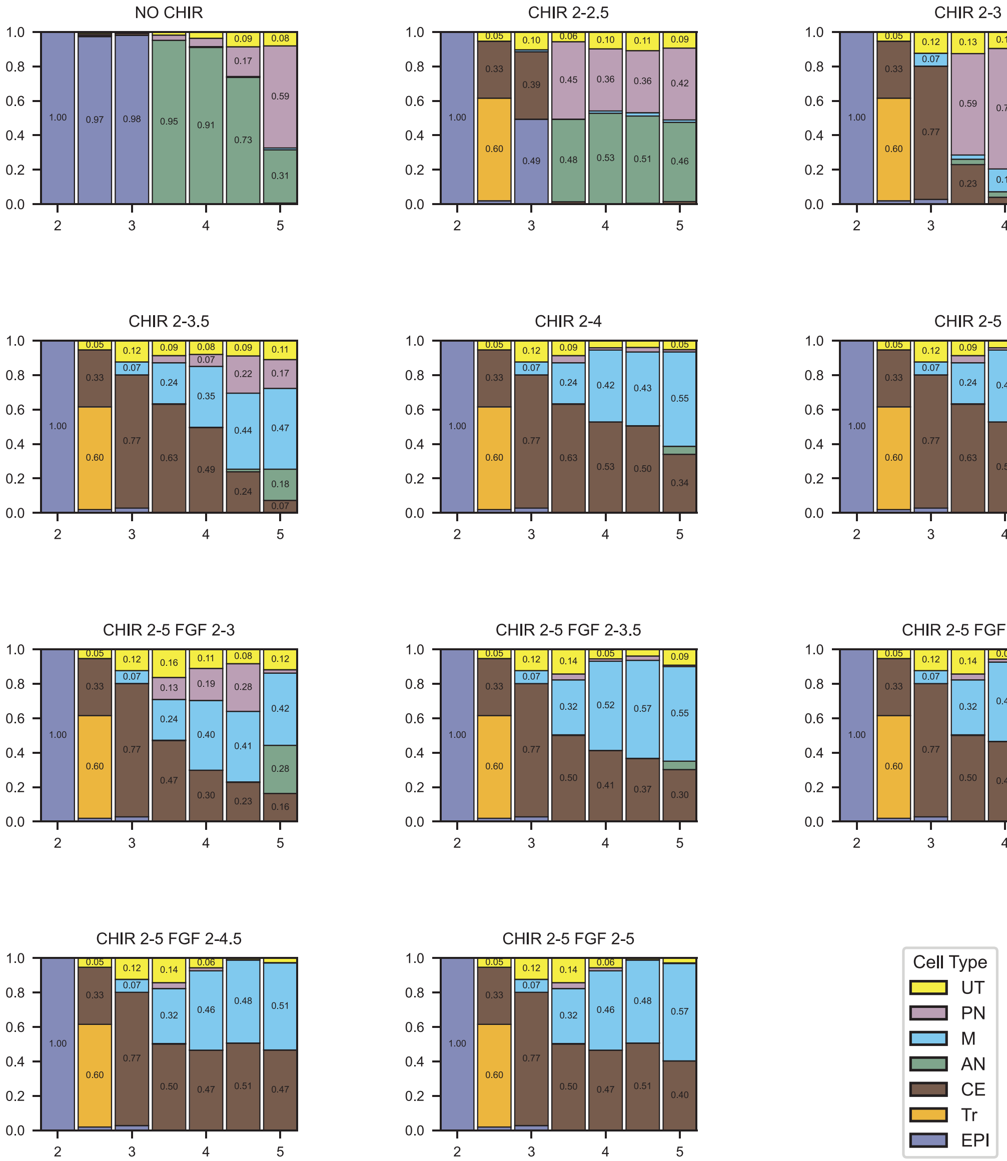

Figure S9: Recapitulation of the clustering algorithm performed on the initial experimental series conducted in Sáez et al. [1]. For each of the 11 conditions in the initial experimental series, cells are clustered among six distinguishable cell types. Our results match those of the original work, up to post hoc adjustment of the number of AN and PN cells for days 4.5 and 5.

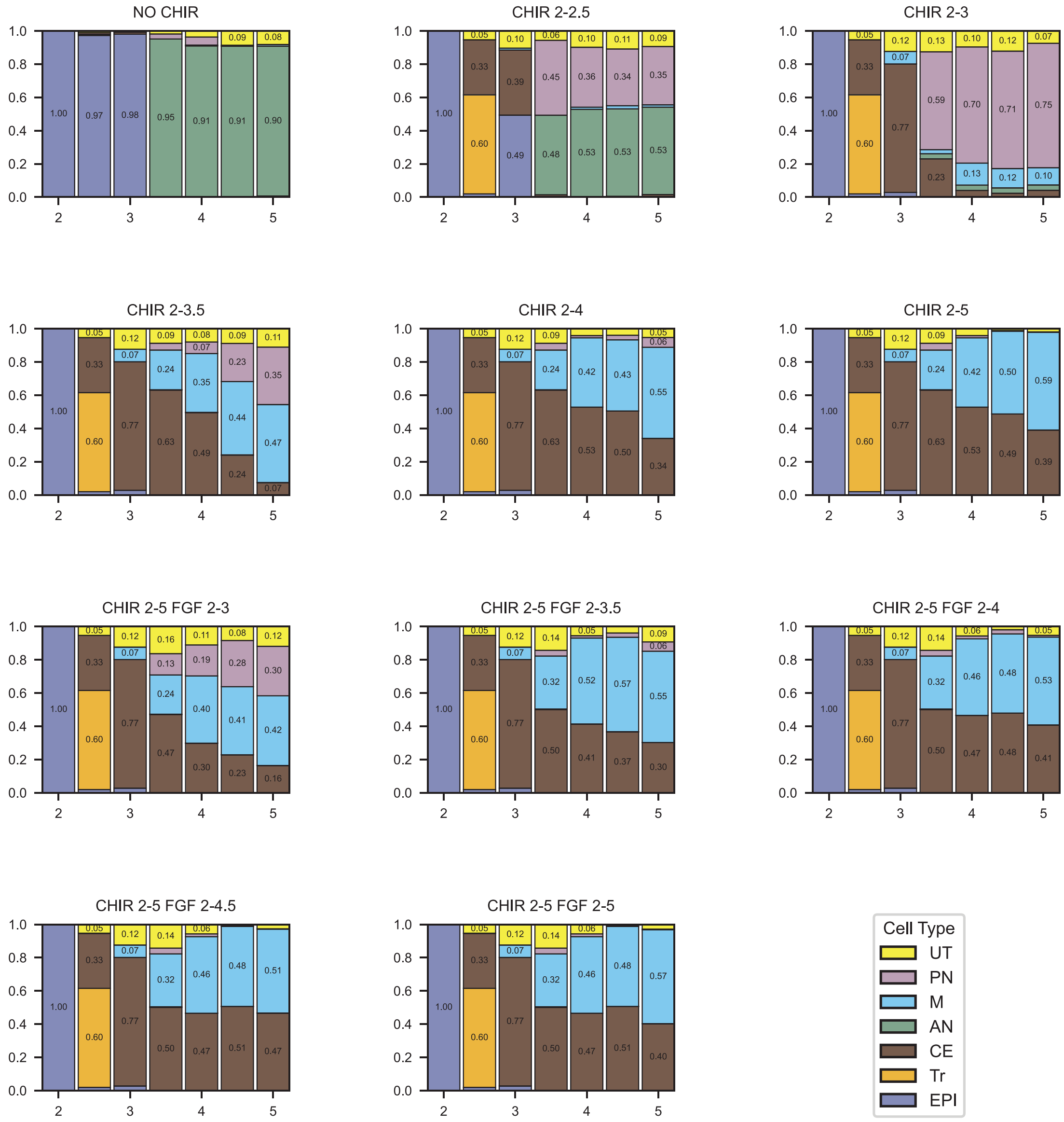

Figure S10: Recapitulation of the clustering algorithm performed on the initial experimental series conducted in Sáez et al. [1], with correction of AN and PN cells at D4.5 and D5.

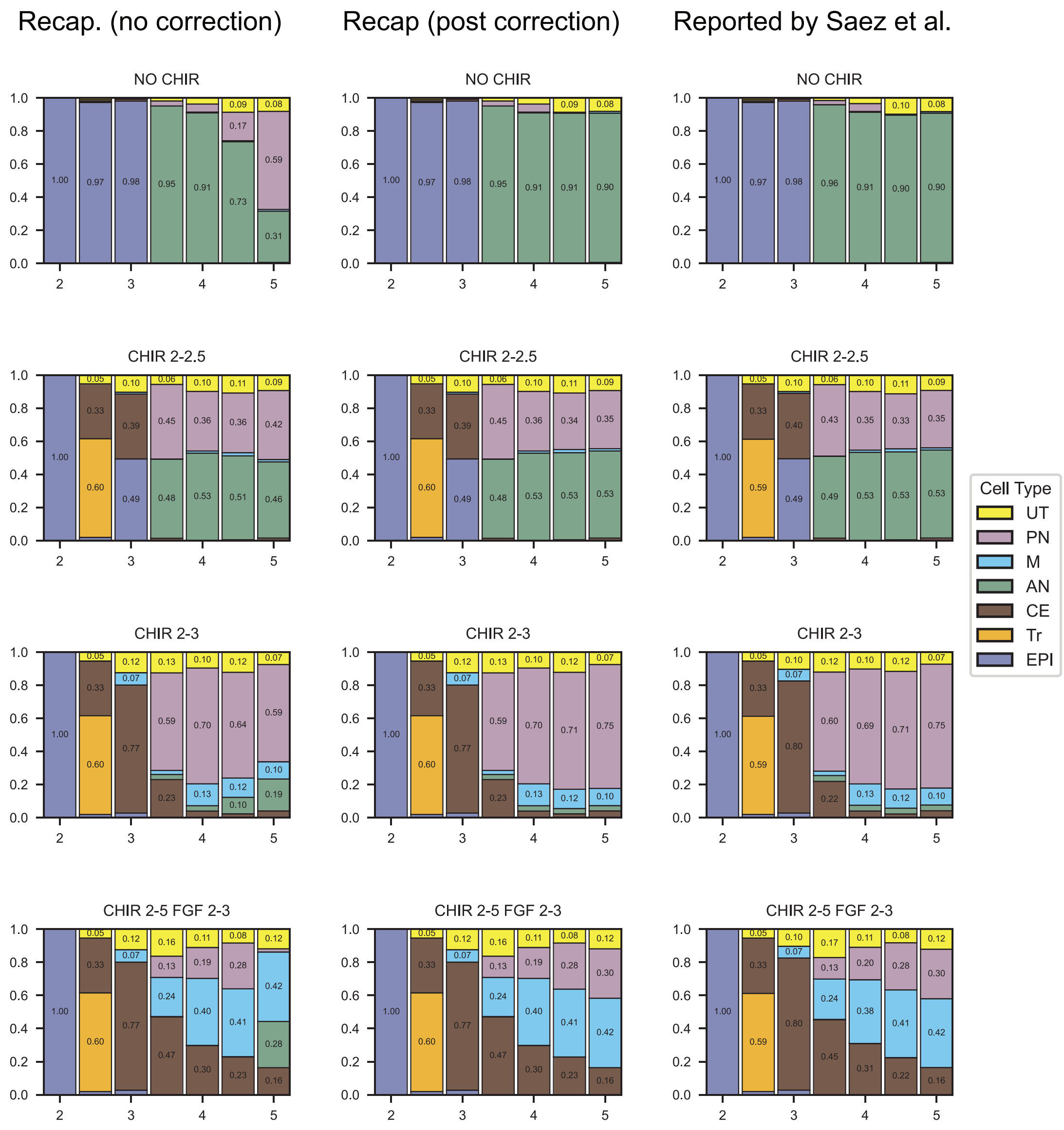

Figure S11: A side by side comparison of the cell type distributions as reported in Sáez et al. [1] (right column) with our recapitulation, before and after adjusting the number of neural cells (left and center columns, respectively).

we do not want to try to fit the model to capture the PN and M attractors. We cannot simply exclude cells in the later states, PN and M, as this would bias the distribution of cells over the landscape, overestimating the fraction of cells transitioning to the AN state relative to the CE state. The crux of the problem is that cells continue to differentiate, passing out of the CE attractor and into either the PN or M attractor. To accommodate the fact that cells leave the CE attractor, we count the number of cells labeled as either PN or M, and remove these from the landscape. Then, to make up for this removal, we add the same number of cells back to the landscape by generating pseudo-cells from the fitted GMM components corresponding to the CE attractor. The idea is that those cells labeled as either PN or M could only have differentiated from the CE state. Thus, we will approximate a distribution of cells where the subsequent transition from the CE state is not permitted, and cells instead remain in the location of the CE attractor. The best estimate we have of this attractor is the component (or components) of the fitted GMM corresponding to the CE state, which we know from the fitting procedure. In the case of multiple CE components in the fitted GMM, we respect the inferred weighting between them, and sample the corresponding fraction of cells to be replaced from each component's multivariate normal distribution. This removal, followed by subsequent re-population, is depicted in Fig. S12. Importantly, we have not only relabeled particular cells as CE, but have also generated new expression data for those cells.

We have a similar concern when it comes to the second decision. In this case, we would like to isolate the transition from the CE state to either the PN or M state, while somehow excluding the EPI, Tr, and AN states. We will exclude the data at D2 and D2.5, and use for the initial state of the system the time D3, at which point the data suggest that the remaining EPI cells will exclusively transition to the AN state. Thus, we can simply remove any remaining EPI and AN cells as part of the first decision. In addition, note that by D3 the Tr population is no longer present, so we need not be concerned about that particular state. The resulting data is shown in Fig. S13. The NO CHIR condition results in very few cells confidently identified as being part of the second decision, as for this signal profile cells transition to the AN state as opposed to the CE state. For this reason, we will exclude this condition when considering the second fate decision.

#### S5C Dimension reduction and projection

Once the data relevant to each decision is isolated, we perform a step of dimension reduction in order to acquire a 2-dimensional representation of each cell, in a space that will capture the dynamics that we are interested in. This step of dimension reduction is a necessary component that is currently done prior to and independent of the landscape inference, but can in theory be done simultaneously, by learning a latent space in which the dynamics of interest take place. Here, though, we describe the independent dimension reduction applied to the cells of interest, for each of the two decisions. Ultimately, we infer a coordinate space using Principal Component Analysis (PCA) and project all cells onto this space. The resulting 2-dimensional data is then used as the input data for the model training procedure.

There are a number of choices that we must make when it comes to applying PCA. Perhaps most importantly, out of all of the cells identified as part of a particular decision, we choose to apply PCA on only a subset of these in order to infer the principal axes. Specifically, we subset the cells to include those at the beginning and end of the decision of interest. After the principal axes have been inferred from the subset, the remaining cells at the intermediate times can be projected onto those axes. For the first decision, we select only the cells sampled at D2 and D3.5, thereby capturing the initial population of EPI cells, and the two terminal states AN and CE. For the second decision, we select the cells at D3 and D5, capturing the initial CE state and the terminal PN and M states. The fact that in both cases we have three distinguishable cell types suggests a plane defined by the modes of these populations. By not including cells at the intermediate times, we hope to ensure that the variance across cells on which PCA is applied is primarily due to the existence of discernible populations, and that the resulting axes will capture the directions in gene expression space separating these groups. The resulting data, after dimension reduction, for the first and second decision are shown in Fig. S14 and S15, respectively.

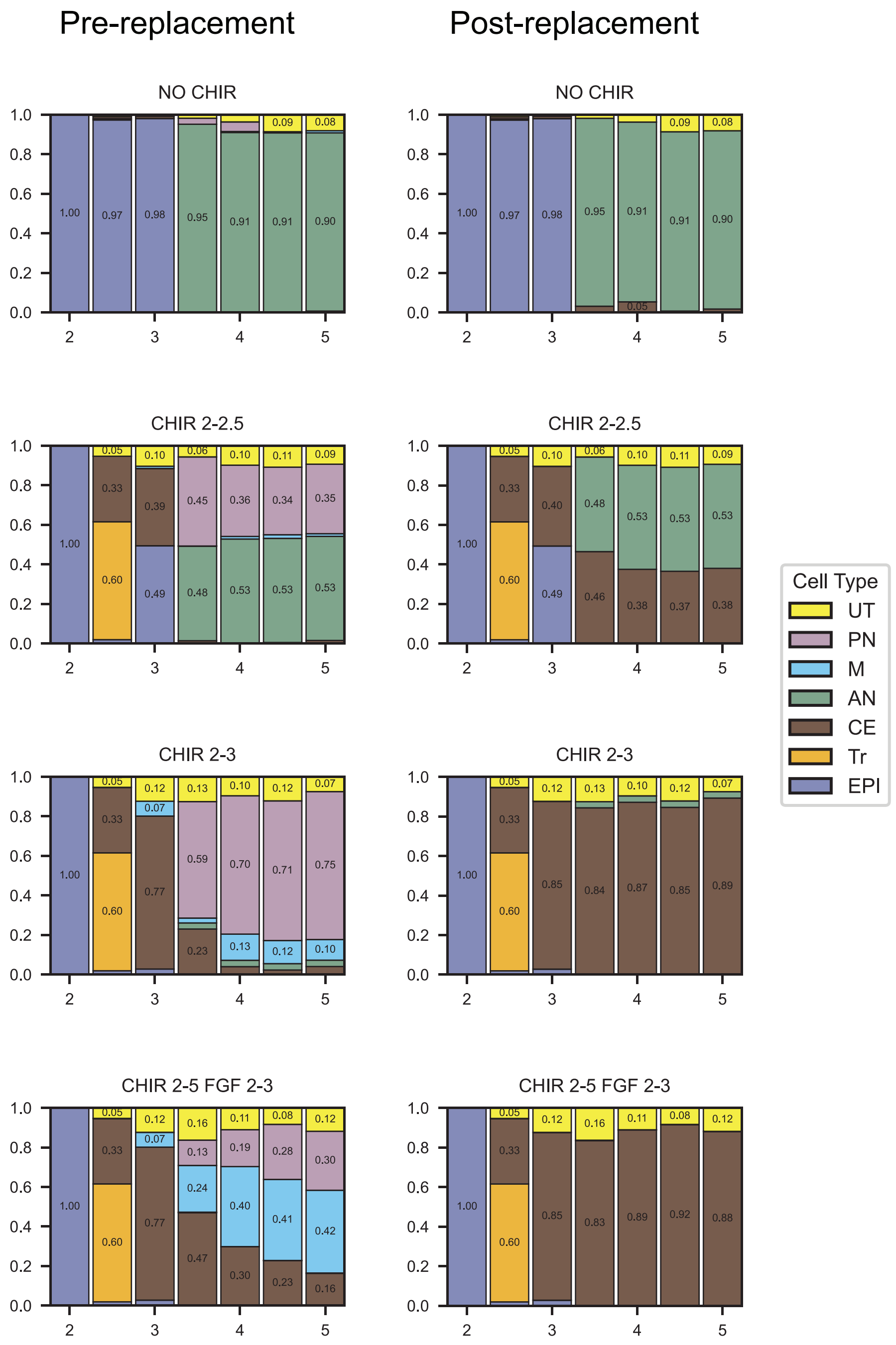

Figure S12: Isolation of the first binary decision, in which EPI cells either transition to the AN state or to the CE state. Starting with the labeled cells, at each timepoint we remove those labeled as PN or M. Then, to accommodate the resulting bias overestimating the fraction of cells labeled as AN relative to CE, we sample from the inferred CE attractors of the fitted GMM in order to repopulate the removed cells.

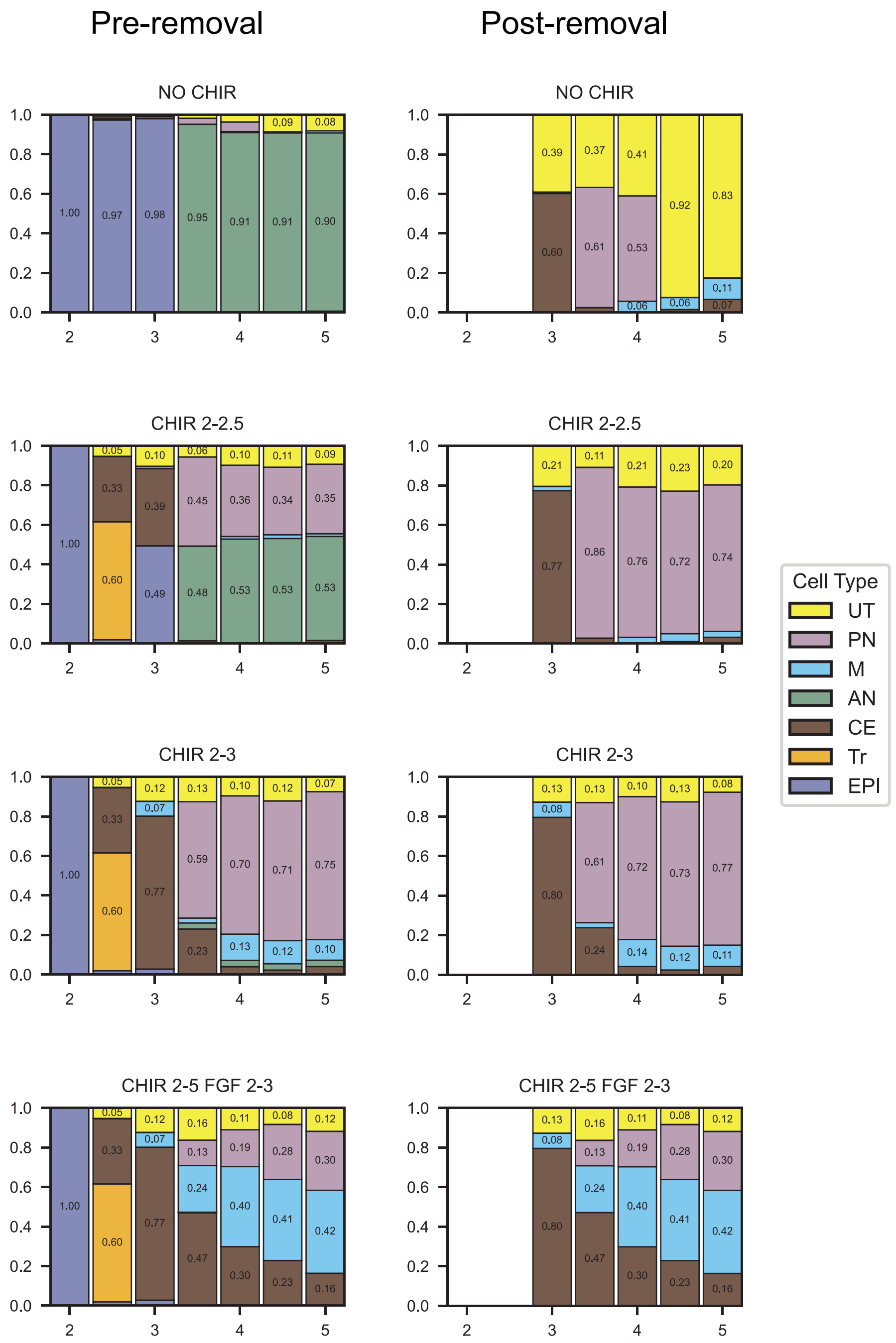

Figure S13: Isolation of the second binary decision, in which CE cells either transition to the PN state or to the M state. We restrict ourselves to the data beginning at D3, and remove any remaining EPI and AN cells. Note that the NO CHIR condition results in almost no cells involved in the second binary decision.

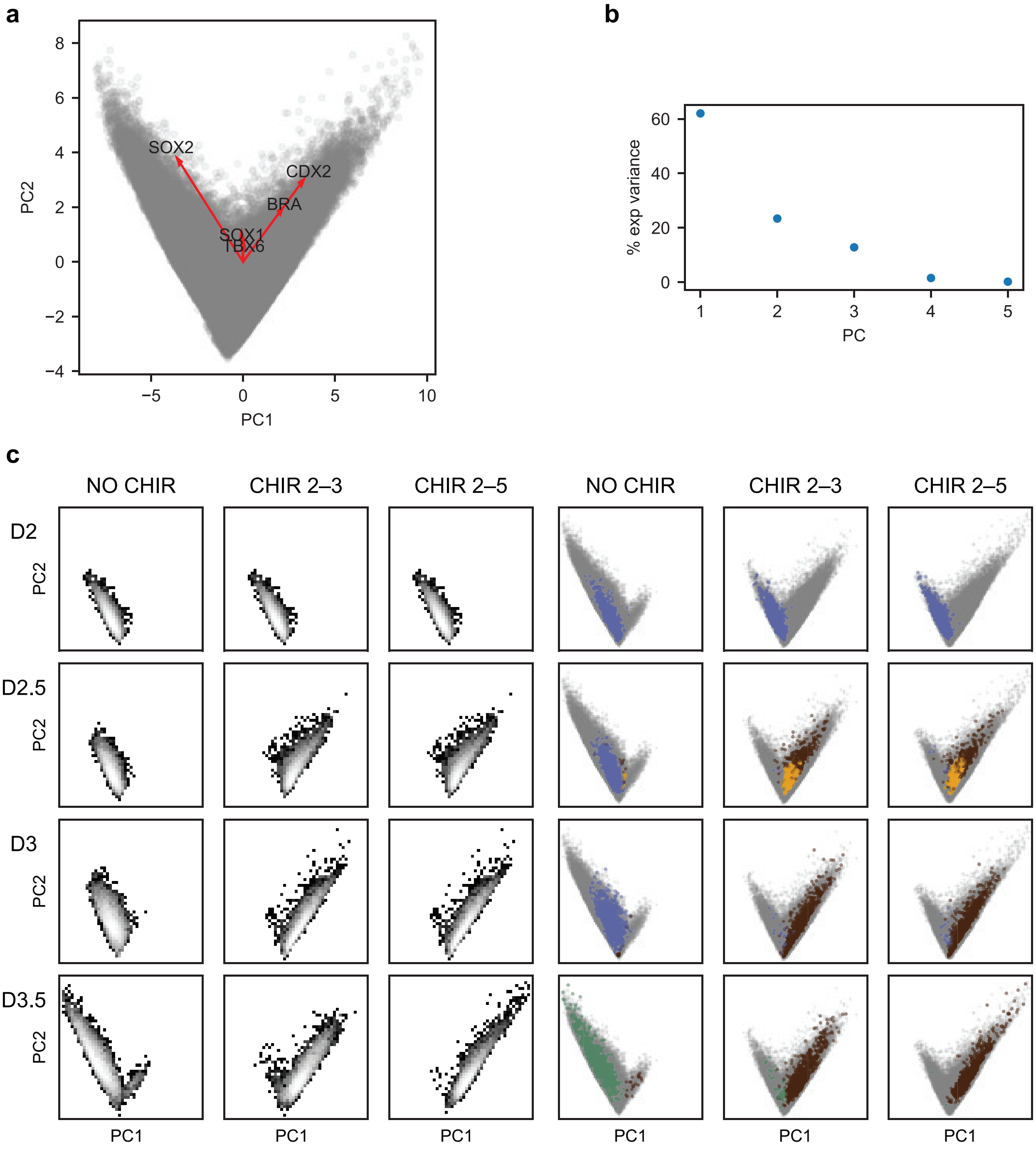

Figure S14: Dimension reduction applied to the cells identified as part of the first binary decision. **(a)** The data projected onto the first two principal components, with the loadings corresponding to each gene projected as well. **(b)** Scree plot showing the proportion of explained variance for each PC. **(c)** Temporal evolution of the distribution of cells for each experimental condition.

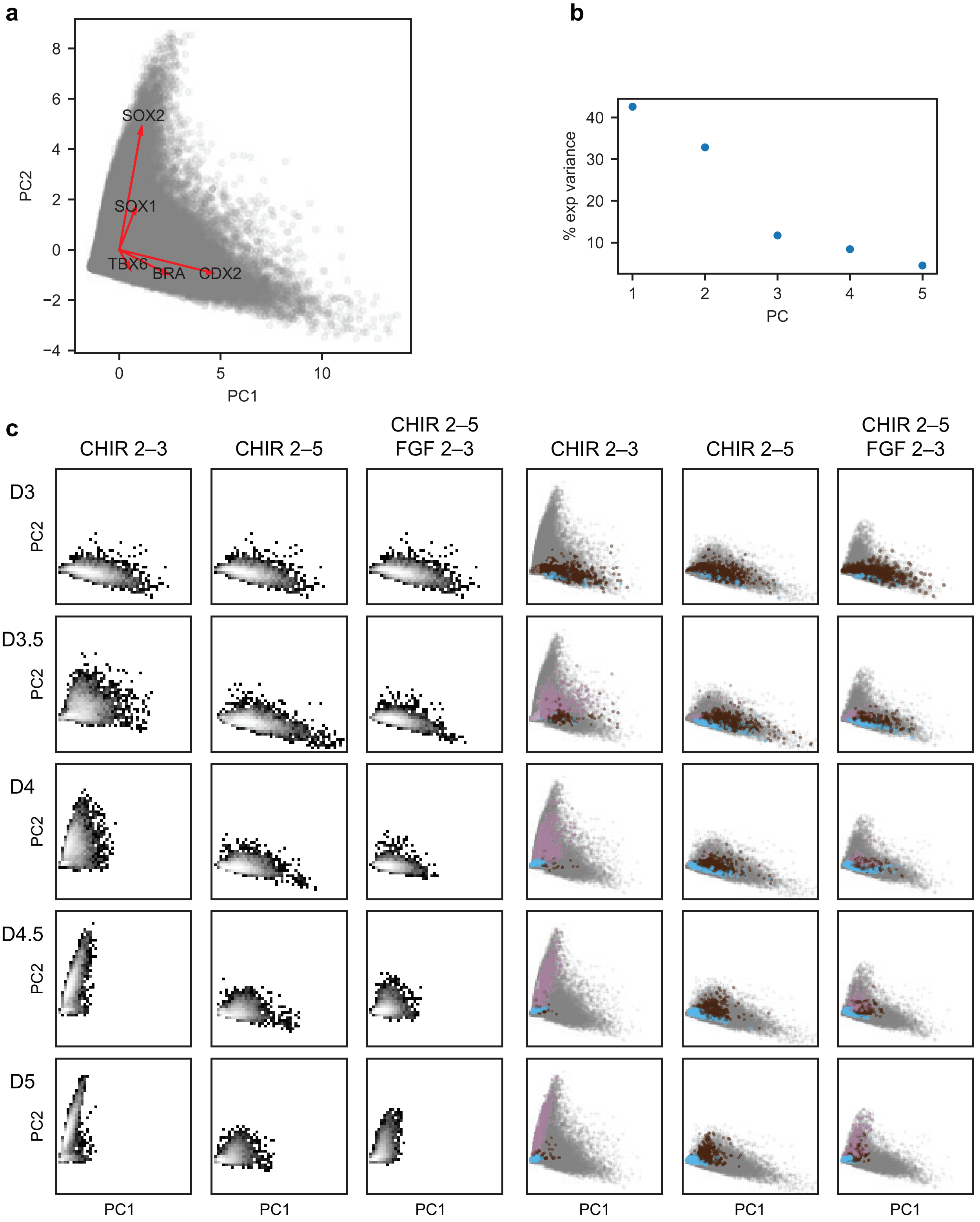

Figure S15: Dimension reduction applied to the cells identified as part of the second binary decision. **(a)** The data projected onto the first two principal components, with the loadings corresponding to each gene projected as well. **(b)** Scree plot showing the proportion of explained variance for each PC. **(c)** Temporal evolution of the distribution of cells for each experimental condition.

### S5D Model training

Tables S12 and S13 specify the model architecture and training hyperparameters used to infer the first *in vitro* decision. Tables S14 and S15 specify those used for the second decision.

| Parameter | Value | Description |
|---|---|---|
| $\dim(\mathbf{\Phi})$ | $[2, 16, 32, 32, 16, 1]$ | $\mathbf{\Phi}^{nn}$ layer sizes from input to output. |
| $\alpha_{\mathbf{\Phi}}^{h}$ | `softplus` | Activation function for the hidden layers of $\mathbf{\Phi}^{nn}$. |
| $\alpha_{\mathbf{\Phi}}^{f}$ | `identity` | Activation function for the final layer of $\mathbf{\Phi}^{nn}$. |
| $C_{\text{conf}}$ | 0.1 | Confinement factor. |
| $\sigma_0$ | 0.05 | Initial value of noise parameter $\sigma$. |
| $\mathbf{\Phi}_{W}^{0}$ | `uniform xavier` | Initialization scheme for the weights of $\mathbf{\Phi}^{nn}$. |
| $\mathbf{\Phi}_{b}^{0}$ | 0.0 | Initialization scheme for the biases of $\mathbf{\Phi}^{nn}$. |
| $\mathbf{A}^{0}$ | `uniform xavier` | Initialization scheme for the weights of $\mathbf{\Psi}$. |

Table S12: Hyperparameters used to define the architecture and initialization of the PLNN inferring the first *in vitro* binary decision landscape.

| Parameter | Value | Description |
|---|---|---|
| $N_{\text{epochs}}$ | 1000 | Number of training epochs. |
| `patience` | 200 | Number of epochs before early halting. |
| `min_epochs` | 10 | Minimum number of required epochs. |
| $B$ | 50 | Batch size. |
| `passes_per_epoch` | 10 | Times each datapoint is sampled per epoch. |
| $N_{\text{cells}}$ | 800 | Number of cells simulated in each ensemble. |
| `solver` | `heun` | SDE Solver method. |
| $\Delta t$ | $0.005, \{100, 300, 500, 700\} : 0.5$ | Timestep schedule (initial, [epoch:reduction]). |
| $\mathcal{L}$ | $MMD$ | Loss function. |
| `kernel` | Gaussian | MMD kernel function. |
| $\gamma$ | [0.2, 0.5, 1.0, 1.673845] | kernel bandwidth parameter(s). |
| `optimizer` | `RMSProp` | Optimizer algorithm. |
| `momentum` | 0.5 | Optimizer hyperparameter. |
| `weight decay` | 0.9 | Optimizer hyperparameter. |
| `clipping` | 1.0 | Gradient clipping hyperparameter. |
| `lr` | `exponential`$[10^{-2}, 10^{-5}, 50]$ | Learning rate schedule [initial, final, warmup]. |

Table S13: Hyperparameters used for the training of the PLNN inferring the first *in vitro* binary decision landscape.

| Parameter | Value | Description |
| --- | --- | --- |
| $\dim(\mathbf{\Phi})$ | $[2, 16, 32, 32, 16, 1]$ | $\mathbf{\Phi}^{nn}$ layer sizes from input to output. |
| $\alpha^h_{\mathbf{\Phi}}$ | `softplus` | Activation function for the hidden layers of $\mathbf{\Phi}^{nn}$. |
| $\alpha^f_{\mathbf{\Phi}}$ | `identity` | Activation function for the final layer of $\mathbf{\Phi}^{nn}$. |
| $C_{\text{conf}}$ | 0.1 | Confinement factor. |
| $\sigma_0$ | 0.05 | Initial value of noise parameter $\sigma$. |
| $\mathbf{\Phi}^0_W$ | `uniform xavier` | Initialization scheme for the weights of $\mathbf{\Phi}^{nn}$. |
| $\mathbf{\Phi}^0_b$ | 0.0 | Initialization scheme for the biases of $\mathbf{\Phi}^{nn}$. |
| $\mathbf{A}^0$ | `uniform xavier` | Initialization scheme for the weights of $\mathbf{\Psi}$. |

Table S14: Hyperparameters used to define the architecture and initialization of the PLNN inferring the second *in vitro* binary decision landscape.

| Parameter | Value | Description |
| --- | --- | --- |
| $N_{\text{epochs}}$ | 1000 | Number of training epochs. |
| `patience` | 200 | Number of epochs before early halting. |
| `min_epochs` | 10 | Minimum number of required epochs. |
| $B$ | 50 | Batch size. |
| `passes_per_epoch` | 10 | Times each datapoint is sampled per epoch. |
| $N_{\text{cells}}$ | 800 | Number of cells simulated in each ensemble. |
| `solver` | `heun` | SDE Solver method. |
| $\Delta t$ | $0.005, \{100, 300, 500, 700\} : 0.5$ | Timestep schedule (initial, [epoch:reduction]). |
| $\mathcal{L}$ | $MMD$ | Loss function. |
| `kernel` | Gaussian | MMD kernel function. |
| $\gamma$ | [0.2, 0.5, 1.0, 1.275068] | kernel bandwidth parameter(s). |
| `optimizer` | `RMSProp` | Optimizer algorithm. |
| `momentum` | 0.5 | Optimizer hyperparameter. |
| `weight decay` | 0.9 | Optimizer hyperparameter. |
| `clipping` | 1.0 | Gradient clipping hyperparameter. |
| `lr` | `exponential`$[10^{-2}, 10^{-5}, 50]$ | Learning rate schedule [initial, final, warmup]. |

Table S15: Hyperparameters used for the training of the PLNN inferring the second *in vitro* binary decision landscape.

### S5E Inferred decision landscapes

The results of the training process for the first and second *in vitro* decisions are shown in Figs. S16 and S17 (decision 1) and Figs. S18 and S19 (decision 2).

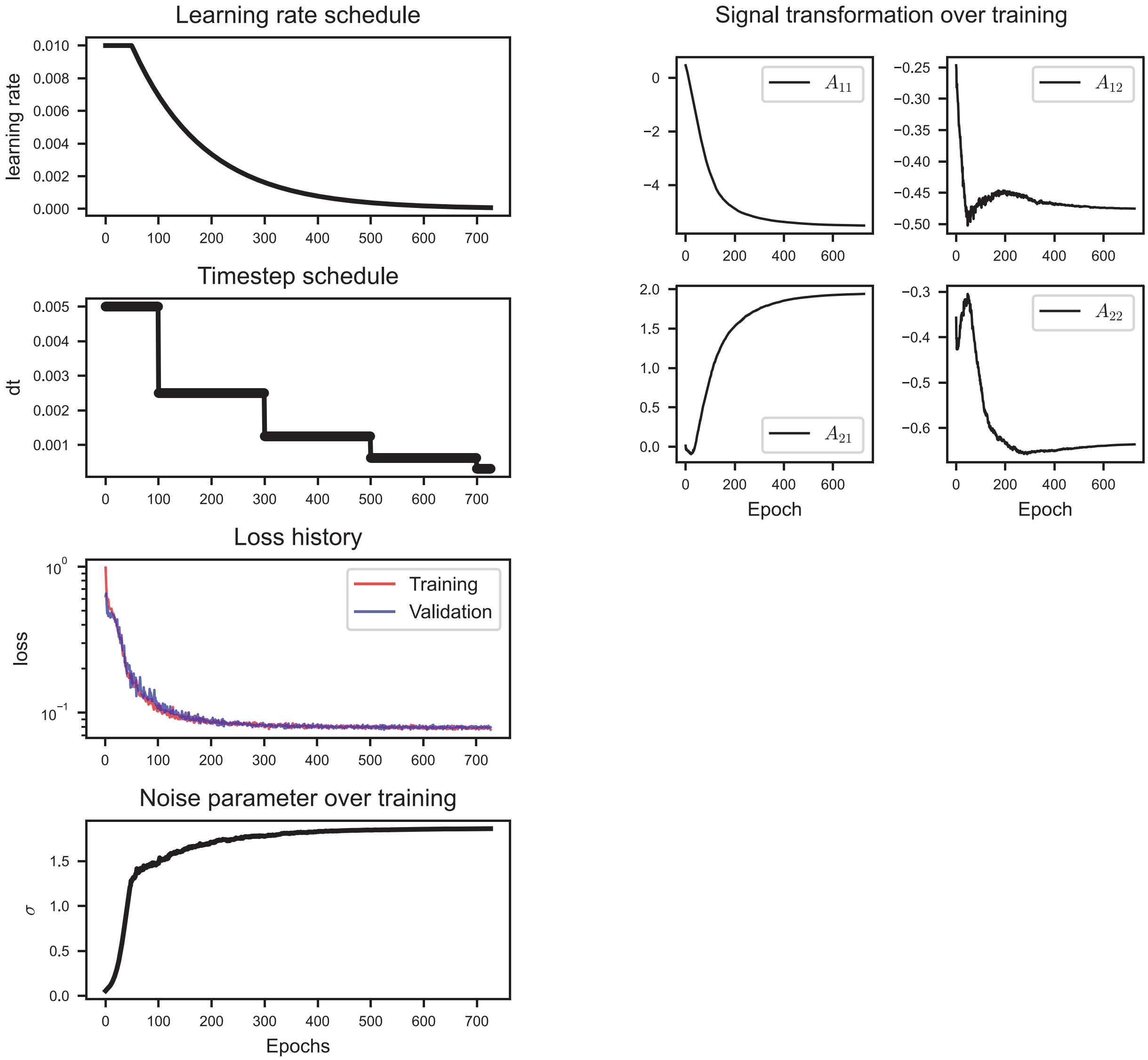

Figure S16: Details of the training process for the PLNN inferring the first *in vitro* decision.

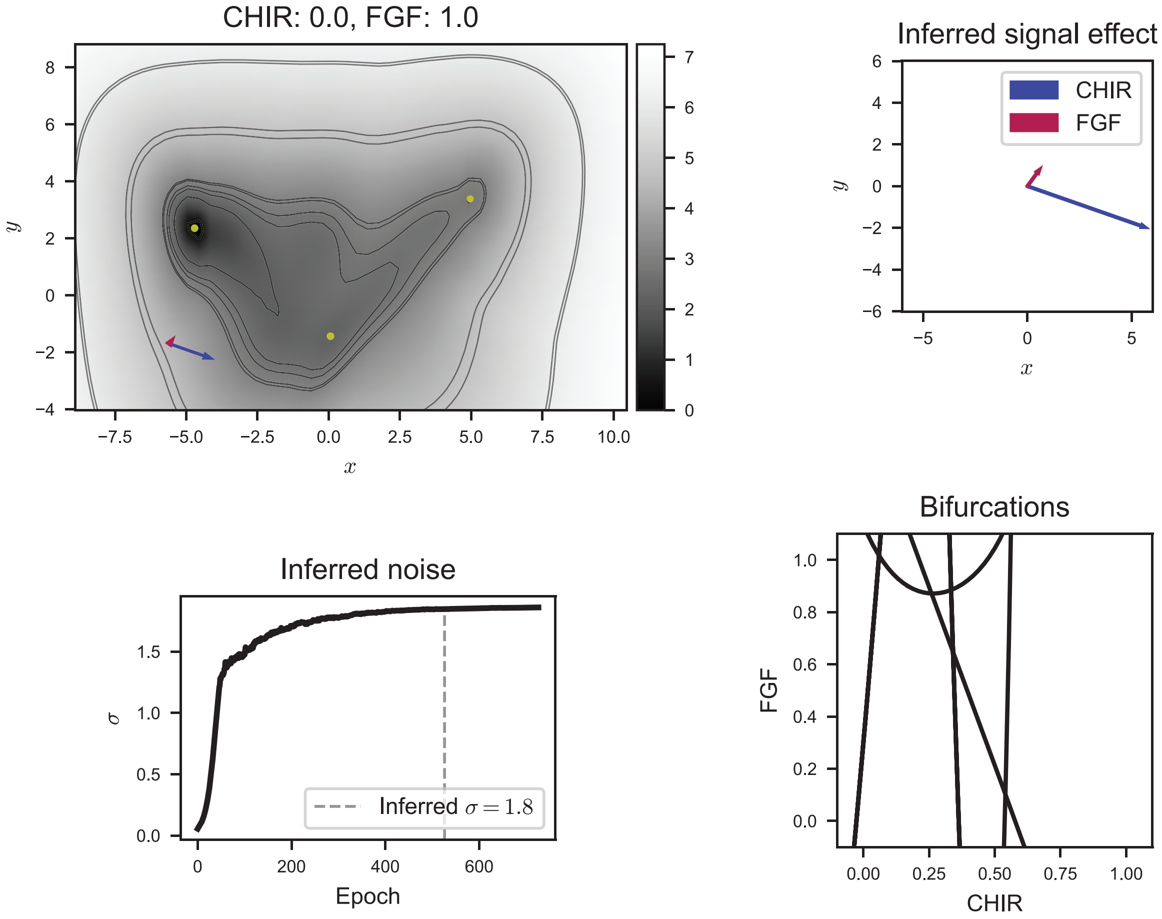

Figure S17: The inferred binary decision landscape corresponding to the first *in vitro* decision.

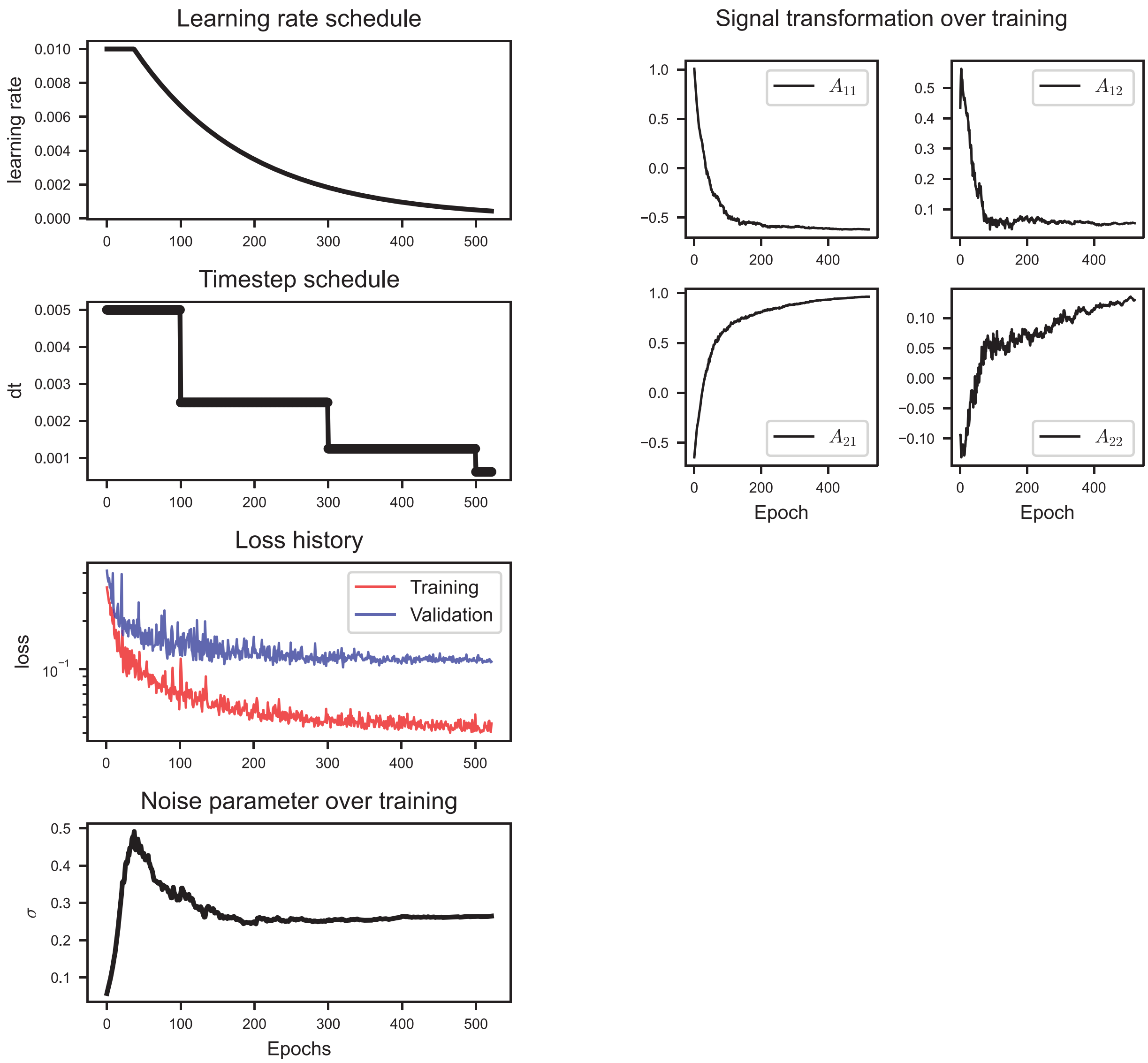

Figure S18: Details of the training process for the PLNN inferring the second *in vitro* decision.

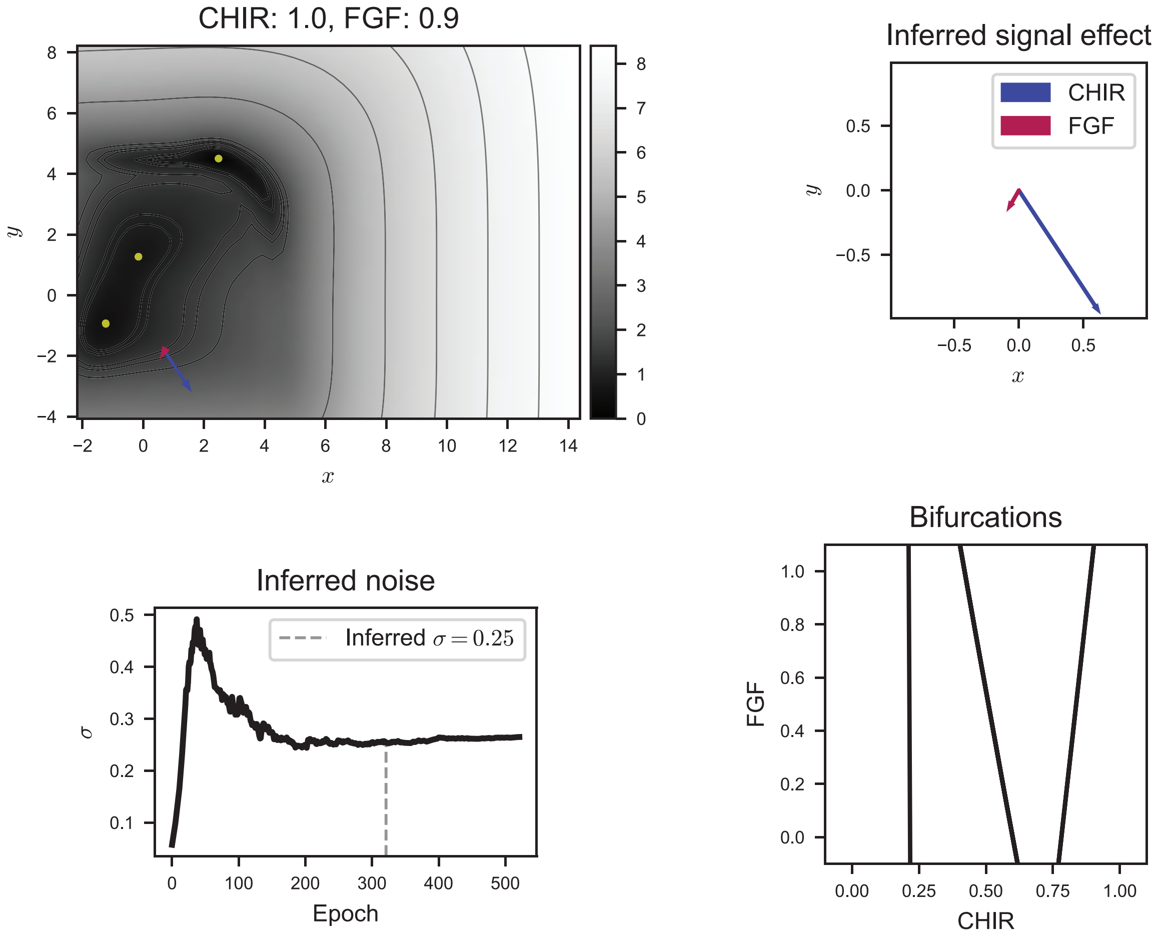

Figure S19: The inferred binary decision landscape corresponding to the second *in vitro* decision.

### SA Tracing fold curves

We employ an arclength continuation algorithm [38–41] to trace the locus of saddle-node bifurcations exhibited by the dynamics

$$\dot{\boldsymbol{x}} = \boldsymbol{F}(\boldsymbol{x}, \boldsymbol{p}) \tag{S33}$$

induced by a parameterized landscape, where here we refer to the landscape parameters with $\boldsymbol{p}$. The goal is to trace curves $(\boldsymbol{x}(s), \boldsymbol{p}(s))$ of saddle-node bifurcations, parameterized by a scalar $s$, in a combined $(\boldsymbol{x}, \boldsymbol{p})$ phase-parameter space of dimension $\mathbb{R}^{d+k}$, where we assume the system is two-dimensional and governed by two parameters, $d = k = 2$. Writing $\boldsymbol{x} = \boldsymbol{x}(s)$ and $\boldsymbol{p} = \boldsymbol{p}(s)$, we define the tangent (row) vector

$$\gamma(s) = (\dot{\boldsymbol{x}}(s), \dot{\boldsymbol{p}}(s)) := \left(\frac{d\boldsymbol{x}(s)}{ds}, \frac{d\boldsymbol{p}(s)}{ds}\right). \tag{S34}$$

Since $\boldsymbol{F}(\boldsymbol{x}(s), \boldsymbol{p}(s)) = 0$ for all $s$,

$$0 = \frac{d}{ds}\boldsymbol{F}(\boldsymbol{x}(s), \boldsymbol{p}(s)) = \boldsymbol{F}_{\boldsymbol{x}}\dot{\boldsymbol{x}} + \boldsymbol{F}_{\boldsymbol{p}}\dot{\boldsymbol{p}} = [\boldsymbol{F}_{\boldsymbol{x}} \mid \boldsymbol{F}_{\boldsymbol{p}}] \begin{bmatrix} \dot{\boldsymbol{x}} \\ \dot{\boldsymbol{p}} \end{bmatrix}, \tag{S35}$$

with $[\boldsymbol{F}_{\boldsymbol{x}} \mid \boldsymbol{F}_{\boldsymbol{p}}]$ the extended Jacobian of dimension $d \times (d + k)$ Then, introducing the normalized tangent (row) vector

$$\hat{\gamma} = (\hat{\gamma}^{(\boldsymbol{x})}, \hat{\gamma}^{(\boldsymbol{p})}) = \frac{(\dot{\boldsymbol{x}}, \dot{\boldsymbol{p}})}{\|(\dot{\boldsymbol{x}}, \dot{\boldsymbol{p}})\|_2}, \tag{S36}$$

we have $\hat{\gamma}\hat{\gamma}^T = 1$. Given a point $(\boldsymbol{x}^0, \boldsymbol{p}^0) := (\boldsymbol{x}(s_0), \boldsymbol{p}(s_0))$ on the curve of interest, the plane perpendicular to $\hat{\gamma}(s_0)$ in the $(\boldsymbol{x}, \boldsymbol{p})$ space a distance $s - s_0$ from $s_0$ is defined by

$$\hat{\gamma}(\boldsymbol{x} - \boldsymbol{x}^0,\ \boldsymbol{p} - \boldsymbol{p}^0)^T = s - s_0. \tag{S37}$$

On the curve of interest, $\boldsymbol{F}(\boldsymbol{x}^0, \boldsymbol{p}^0) = 0$. In addition, as the singularity we are interested in is a saddle-node bifurcation, the rank of $\boldsymbol{F}_{\boldsymbol{x}}(\boldsymbol{x}^0, \boldsymbol{p}^0)$ is $d - 1$ and with $\boldsymbol{F}_{\boldsymbol{p}}(\boldsymbol{x}^0, \boldsymbol{p}^0) \notin \text{range}(\boldsymbol{F}_{\boldsymbol{x}}(\boldsymbol{x}^0, \boldsymbol{p}^0))$. We can therefore find a nontrivial, unit vector $\boldsymbol{\varphi} \in \mathbb{R}^d$ such that $\boldsymbol{F}_{\boldsymbol{x}}(\boldsymbol{x}^0, \boldsymbol{p}^0)\boldsymbol{\varphi} = 0$ and $\boldsymbol{\varphi}^T\boldsymbol{\varphi} = 1$.

Given a distance $s$ from a known point on the curve, we therefore look for solutions of the system $H(\boldsymbol{y}, s) = 0$ where $\boldsymbol{y}$ is the augmented vector $\boldsymbol{y} := (\boldsymbol{x}, \boldsymbol{\varphi}, \boldsymbol{p}) \in \mathbb{R}^{2d+k}$ and

$$H(\boldsymbol{y}, s) = \begin{bmatrix} \boldsymbol{F}(\boldsymbol{x}, \boldsymbol{p}) \\ \boldsymbol{F}_{\boldsymbol{x}}(\boldsymbol{x}, \boldsymbol{p})\boldsymbol{\varphi} \\ \boldsymbol{\varphi}^T\boldsymbol{\varphi} - 1 \\ \hat{\gamma}^{(\boldsymbol{x})}(\boldsymbol{x} - \boldsymbol{x}^0)^T + \hat{\gamma}^{(\boldsymbol{p})}(\boldsymbol{p} - \boldsymbol{p}^0)^T - (s - s_0) \end{bmatrix}. \tag{S38}$$

We then have that

$$H_{\boldsymbol{y}}(\boldsymbol{y}, s) = \begin{bmatrix} \boldsymbol{F}_{\boldsymbol{x}} & \boldsymbol{0}_{d\times d} & \boldsymbol{F}_{\boldsymbol{p}} \\ \partial_{\boldsymbol{x}}[\boldsymbol{F}_{\boldsymbol{x}}\boldsymbol{\varphi}] & \boldsymbol{F}_{\boldsymbol{x}} & \partial_{\boldsymbol{p}}[\boldsymbol{F}_{\boldsymbol{x}}\boldsymbol{\varphi}] \\ \boldsymbol{0}_{1\times d} & 2\boldsymbol{\varphi} & \boldsymbol{0}_{1\times d} \\ \hat{\gamma}^{(\boldsymbol{x})} & \boldsymbol{0}_{1\times d} & \hat{\gamma}^{(\boldsymbol{p})} \end{bmatrix} \tag{S39}$$

and incrementing the arclength parameter $s$ by an amount $\Delta s$, we can proceed by Newton iterations to solve

$$H_{\boldsymbol{y}}(\boldsymbol{y}^0)\Delta\boldsymbol{y} = -H(\boldsymbol{y}^0) \tag{S40}$$

and trace the saddle-node bifurcation curve in the augmented space.

In our application, the components of $H$ and $H_{\boldsymbol{y}}$ are easily computed from a PLNN model using autodifferentiation. In particular, considering now the tilt vector $\boldsymbol{\tau}$ to be the governing parameter, we have

$$\begin{aligned} \boldsymbol{F}(\boldsymbol{x}, \boldsymbol{\tau}) &= -\nabla_{\boldsymbol{x}}\boldsymbol{\Phi}(\boldsymbol{x}) - \boldsymbol{\tau} \\ \boldsymbol{F}_{\boldsymbol{x}}(\boldsymbol{x}, \boldsymbol{\tau}) &= -\partial_{\boldsymbol{x}}[\nabla_{\boldsymbol{x}}\boldsymbol{\Phi}(\boldsymbol{x})] \end{aligned} \tag{S41}$$

and we see that $\boldsymbol{F}_{\boldsymbol{x}}$ does not depend on $\boldsymbol{\tau}$, so that the term $\partial_{\boldsymbol{\tau}}[\boldsymbol{F}_{\boldsymbol{x}}\boldsymbol{\varphi}]$ appearing in $H_{\boldsymbol{y}}$ is a zero matrix.

The python code implementing the arclength continuation algorithm can be found in the `plnn` package provided at https://github.com/AddisonHowe/plnn.

1545-7885. doi:10.1371/journal.pbio.2000640. URL https://journals.plos.org/plosbiology/article?id=10.1371/journal.pbio.2000640. Publisher: Public Library of Science.

[24] Patrick Kidger. *On Neural Differential Equations*. PhD thesis, University of Oxford, 2021.

[25] Martín Abadi, Ashish Agarwal, Paul Barham, Eugene Brevdo, Zhifeng Chen, Craig Citro, Greg S. Corrado, Andy Davis, Jeffrey Dean, Matthieu Devin, Sanjay Ghemawat, Ian Goodfellow, Andrew Harp, Geoffrey Irving, Michael Isard, Yangqing Jia, Rafal Jozefowicz, Lukasz Kaiser, Manjunath Kudlur, Josh Levenberg, Dandelion Mané, Rajat Monga, Sherry Moore, Derek Murray, Chris Olah, Mike Schuster, Jonathon Shlens, Benoit Steiner, Ilya Sutskever, Kunal Talwar, Paul Tucker, Vincent Vanhoucke, Vijay Vasudevan, Fernanda Viégas, Oriol Vinyals, Pete Warden, Martin Wattenberg, Martin Wicke, Yuan Yu, and Xiaoqiang Zheng. TensorFlow: Large-scale machine learning on heterogeneous systems, 2015. URL https://www.tensorflow.org/. Software available from tensorflow.org.

[26] Jason Ansel, Edward Yang, Horace He, Natalia Gimelshein, Animesh Jain, Michael Voznesensky, Bin Bao, Peter Bell, David Berard, Evgeni Burovski, Geeta Chauhan, Anjali Chourdia, Will Constable, Alban Desmaison, Zachary DeVito, Elias Ellison, Will Feng, Jiong Gong, Michael Gschwind, Brian Hirsh, Sherlock Huang, Kshiteej Kalambarkar, Laurent Kirsch, Michael Lazos, Mario Lezcano, Yanbo Liang, Jason Liang, Yinghai Lu, CK Luk, Bert Maher, Yunjie Pan, Christian Puhrsch, Matthias Reso, Mark Saroufim, Marcos Yukio Siraichi, Helen Suk, Michael Suo, Phil Tillet, Eikan Wang, Xiaodong Wang, William Wen, Shunting Zhang, Xu Zhao, Keren Zhou, Richard Zou, Ajit Mathews, Gregory Chanan, Peng Wu, and Soumith Chintala. PyTorch 2: Faster Machine Learning Through Dynamic Python Bytecode Transformation and Graph Compilation. In *29th ACM International Conference on Architectural Support for Programming Languages and Operating Systems, Volume 2 (ASPLOS '24)*. ACM, April 2024. doi:10.1145/3620665.3640366. URL https://pytorch.org/assets/pytorch2-2.pdf.

[27] Patrick Kidger and Cristian Garcia. Equinox: neural networks in JAX via callable PyTrees and filtered transformations. *Differentiable Programming workshop at Neural Information Processing Systems 2021*, 2021.

[28] DeepMind, Igor Babuschkin, Kate Baumli, Alison Bell, Surya Bhupatiraju, Jake Bruce, Peter Buchlovsky, David Budden, Trevor Cai, Aidan Clark, Ivo Danihelka, Antoine Dedieu, Claudio Fantacci, Jonathan Godwin, Chris Jones, Ross Hemsley, Tom Hennigan, Matteo Hessel, Shaobo Hou, Steven Kapturowski, Thomas Keck, Iurii Kemaev, Michael King, Markus Kunesch, Lena Martens, Hamza Merzic, Vladimir Mikulik, Tamara Norman, George Papamakarios, John Quan, Roman Ring, Francisco Ruiz, Alvaro Sanchez, Laurent Sartran, Rosalia Schneider, Eren Sezener, Stephen Spencer, Srivatsan Srinivasan, Miloš Stanojević, Wojciech Stokowiec, Luyu Wang, Guangyao Zhou, and Fabio Viola. The DeepMind JAX Ecosystem, 2020. URL http://github.com/google-deepmind.

[37] E. Hairer, S.P. Nørsett, and G. Wanner. *Solving Ordinary Differential Equations I Nonstiff Problems*. Springer, Berlin, second revised edition edition, 2008.

[38] A. Jepson and A. Spence. Folds in Solutions of Two Parameter Systems and Their Calculation. Part I. *SIAM Journal on Numerical Analysis*, 22(2):347–368, April 1985. ISSN 0036-1429, 1095-7170. doi:10.1137/0722021. URL http://epubs.siam.org/doi/10.1137/0722021.

[39] Willy J. F. Govaerts. *Numerical Methods for Bifurcations of Dynamical Equilibria*. Society for Industrial and Applied Mathematics, 2000. doi:10.1137/1.9780898719543. URL https://epubs.siam.org/doi/abs/10.1137/1.9780898719543.

[40] Rüdiger Seydel. *Practical Bifurcation and Stability Analysis*, volume 5 of *Interdisciplinary Applied Mathematics*. Springer New York, New York, NY, 2010. ISBN 978-1-4419-1739-3 978-1-4419-1740-9. doi:10.1007/978-1-4419-1740-9. URL https://link.springer.com/10.1007/978-1-4419-1740-9.

[41] Alastair Spence and Ivan G. Graham. Numerical Methods for Bifurcation Problems. In Mark Ainsworth, Jeremy Levesley, and Marco Marletta, editors, *The Graduate Student's Guide to Numerical Analysis '98: Lecture Notes from the VIII EPSRC Summer School in Numerical Analysis*, pages 177–216. Springer, Berlin, Heidelberg, 1999. ISBN 978-3-662-03972-4. doi:10.1007/978-3-662-03972-4_5. URL https://doi.org/10.1007/978-3-662-03972-4_5.
